# metaDMG – A Fast and Accurate Ancient DNA Damage Toolkit for Metagenomic Data

**DOI:** 10.1101/2022.12.06.519264

**Authors:** Christian Michelsen, Mikkel Winther Pedersen, Antonio Fernandez-Guerra, Lei Zhao, Troels C. Petersen, Thorfinn Sand Korneliussen

**Affiliations:** Niels Bohr Institute, University of Copenhagen, Blegdamsvej 17, 2100 Copenhagen, Denmark; GLOBE, Section for Geogenetics, Øster Voldgade 5-7, 1350, Copenhagen, Denmark

**Keywords:** ancient DNA, DNA damage estimation, DNA damage, metaDMG, metagenomics

## Abstract

1. **Motivation** Under favourable conditions DNA molecules can persist for hundreds of thousands of years. Such genetic remains make up invaluable resources to study past assemblages, populations, and even the evolution of species. However, DNA is subject to degradation, and hence over time decrease to ultra low concentrations which makes it highly prone to contamination by modern sources. Strict precautions are therefore necessary to ensure that DNA from modern sources does not appear in the final data is authenticated as ancient. The most generally accepted and widely applied authenticity for ancient DNA studies is to test for elevated deaminated cytosine residues towards the termini of the molecules: DNA damage. To date, this has primarily been used for single organisms and recently for read assemblies, however, these methods are not applicable for estimating DNA damage for ancient metagenomes with tens and even hundreds of thousands of species.
2. **Methods** We present `metaDMG`, a novel framework and toolkit that allows for the estimation, quantification and visualization of postmortem damage for single reads, single genomes and even metagenomic environmental DNA by utilizing the taxonomic branching structure. It bypasses any need for initial classification, splitting reads by individual organisms, and realignment. We have implemented a Bayesian approach that combines a modified geometric damage profile with a beta-binomial model to fit the entire model to the individual misincorporations at all taxonomic levels.
3. **Results** We evaluated the performance using both simulated and published environmental DNA datasets and compared to existing methods when relevant. We find `metaDMG` to be an order of magnitude faster than previous methods and more accurate – even for complex metagenomes. Our simulations show that `metaDMG` can estimate DNA damage at taxonomic levels down to 100 reads, that the estimated uncertainties decrease with increased number of reads and that the estimates are more significant with increased number of C to T misincorporations.
4. **Conclusion** `metaDMG` is a state-of-the-art program for aDNA damage estimation and allows for the computation of nucleotide misincorporation, GC-content, and DNA fragmentation for both simple and complex ancient genomic datasets, making it a complete package for ancient DNA damage authentication.

## 1 INTRODUCTION

Throughout the life of an organism it contaminates its environment with DNA, cells, or tissue, thus leaving genetic traces behind. As the cell leaves its host, DNA repair mechanisms stops and the DNA is subjected to intra and extra cellular enzymatic, chemical, and mechanical degradation, resulting in fragmentation and molecular alterations that over time lead to the characteristics of ancient DNA (Briggs et al., 2007; Dabney, Meyer, and Pääbo, 2013). Ancient DNA (aDNA) has been shown to persist in a diverse variety of environmental contexts, e.g. within fossils such as bones, soft-tissue, and hair, as well as in geological sediments, archaeological layers, ice-cores, permafrost soil for hundreds of thousands of years (Valk et al., 2021; Zavala et al., 2021). Common for all is that they have an accumulated amount of deaminated cytosines towards the termini of the DNA strand, which, when amplified, results in misincorporations of thymines on the cytocines (Ginolhac et al., 2011; Dabney, Meyer, and Pääbo, 2013).

Even though postmortem DNA damage (PMD) is characterized by the four Briggs parameters (Briggs et al., 2007), they are rarely used directly for asserting “ancientness”. Researchers working with ancient DNA tend to simply use the empirical C→T on the first position of the fragment together with other supporting summary statistic of the experiment (Jónsson et al., 2013). Quantifying PMD have become standard for single individual sources like hair, bones, teeth and also applied to smaller subsets of species in ancient environmental metagenomes (Pedersen et al., 2016; Murchie et al., 2021; Wang, Pedersen, et al., 2021; Zavala et al., 2021). While this is a relatively fast process for single individuals it becomes increasingly demanding, iterative, and time consuming as the samples and the diversity within increases, as in the case for metagenomes from ancient soil, sediments, dental calculus, coprolites, and other ancient environmental sources. It has therefore been practice to estimate damage for only the key taxa of interest in a metagenome, as metagenomic samples easily include tens of thousands of different taxonomic entities, which makes a complete estimate across the metagenomes computationally intractable, if not an impossible task (Pedersen et al., 2016). To overcome these limitations, we designed a toolkit called `metaDMG` (pronounced metadamage) which allows for the rapid computation of various statistics relevant for the quantification of PMD at read level, single genome level, and even metagenomic level by taking into account the intricate branching structure of the taxonomy of the possible multiple alignments for the single reads.

Our thorough analysis with both simulated and real data shows that `metaDMG` is both faster at ancient DNA damage estimation and provides more accurate damage estimates. Furthermore, as `metaDMG` is designed with the increasingly large datasets that are currently generated in the field of ancient environmental DNA in mind, `metaDMG` is able to process complex metagenomes within hours instead of days. At the same time, it outperforms standard tools that estimate DNA damage for single genomes and samples with low complexity. Furthermore, it can compute a global damage estimate for a metagenome as a whole. Lastly, `metaDMG` is compatible with the NCBI taxonomy and use ngsLCA (Wang, T. S. Korneliussen, et al., 2022) to perform a lowest common ancestor (LCA) classification of the aligned reads to get precise damage estimates at all taxonomic levels. It also allows for custom taxonomies and thus also the use of metagenomic assembled genomes (MAGs) as references.

This paper is organized as follows. In ***section 2*** we present our statistical models including two novel test statistics, *D*_fit_ and *Z*_fit_. We quantify the performance of our test statistics using various simulation approaches in ***section 3***. The results of these simulations is shown in ***section 4*** and finally, the method and results are discussed in ***section 5***.

## 2 METHODS & MATERIALS

To quantify ancient damage, one can either compute it on a per read level or across an entire taxa. A priori, the actual biochemical changes which characterizes post mortem damage in a single read cannot be directly observed, but by aligning each fragment and considering the observed difference between the reference and read, the possible PMD can be computed. We have (re)implemented the approach used in PMDtools (Skoglund et al., 2014) which allows for the extraction of single DNA reads which are estimated to contain PMD, see ***Appendix 1***. This approach, will preferentially choose reads that has excess of C→T in the first positions and can not be used directly for asserting or quantifying to what degree a given library might contain damaged fragments. We have therefore developed a novel statistical method that aims to mitigate this caveat by using all reads or reads that aligns to specific taxa. First we will define the mismatch matrices in ***subsection 2.1*** followed by the lowest common ancestor method in ***subsection 2.2***. The mismatch matrices can further be improved by multinomial regression, see ***subsection 2.3***, however, this requires more data than than what is usually available in metagenomic studies. As such, we present the beta-binomial damage model in ***subsection 2.4*** which aims to work even on extremely low-coverage data.

### 2.1 Mismatch matrices/nucleotide misincorporation patterns

We seek to obtain the pattern or signal of damage across multiple reads by generating what is called the mismatch matrix or the nucleotide misincorporation matrix. This matrix represents the nucleotide substitution counts across reads and provides us with the position dependent mismatch matrices, *M*(*x*), with *x* denoting the position in the read, starting from 1. At a specific position *x*, *M*_ref→obs_(*x*) describes the number of nucleotides that was mapped to a reference base 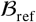 but was observed to be 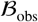, where 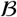 is one of the four bases: A, C, G, T. The number of C→T transitions at the first position, e.g., is denoted as *M*_*C*→*T*_ (*x* = 1).

Alignments for a read can be discarded based on their mapping quality, and we also give the user the possibility of filtering out specific nucleotides of the read if the base quality score fall below some threshold. The quality scores could also be used as probabilistic weights, however, due to the four-bin discretization of quality scores on modern day sequencing machines, we limit the use of these to filtering.

### 2.2 Lowest Common Ancestor and Mismatch matrices

For environmental DNA (eDNA) studies a competitive alignment approach is routinely applied. Here all possible alignments for a given read are considered. Each read is mapped against a multi species reference databases (e.g. nucleotide or RefSeq from NCBI or custom downloaded). A single read might map to a highly conserved gene that is shared across higher taxonomic ranks such as class or even domains. This read will not provide relevant information due to the generality, whereas a read that maps solely to a single species, e.g, would be indicative of the read being well classified. We limit the tabulation and construction of the mismatch matrices to the subset of reads that are well classified.

For each read, we compute the lowest common ancestor using all alignments contained within the user defined taxanomic threshold (species, genus or family) and tabulate the mismatches matrices for each cycle (Wang, T. S. Korneliussen, et al., 2022). If none of the alignments pass the filtering thresholds (excess similarity, mapping quality, etc.), the read is discarded. Depending on the run mode, we allow for the construction of these mismatch matrices on three different levels. Firstly, we can obtain a basic single global mismatch matrix which could be relevant in a standard single genome aDNA study and similar to the tabulation used in mapDamage (Jónsson et al., 2013). Secondly, we can obtain the per reference counts, or, finally, if a taxonomy database has been supplied, we can build mismatch matrices at the species level and aggregate from leaf nodes to the internal taxonomic ranks (genus, kingdom etc) towards the root. We will use the term “taxa” to refer to either of these levels; i.e. a specific taxa can either refer to a specific LCA, a specific reference, or all reads in a global estimate, depending on the run-mode.

When aggregating the mismatch matrices for the internal nodes in our taxonomic tree, two different approaches can be taken. Either all alignments of the read will be counted, which we will refer to as weight-type 0, or the counts will be normalized by the number of alignments of each read; weight-type 1, which is the default.

### 2.3 Regression Framework

The nucleotide misincorporation frequencies are routinely used as the basis for assessing whether or not a given library is ancient by looking at the expected drop of C→T (or its complementary G→A) frequencies as a function of the position of the reads. This signal is caused by a higher deamination rate in the single-strand part of the damaged fragment than that in the double strand part. The mismatch matrix is constructed based on the empirical observations and are subject to stochastic noise. The effect of noise in the mismatch matrix can be limited by the use of the multinomial regression model. We continue the work of Cabanski et al., 2012 to provide four different regression methods to stabilize the raw mismatch matrix across all combinations of reference bases, observed bases, strands and positions, see ***Appendix 2*** for details, derivation and results. Given enough sequencing data, this approach will provide an improved, noise-reduced mismatch matrix which would be relevant for single genome ancient DNA studies. However, for extremely low coverage studies, such as environmental DNA, the method is likely to overfit and would not be as suitable as the simplified model described in the ***subsection 2.4***.

### 2.4 Damage Estimation

In standard ancient DNA context it is generally not possible to obtain vast amounts of data and thus we propose two novel tests statistics, *D*_fit_ and *Z*_fit_, that are especially suited for this common scenario. The damage pattern observed in aDNA has several features which are well characterized. By modelling these, one can construct observables sensitive to aDNA signal. We model the damage patterns seen in ancient DNA by looking exclusively at the C→T transitions in the forward direction (5’) and the G→A transitions in the reverse direction (3’). For each taxa, we denote the number of transitions, *k*(*x*), as:

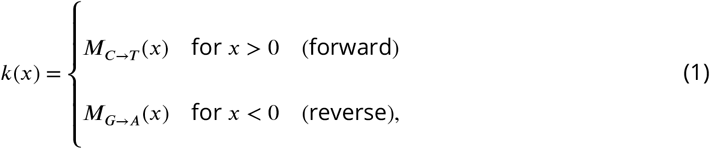

and the number of reference counts *N*(*x*):

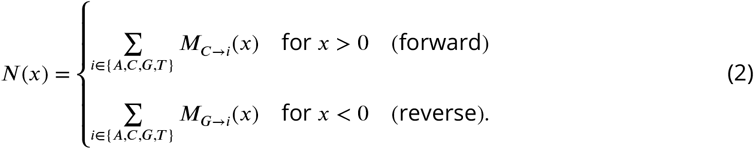

The damage frequency is thus *f* (*x*) = *k*(*x*)∕*N*(*x*).

A natural choice of likelihood model would be the binomial distribution. However, we found that a binomial likelihood lacks the flexibility needed to deal with the large amount of variance (overdispersion) we found in the data due to pooly curated references and possible misalignments.

To accommodate overdispersion, we instead apply a beta-binomial distribution, 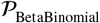, which treats the probability of deamination, *p*, as a random variable following a beta distribution^1^ with mean *μ* and concentration *ϕ*: *p* ~ Beta(*μ*, *ϕ*). The beta-binomial distribution has the the following probability density function:

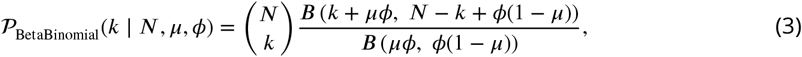

where *B* is defined as the beta function:

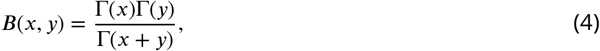

with Γ(⋅) being the gamma function (Cepeda-Cuervo and Cifuentes-Amado, 2017).

The close resemblance to a binomial model is most easily seen by comparing the mean and variance of a random variable *k* following a beta-binomial distribution, 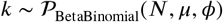:

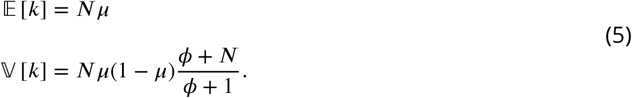

The expected value of *k* is similar to that of a binomial distribution and the variance of the beta-binomial distribution reduces to a binomial distribution as *ϕ* → ∞. The beta-binomial distribution can thus be seen as a generalization of the binomial distribution.

Note that both equation (3) and (5) relates to the damage at a specific base position (cycle), i.e. for a single *k* and *N*. To estimate the overall damage in the entire read using the position dependent counts, *k*(*x*) and *N*(*x*), we model *μ* as being position dependent, *μ*(*x*), and assume a position-independent concentration, *ϕ*. We model the damage frequency with a modified geometric sequence, i.e. exponentially decreasing for discrete values of *x*:

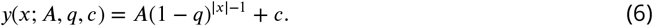

Here *A* is the amplitude of the damage and *q* is the relative decrease of damage pr. position. A background, *c*, was added to reflect the fact that the mismatch between the read and reference might be due to other factors than just ancient damage. As such, we allow for a non-zero amount of damage, even as *x* → ∞. This is visualized in ***Figure 1*** along with a comparison between the classical binomial model and the beta-binomial model.

**Figure 1.**
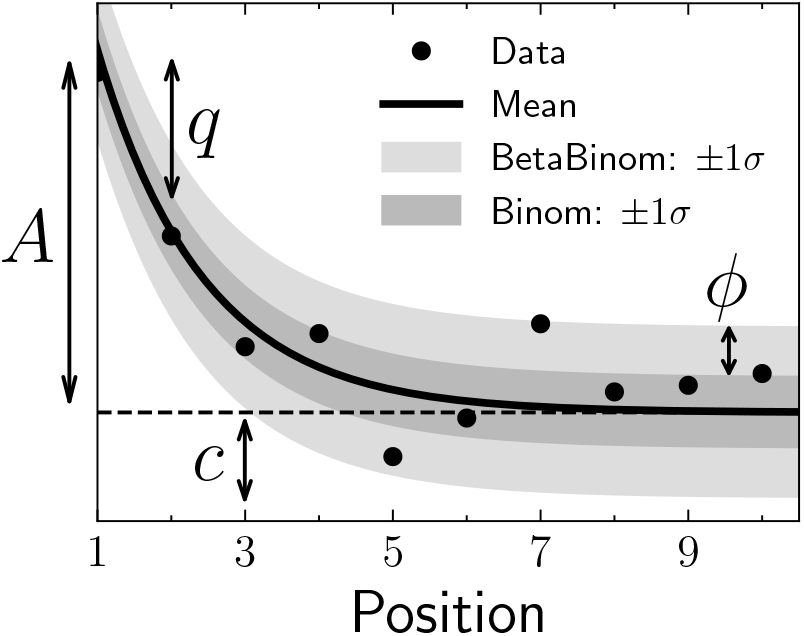
Illustration of the damage model. The figure shows data points as circles and the damage, *f* (*x*), as a solid line. The amplitude of the damage is *A*, the offset is *c*, and the relative decrease in damage pr. position is given by *q*. The damage uncertainty for a binomial model is shown in dark grey and the uncertainty for a beta-binomial model in light grey.

To estimate the four fit parameters, *A*, *q*, *c*, and *ϕ*, we apply Bayesian inference to utilize domain specific knowledge in the form of priors. We assume weakly informative beta-priors^2^ for both *A*, *q*, and *c*. In addition to this, we assume an exponential prior on *ϕ* with the requirement of *ϕ* > 2 to avoid too much focus on 0-or-1 probabilities (McElreath, 2020). The final model is thus:

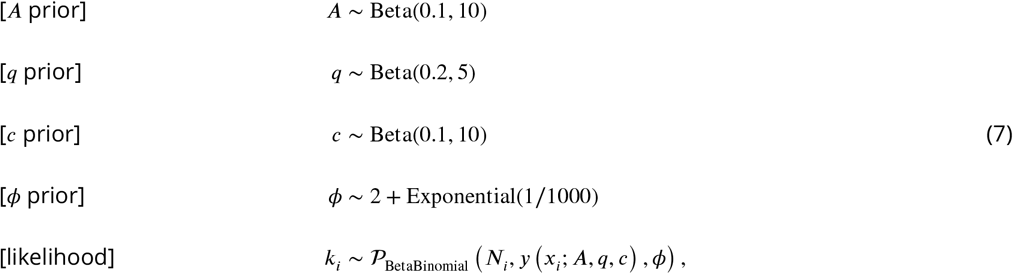

where *i* is an index running over all positions.

We define the damage due to deamination, 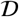, as the background-subtracted damage frequency at the first position: 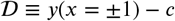. As such, 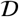 is the damage related to ancientness. Using the properties of the beta-binomial distribution, eq. (5), we find the mean and variance of 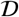:

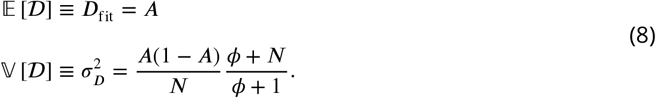

Since 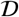 estimates the overexpression of damage due to ancientness, not only is the mean of 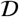, *D*_fit_, relevant but also the certainty of it being non-zero (and positive). We quantify this through the significance *Z*_fit_ = *D*_fit_ ∕*σ*_*D*_ which is thus the number of standard deviations (“sigmas”) away from zero. Assuming a Gaussian distribution of 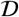, *Z*_fit_ > 2 would indicate a probability of *D* being larger than zero, i.e. containing ancient damage, with more than 97.7% probability. This assumption works well in the case of many reads or a high amount of damage due to central limit theorem. When the assumption breaks down, the significance is still a relevant test statistic, it is only the conversion to a probability that will become biased.

These two values allows us to not only quantify the amount of ancient damage (*D*_fit_) but also the certainty of this damage (*Z*_fit_) without having to run multiple models and comparing these. An intuitive interpretation of our *D*_fit_ statistic is, that this is the excess deamination in the beginning of the read, taking all cycle positions into account and excluding the constant deamination background (*c*). This is visually similar to the *A* parameter in ***Figure 1***.

We perform the Bayesian inference of the parameters models using Hamiltonian Monte Carlo (HMC) sampling which is a particular of Monte Carlo Markov Chain (MCMC) algorithm (Betancourt, 2018). Specifically, we use the NUTS implementation in NumPyro (Phan, Pradhan, and Jankowiak, 2019), a Python package which uses JAX (Bradbury et al., 2018) under the hood for automatic differentiation and JIT compilation. We treat each taxa as being independent and generate 1000 MCMC samples after an initial 500 samples as warm up.

Since running the full Bayesian model is computationally expensive, we also allow for a faster, approximate method by fitting the maximum a posteriori probability (MAP) estimate. We use iMinuit (Dembinski et al., 2021) for the MAP optimization with Numba acceleration (Lam, Pitrou, and Seibert, 2015) for even faster run times. On a Macbook M1 Pro model from 2021, the timings for running the full Bayesian model is 1.41 ± 0.04 s pr. fit and for the MAP it is 4.34 ± 0.07 ms pr. fit, showing more than a 2 order increase in performance (around 300x) for the approximate model. Both models allow for easy parallelisation to decrease the computation time.

### 2.5 Visualisation

We provide an interactive graphical user interface (dashboard) to visualise, explore, and manipulate the results from the modelling phase. An interactive example of this can be found online (https://metadmg.onrender.com/). The structure of the dashboard is explained in ***Figure 2***. The dashboard allows for filtering, styling and variable selection, visualizing the mismatch matrix related to a specific taxa, and exporting of both fit results and plots. By filtering, we include both filtering by sample, by the summary statistics of the data (e.g. requiring *D*_fit_ to be above a certain threshold), and even by taxonomic level (e.g. only looking at taxa that are part of the Mammalia class). We greatly believe that a visual overview of the fit results increase understanding of the data at hand. The dashboard is implemented with Plotly plots and incorporated into a Dash dashboard (Plotly, 2015).

**Figure 2.**
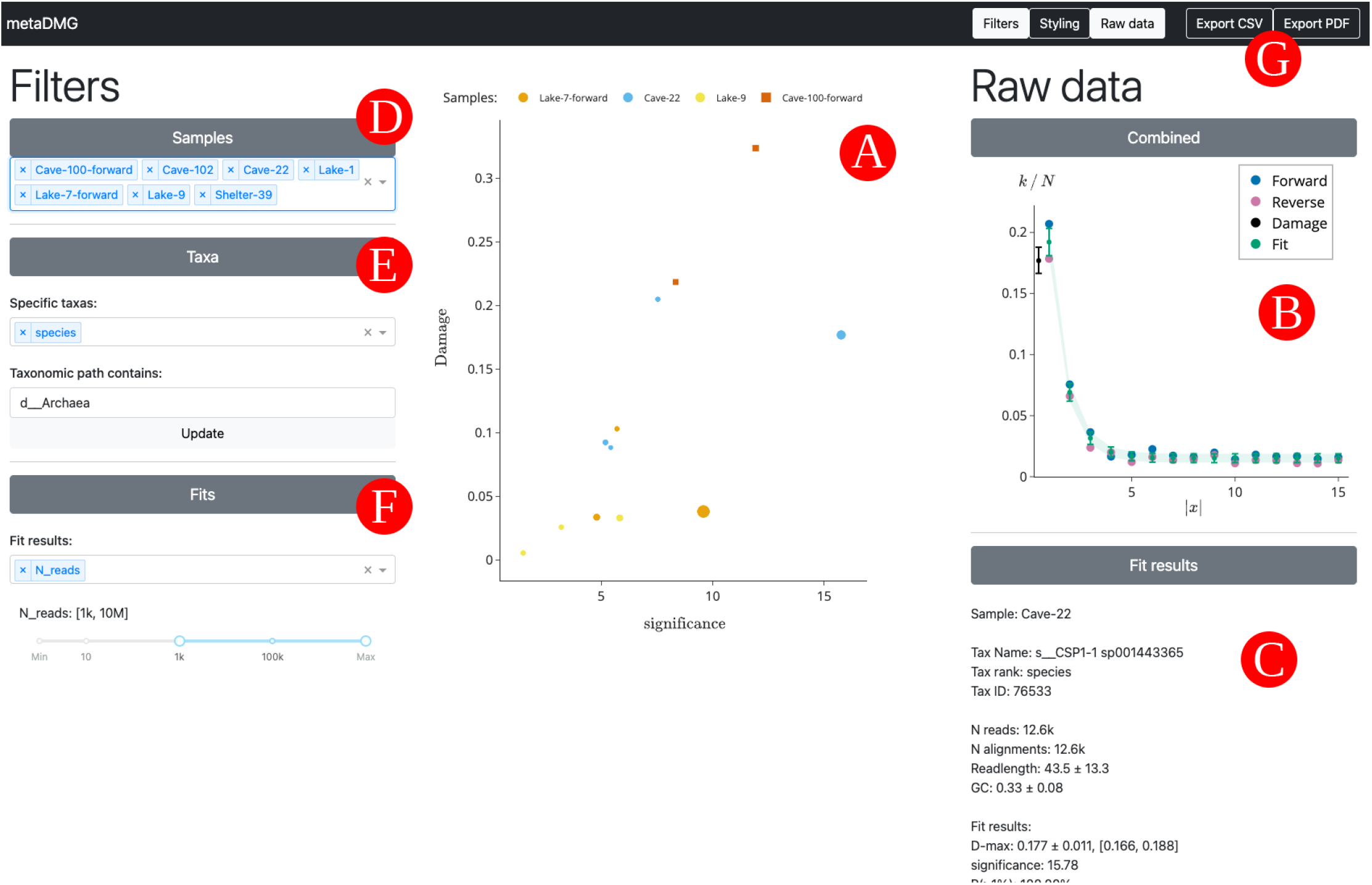
Overview of the interactive `metaDMG` dashboard. A) The main damage plot shows the damage (*D*_fit_) on the y-axis and the significance (*Z*_fit_) on the x-axis. Each point is a single taxa from one of the metagenomic samples, see ***Table 1***. Once clicked on a specific taxa, the right-hand window shows information about the selected taxa and related fit. B) The top window shows a plot of the damage frequency for both the forward and reverse direction along with the estimated fit and damage. C) Below, the results of the fit are shown, including taxonomic information, read-specific information, the fit results, and the full taxonomic path. D) In the left filtering window, the samples to include can be selected. E) This windows allows for selection based on taxa-specific criteria. Here we show a selection of only taxa with “species” as their LCA and taxa that are part of the archaea domain. F) The final filtering window allows for setting fit related thresholds such as the minimum damage or significance. Here it is shown discarding taxa with fever than 1000 reads. G) In the top right, after the selection and filtering process is finished, the final taxa can be exported to a CSV file along with all of the fit information, or the damage plots can be generated and saved.

## 3 SIMULATION STUDY

To determine `metaDMG’s` performance, we performed a set of rigorous in-silica simulations to identify and quantify any possible biases as well the accuracy of our test statistics. Overall, the simulations can be split two groups. The first is based on a genome from a single species and is used to measure the performance of the actual damage estimation and damage model. The second is based on syntethic ancient metagenomic datasets using the statistics and nature of a set of published ancient metagenomes.

### 3.1 Single-genome simulations

The first simulations follow a simple setup in which we extract reads from a set of representative genomes having variable length and GC-content. We next added post-mortem damage misincorporations using NGSNGS (Henriksen, Zhao, and T. Korneliussen, 2022) a recent implementation of the original Briggs model similar to Gargammel (Neukamm, Peltzer, and Nieselt, 2021) and lastly added sequencing errors (Renaud et al., 2017). All reads are hereafter mapped using Bowtie2 against each of the respective reference genomes and ancient DNA damage estimated the DNA damage using `metaDMG`. The simulations were computed with varying amount of damage added by changing the single-stranded DNA deamination, *δ*_SS_ in the original Briggs model (Briggs et al., 2007).

In detail, we focused on the following genomes; Homo Sapiens mitochondrial^3^, a Betula nana chloroplast^4^, and three microbial genomes (Fusobacterium pseudoperiodonticum^5^, Neisseria cinerea^6^, and Actinomyces oris strain S64C^7^) with the varying GC-content, low (28%), medium (37%), and high (50%) respectively. For each simulation, we performed 100 independent replications to measure the variability of the parameter estimation and quantify the robustness of the estimates. We further simulated eight different sets of damage (0%, 1%, 2%, 5%, 10%, 15%, 20%, and 30% damage on position 1), all with 13 sets of different number of reads (10, 25, 50, 100, 250, 500, 1.000, 2.500, 5.000, 10.000, 25.000, 50.000, and 100.000 reads). We also sought to measure the effect of the fragment lengths using three sets of different fragment length distributions sampled from a *lognormal* distribution with mean 35, 60, and 90, each with a standard deviation of 10). Furthermore, to investigate whether the damage estimation by `metaDMG` is independent of contig size, we artificially created three different genomes by sampling 1.000, 10.000 or 100.000 different basepairs from a uniform categorical distribution of A, C, G, and T. Based on these three genomes, we added artificial deamination for a different number of reads, as for the other simulations. Lastly, we also created 1000 repetitions of non-damaged simulations for Homo Sapiens to measure the rate of false positives. The exact commands used can be found in ***Appendix 3***.

To compare the damage estimates to known values, for each of the genomes mentioned above and for each amount of artificial damage, we generated 1.000.000 reads using NGSNGS without any added sequencing noise. The values we compare is the difference in damage frequency at position 1 and 15:

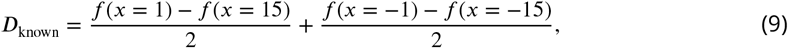

which is the average of the C→T damage frequency difference and the G→A damage frequency difference.

### 3.2 Metagenomic Simulations

A metagenome contains a complex mixture of organisms, all with highly different characteristics in GC content, read length, abundance, or degree of DNA damage, and there are large differences between different environments. It is therefore far from simple to obtain DNA damage estimates for such multitude of organisms. In order to test the accuracy and sensitivity of `metaDMG`, we simulated six of the nine ancient metagenomes (with more than 1 million reads) covering a wide span of environments and ages (***Table 1***).

**Table 1.** Metagenomic samples. “Name” is the name of the sample used throughout this paper. “Site” is the type of metagenomic site. “Type” is the type of environment. “Age” is the approximate age of the sample in kyr Bp. “Sediment” is the name type of sediment. “Instrument” is the Illumina model. “Library” is the library type where D. means double stranded and S. means single stranded. “Reads” is the raw number of reads (in millions). “Source” is the source of the data. The dagger (†) indicates samples that were not a part of the metagenomic simulation pipeline.

| Name | Site | Type | Age (kyr) | Sediment | Instrument | Library | Reads (M) | Source |
| --- | --- | --- | --- | --- | --- | --- | --- | --- |
| Library-0<sup>†</sup> | Control | Control | 0 | Reagents | HiSeq4000 | D. | 19.7 | (Ardelean et al., 2020) |
| Pitch-6 | Syltholmen pitch | Chewed organic material | 5.7 | Organic material | HiSeq2500 | D. | 150.3 | (Jensen et al., 2019) |
| Lake-1<sup>†</sup> | Spring Lake | Lake gyttja/sediment | 1.4 | Organic material | HiSeq 100 | D. | 49.8 | (Pedersen et al., 2016) |
| Lake-7 | Lake CH12 | Lake gyttja/sediment | 6.7 | Organic material | HiSeq2500 | S. | 291.9 | (Schulte et al., 2021) |
| Lake-9 | Spring Lake | Lake gyttja/sediment | 9.2 | Organic material | HiSeq 100 | D. | 128.4 | (Pedersen et al., 2016) |
| Shelter-39<sup>†</sup> | Abri Pataud | Rock shelter | 39.4 | Sediment | MiSeq | S. | 0.4 | (Braadbaart et al., 2020) |
| Cave-22 | Chiquihuite cave | Cave sediment | 22.2 | Carbonate rock | HiSeq4000 | D. | 5.7 | (Ardelean et al., 2020) |
| Cave-100 | Eustatuas Cave | Cave sediment | 100 | Carbonate rock | HiSeq2500 | S. | 21.8 | (Vernot et al., 2021) |
| Cave-102 | Pesturina Cave | Neanderthal tooth | 102 | Dental calculus | HiSeq4000 | D. | 12.3 | (Fellows Yates et al., 2021) |

First, we mapped all reads of each metagenome with bowtie2 against a database consisting of the GTDB (r202) (Parks et al., 2018) species cluster reference sequences, all organelles from NCBI RefSeq (NCBI Resource Coordinators, 2018), and the reference sequences from CheckV (Nayfach et al., 2021). We then used bam-filter v1.0.11 (Fernandez-Guerra, 2022a) with the `flag --read-length-freqs` to get read length distributions for each genome reads aligned to and their respective abundance. Next, we filtered genomes with an observed-to-expected coverage ratio greater than 0.75 using bamfilter. The filtered BAM files were then processed by `metaDMG` to obtain misincorporation matrices for each genome. The abundance tables, fragment length distribution, and misincorporation matrices were then used in aMGSIM-smk v0.0.1 (Fernandez-Guerra, 2022b), a Snakemake workflow (Mölder et al., 2021) that facilitates the generation of multiple synthetic ancient metagenomes. The underlying tools in this workflow is the gargammel toolkit (Renaud et al., 2017), that based on input read length distribution extract a subset of sequences (FragSim) with similar length. This is then followed by the addition of *C* → *T* substitutions (DeamSim) which mimics the postmortem damage process. Finally the deaminated sequences are passed to the ART (Huang et al., 2012) for sequence simulation. The data used and generated by the workflow can be obtained from [ERDA](https://sid.erda.dk/cgi-sid/ls.py?share_id=l7NGWfSkXq). We then performed taxonomic profiling and damage estimation using identical parameters as for the synthetic reads generated by aMGSIM-smk.

## 4 RESULTS

We tested the accuracy and performance of the `metaDMG` damage estimates, *D*_fit_, using a set of different simulation scenarios and subsequently tested on 9 real-life ancient metagenomic dataset.

### 4.1 Single-genome simulation results

To illustrate the results the performance on single-genomes, we first focus on a single, specific set of simulation parameters. This simulation is based on the Homo Sapiens genome with the Briggs parameter *δ*_SS_ = 0.31 (approximately 10% damage) and a mean fragment length of 60. In general, we use *δ* = 0.0097, *ν* = 0.024, and *λ* = 0.36 as Briggs parameters, while varying *δ*_SS_ (Briggs et al., 2007). We show the `metaDMG` damage results for the 100 independent replications in ***Figure 3***. The left part of the figure shows the individual `metaDMG` damage estimates for an arbitrary choice of 20 replications (iteration 60 to 79). When the damage estimates are very low, the distribution of *D*_fit_ is skewed (restricted to positive values), sometimes leading to errorbars going into negative damage, which represents unrealistic estimates. The right hand side of the figure visualizes the average amount of damage based on all 100 replications across a varying number of reads. This shows that the damage estimates converge to the known value with more data, and that one needs more than 100 reads to even get strictly positive damage estimates (when including uncertainties) for this specific set of simulation parameters.

**Figure 3.**
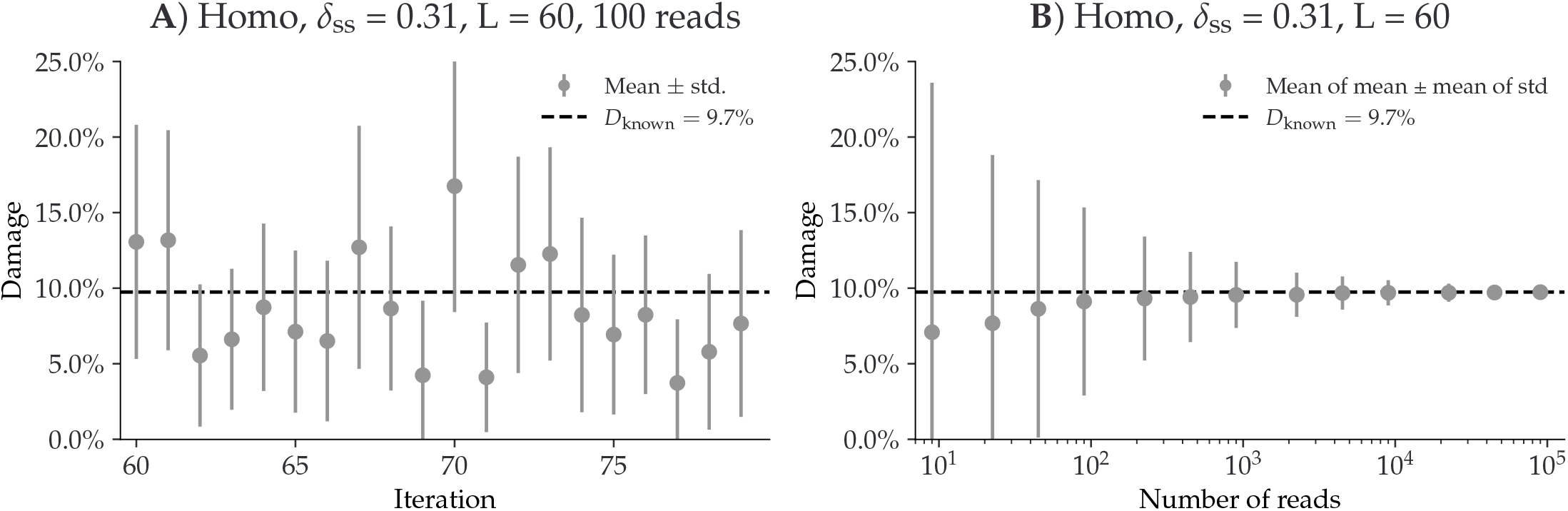
Overview of the single-genome simulations based on the Homo Sapiens genome with a fragment length distribution with mean 60 and the Briggs parameter *δ*_SS_ = 0.31 (approximately 10% damage). **A**) This plot shows the estimated damage (*D*_fit_) of 20 replicates, each with 100 reads. The grey points shows the mean damage (with its standard deviation as errorbars). The known damage (*D*_known_) is shown as a dashed line, see eq. (9). **B**) This plot shows the average damage as a function of the number of reads. The grey points show the average of the individual means (with the average of the standard deviations as errors).

Across multiple simulations, each with 8 different damage levels, 13 different numbers of reads, and 100 replications, we find no significant difference in test statistic across different species (***Figure S5*** and ***Figure S6***), across different GC-levels (***Figure S7***–***Figure S9***), different fragment length distributions (***Figure S10***-***Figure S12***), or even different contig lengths (***Figure S13***–***Figure S15***), see ***Appendix 4***. Based on the single-genome simulations, we compute the relationship between the amount of damage in a taxa and the number of reads required to correctly infer that the reads from that taxa are damaged, see ***Figure 4***. If we want to assert damage with a significance of more than 2 (solid blue line) in a sample with around 5% expected damage, it requires about 1000 reads to be 95% certain that we will find results this good, whereas we only need 100 reads if our target organism has 30 % damage.

**Figure 4.**
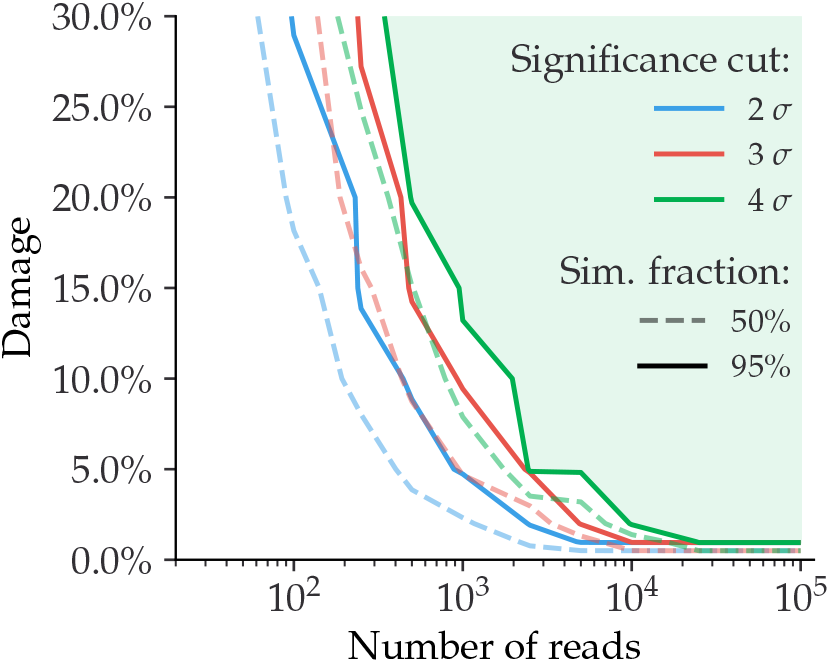
Relationship between the damage and the number of reads for simulated data (single-genome). Given a specific significance cut, the solid contour line shows the relationship between the amount of damage and the number of reads required to be able to correctly infer damage in 95% of the taxa. The dashed line shows the similar value for a simulation fraction of 50%. The green part of the figure shows the “good” region of number of reads and estimated damage, given than one wants to be more than 95% certain of correctly identifying damage with more than 4*σ* confidence.

Finally, to quantify the risk of incorrectly classifying a non-ancient taxa as damaged, we created 1000 independent replications for a varying number of reads, where none of them had any artificial ancient damage applied, only sequencing noise. ***Figure 5*** shows the damage (*D*_fit_) as a function of the significance (*Z*_fit_) for the case of 1000 reads. Even though the estimated damage is larger than zero, the damage is non-significant since the significance is less than one. When looking at all the figures across the different number of reads, see ***Appendix 5***, we note that a relaxed significance threshold requiring that *D*_fit_ > 1% and *Z*_fit_ > 2 would filter out all of non-damaged points. Overall the conclusion being that our novel test statistic is conservative and has low false positive rate.

**Figure 5.**
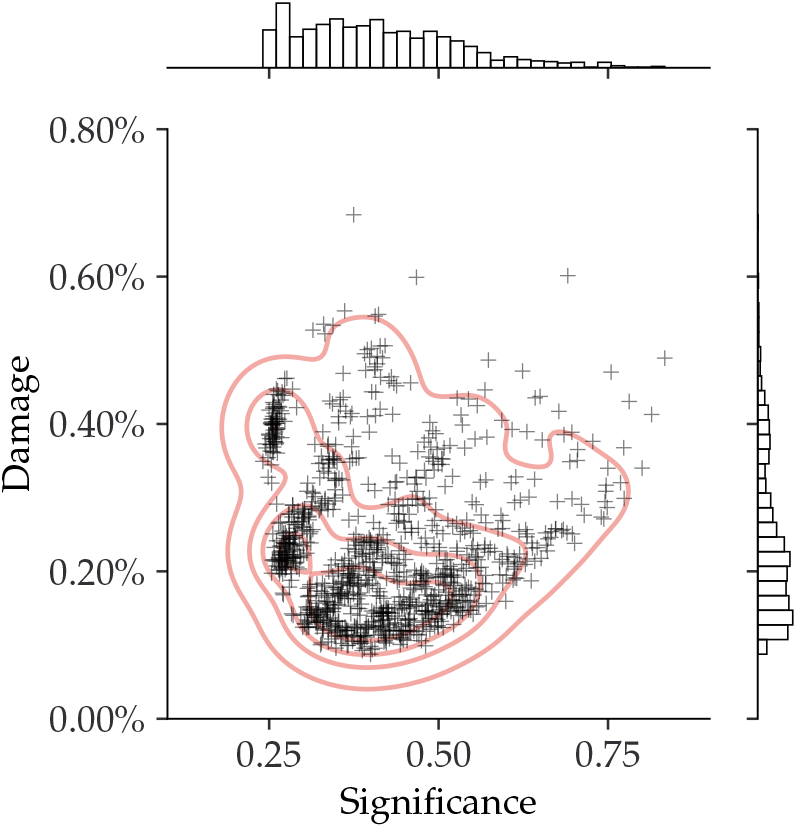
Inferred damage of modern, simulated data (single-genome). The plot shows the inferred damage estimates of 1000 replicates, each with 1000 reads and no artificial ancient damage applied. Each single cross corresponds to a simulation and the red lines outlines the kernel density estimate (KDE) of the damage estimates. The marginal distributions are shown as histograms next to the scatter plot.

### 4.2 Metagenomic simulation results

With the full metagenomic simulation pipeline we can further probe the performance of `metaDMG`. By considering the different metagenomic scenarios, see ***Table 1***, at different steps in the pipeline, we are able to show that `metaDMG` provides relevant and accurate damage estimates.

To verify that the risk of getting false positives is non-significant, we run `metaDMG` on the metagenomic assemblages after fragmentation with FragSim, but before any no deamination with Deam-Sim has yet been added. We find that the previously established relaxed significance threshold (*D*_fit_ > 1% and *Z*_fit_ > 2) correctly filters out all of the taxa, see ***Figure 6***. This is as expected, as there has not yet been added any artificial post mortem damage in the form of deamination.

**Figure 6.**
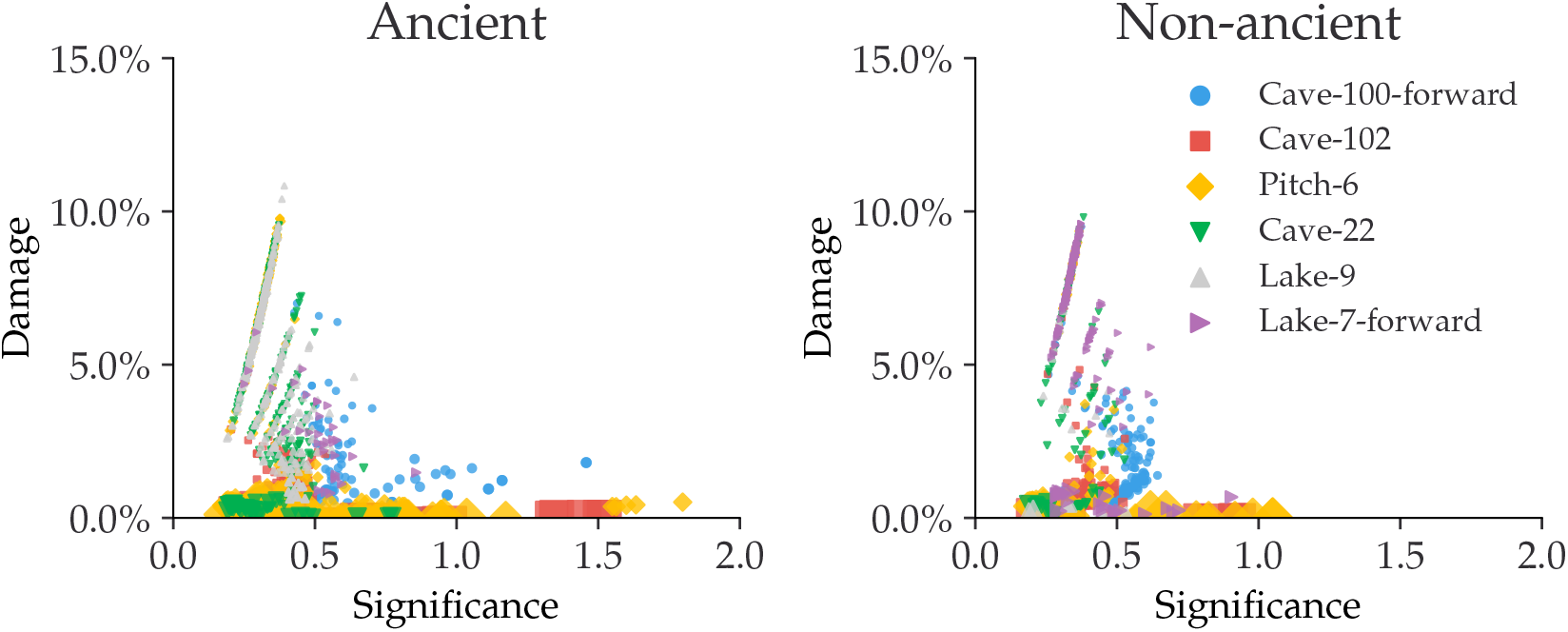
Estimated amount of damage as a function of significance for metagenomic simulations. This figure shows the metagenomic simulations after FragSim has been applied, but before including any deamination or sequencing errors. We generate both non-ancient and ancient taxa in the simulation pipeline. The left subfigure shows the damage of the ancient taxa and similarly for the non-ancient taxa in the right subfigure.

We see a clear difference in the damage estimates between the ancient and the non-ancient taxa once we add deamination with DeamSim and sequencing errors with ART, see ***Figure 7***. The non-ancient taxa would still not pass the relaxed threshold, in contrast to the taxa in the ancient samples.

**Figure 7.**
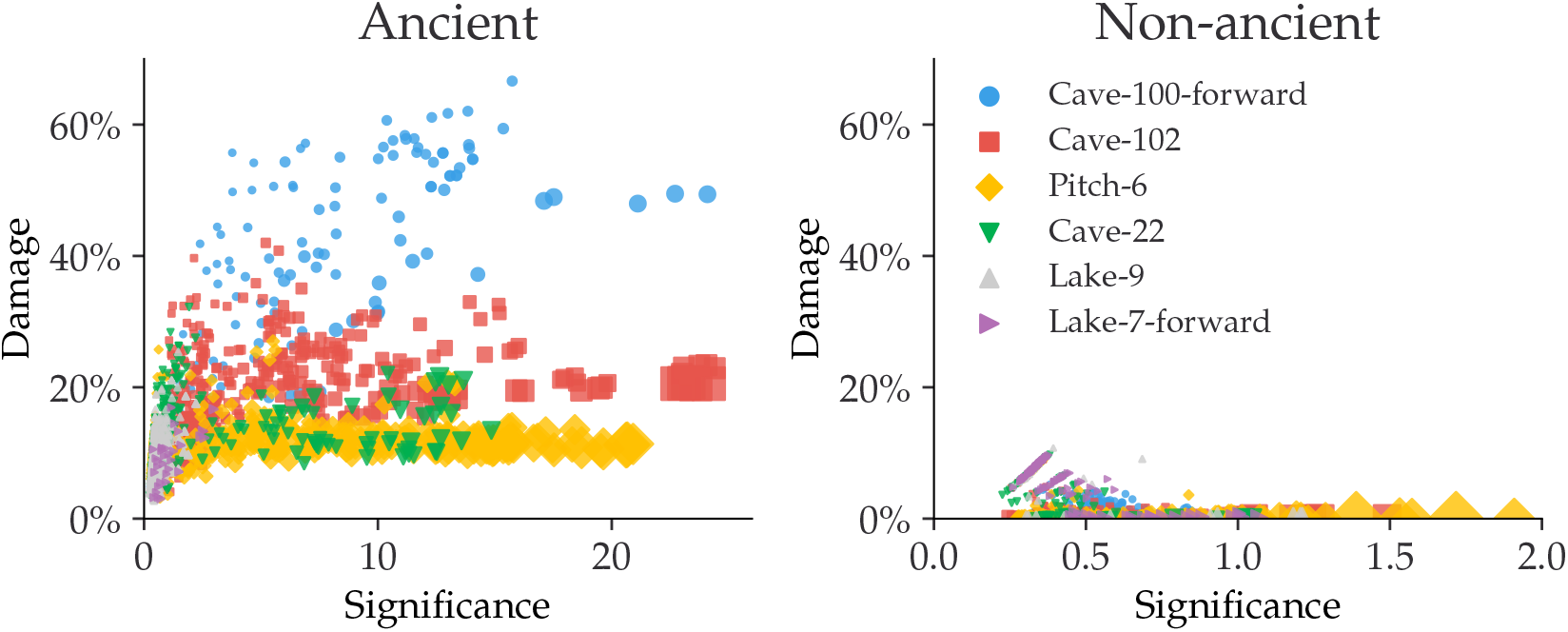
Estimated amount of damage as a function of significance for metagenomic simulations. This figure shows the metagenomic simulations after fragmentation, deamination, and sequencing errors have been applied. The left subfigure shows the damage of the ancient taxa and similarly for the non-ancient taxa in the right subfigure.

The results of ***Figure 7*** are summarized in ***Table 2***. We find that Cave-100-forward, Cave-102, Pitch-6 all have more than 60% of their ancient taxa correctly labelled as damaged according to the relaxed threshold, while it for Cave-22 and Lake-7-forward is a bit lower and Lake-9 does not show any clear support of damage. However, once we condition on the requirement of having more than 100 reads, the fraction of ancient taxa correctly identified as ancient increases to more than 90% for most of the samples. A small investigation of one of the ancient taxa (Stenotrophomonas Maltophilia) in the simulation that did not meet the criteria to be ancient `metaDMG`, i.e. a false negative, can be found in ***Appendix 6***.

**Table 2.** `metaDMG` damage results for the six different metagenomic simulations. The first column is the total number of taxa, the second column is the total number of taxa that would pass the threshold of *D*_fit_ > 1% and *Z*_fit_ > 2, the third column is the number of taxa with more than 100 reads, and the final column is the number of taxa with more than 100 reads that also do pass the cut.

| Sample | Total | Pass | +100 Reads | +100 Reads and Pass |
| --- | --- | --- | --- | --- |
| Cave-100-forward | 135 | 107 79.3% | 88 | 87 98.9% |
| Cave-102 | 500 | 326 65.2% | 309 | 285 92.2% |
| Pitch-6 | 415 | 260 62.7% | 274 | 260 94.9% |
| Cave-22 | 393 | 71 18.1% | 73 | 69 94.5% |
| Lake-9 | 410 | 2 0.5% | 8 | 0 0% |
| Lake-7-forward | 32 | 4 12.5% | 6 | 4 66.7% |

### 4.3 Real Data

The results from running the real metagenomic data through the `metaDMG` pipeline show clear evidence of taxa with significant DNA damage present in the metagenome and a layered pattern similar to what was observed in the simulated ancient metagenomes, see ***Figure 8***.

**Figure 8.**
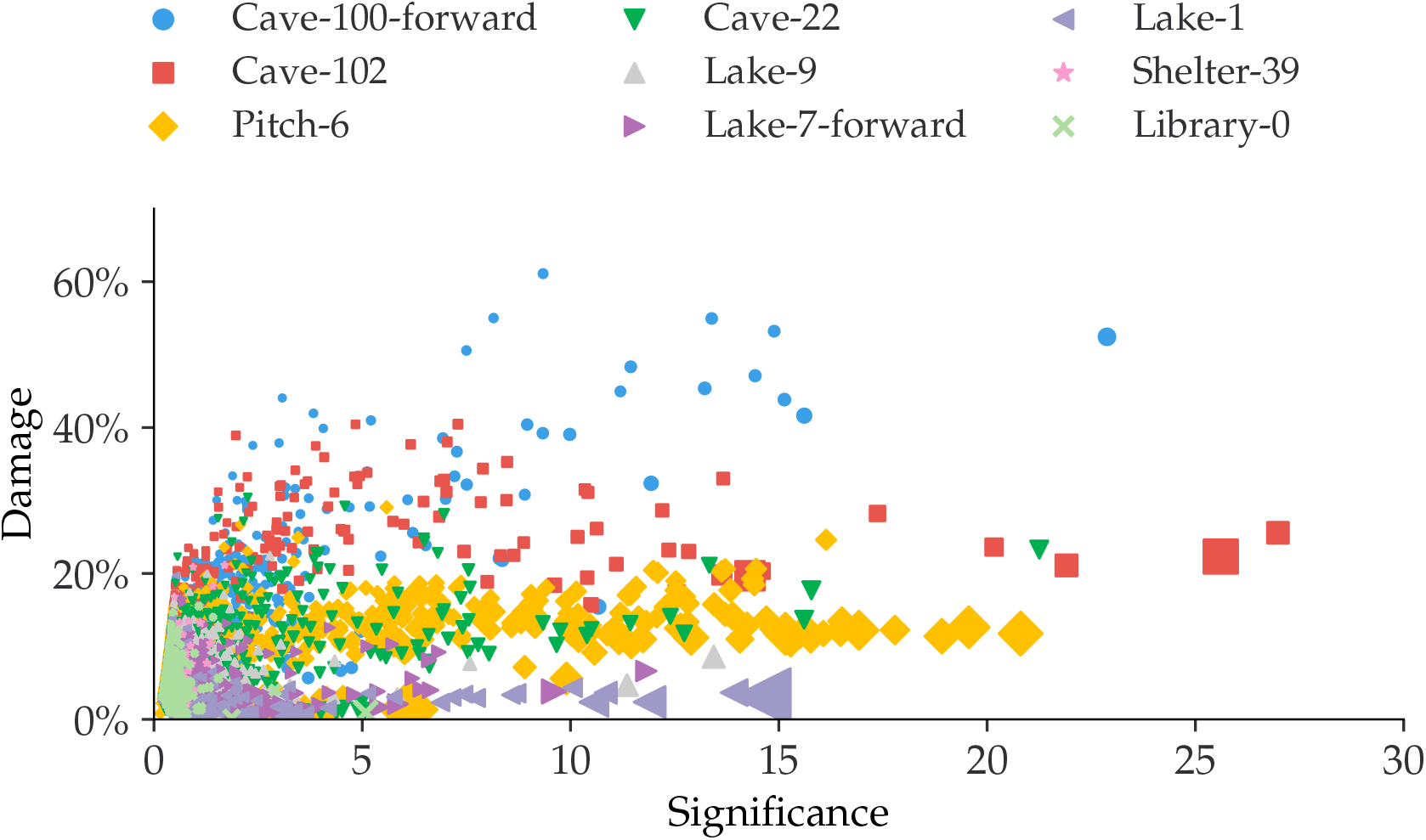
Estimated amount of damage as a function of significance using the real data, see ***Table 1***.

As DNA damage is not a function of time, we cannot expect that there is a direct relation between damage and time, however, we do see that the oldest samples, Cave-100 and Cave-102, see ***Table 1***, which are 100 and 102 thousand years BP, show the highest amount of damage of all the metagenomes. Both the Pitch-6 and Cave-22 samples, which are 6 and 22 thousand year old and thus younger than two above mentioned cave samples, have almost similar levels of damage. This is not unexpected as the micro environment surrounding the layer in which the metagenome was found plays a significant role in the state of DNA. In our case, the younger Pitch-6 derives from a water logged but open air site, while the Cave-22 sample was obtained in dry but cool (~11 degree Celsius year around) cave layers.

The metagenomes with the least DNA damage are the ones from the lake sediments (Lake-1, Lake-7 and Lake-9). These samples do show some taxa with significant DNA damage, although they do not have a strong damage signal.

Importantly, we find that in the true metagenomes, `metaDMG` is able to assign low significance to the taxa that likely are not damaged or that have too little data, see e.g. the upper right hand corner of ***Figure 9***. This subfigure shows the damage plot for the Gallus Gallus species (red junglefowl) from the Lake-1 sample. This particular species only has *D*_fit_ = 2.2% and *Z*_fit_ = 1.0, which does not satisfy the relaxed DNA damage threshold (*D*_fit_ > 1%, *Z*_fit_ > 2). In addition to the Gallus Gallus species, ***Figure 9*** further shows examples of species with large amounts of data (Homo Sapiens in the Pitch-6 sample and Crocuta Crocuta in the Cave-100 sample, based only on forward data), and an example of medium damage (Equisetum Arvense in Lake-7, based only on forward data).

**Figure 9.**
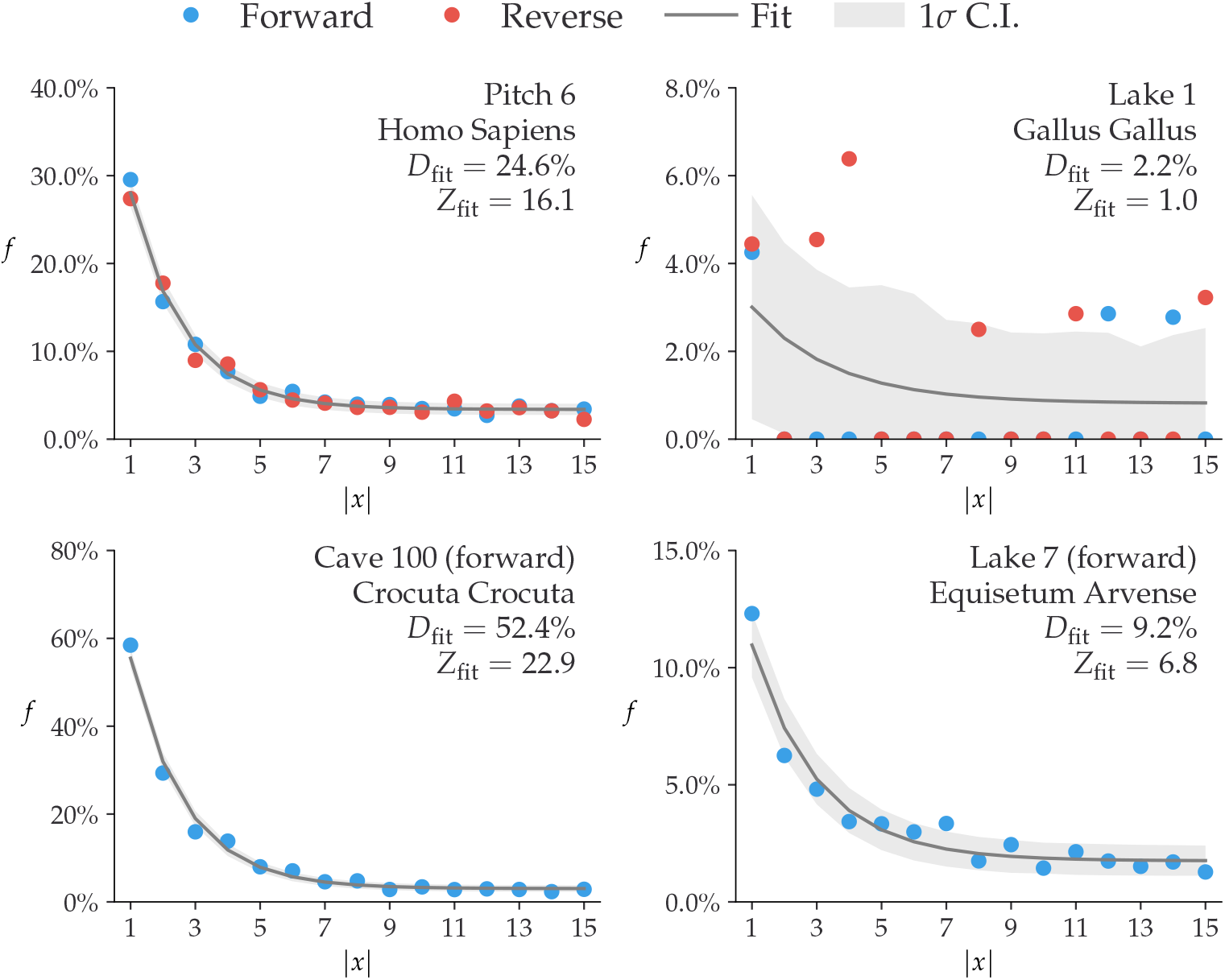
Damage plots of four representative species from the real-data metagenomic samples, see ***Table 1***. Each subfigure shows the damage rate *f* (*x*) = *k*(*x*)∕*N*(*x*) as a function of position *x* for both forward (C→T) and reverse (G→A). The `metaDMG` fit is shown in grey with the 68% credible intervals as shaded regions. In the upper right corner of each subfigure, the information about the sample and the species together with the `metaDMG` damage estimates is shown.

Interestingly, and of high importance for downstream interpretation, is that for certain samples, some taxa were found to have a high significance although with lower DNA damage than what is observed across the given metagenome as a whole. This underlines the need to evaluate the DNA damage variation within each metagenome, perform a proper outlier test and the basic setting of logical thresholds.

We find that when using the relaxed DNA damage threshold, `metaDMG` falsely classifies a single of the taxa from the control test Library-0 as being ancient. However, with a more conservative damage threshold (*D*_fit_ > 2%, *Z*_fit_ > 3, more than 100 reads), none of the taxa from the library control are classified as ancient.

### 4.4 Bayesian vs. MAP

Due to the higher computational burden of computing the full Bayesian model compared to the faster, approximate MAP model in samples with several thousand taxa, the MAP model is in practice the model of choice due to lower computational complexity. We compared the performance of *D*_fit_ and *Z*_fit_ on the real datasets in ***Table 1***, see ***Figure 10***. This figure compares the estimated damage between the Bayesian model and the MAP model (left subfigure) and the estimated significances (right subfigure) for taxa passing a threshold of *D*_fit_ > 1%, *Z*_fit_ > 2, and more than 100 reads. The figure shows that the vast majority of taxa map 1:1 between the Bayesian and the MAP model. It should be noticed that the taxa with the worst correspondence in damage estimates are all based on forward-only fits, i.e. with no information from the reverse strand, which leads to less data to base the fits on. For the comparison with no thresholds applied, see ***Figure S23*** in ***Appendix 7***. We recommend to use the full, Bayesian model in the case of extremely low-coverage data or when used on only a small number of taxa (e.g. when using `metaDMG` in global-mode).

**Figure 10.**
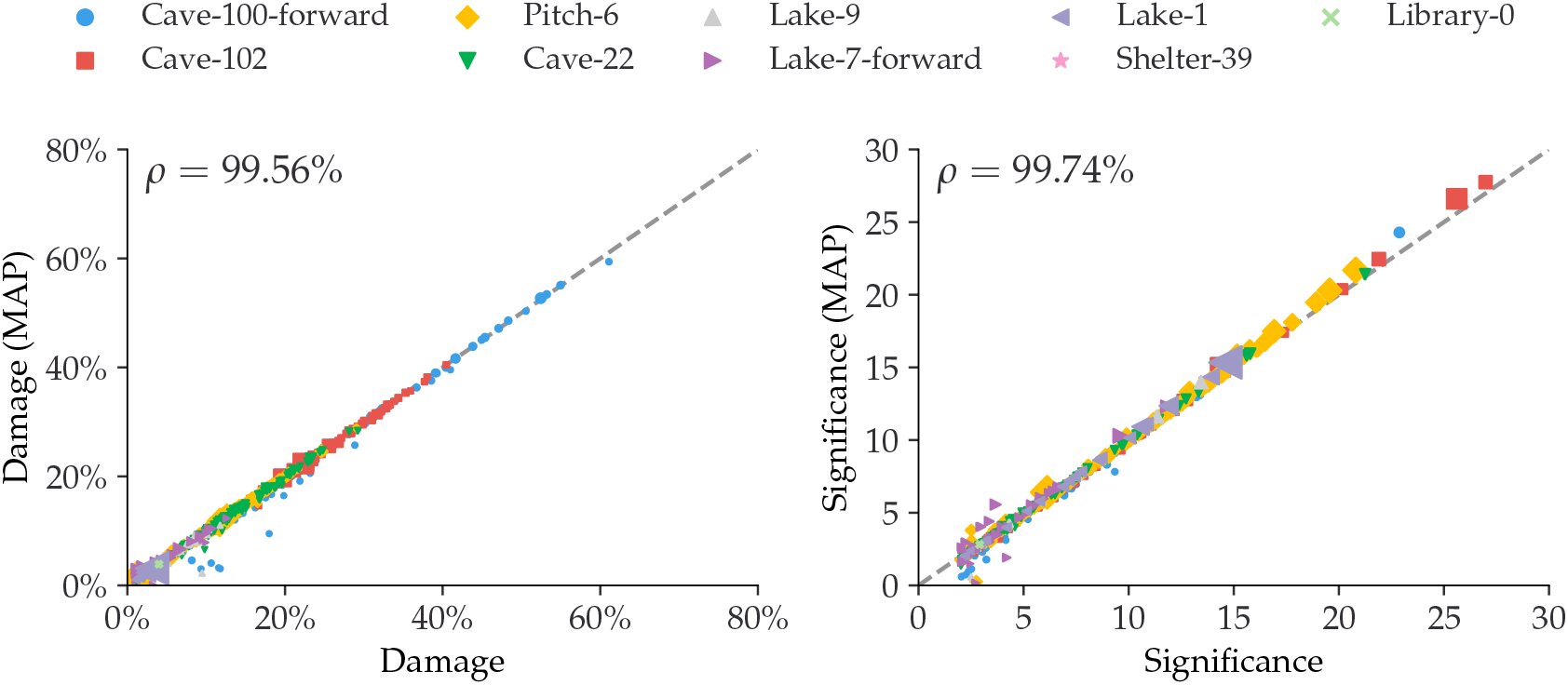
Comparison between the full Bayesian model and the fast, approximate, MAP model for the estimated damage and significance. The figure shows data after a loose cut of *D*_fit_ > 1%, *Z*_fit_ > 2 and more than 100 reads. The dashed, grey line shows the 1:1 ratio and the correlation, *ρ*, is shown in the upper left corner.

### 4.5 Existing Methods

To our knowledge there are not currently available methods for assesing and quantifying postmortem DNA damage in a metagenomic context. We compare the performance of the *D*_fit_ statistic in `metaDMG` to existing methods such as those found in PyDamage (Borry et al., 2021). Since PyDamage is based solely on single genome analysis we use the non-LCA mode of `metaDMG`. This mode iterates through the different referenceIDs for all mapped reads and estimates the damage for each. In general, we find that `metaDMG` is more conservative, accurate and precise in its damage estimates.

One example of this can be found in ***Figure 11***, which shows both the `metaDMG` and PyDamage results of the simulations described in ***subsection 3.1***, in particular the 100 replications of the Homo Sapiens single-genome with 100 reads and 15% added artificial damage (and a fragment length distribution with mean 60). ***Figure 11*** shows that the `metaDMG` estimates are between 5% and 25% damage, while PyDamage estimates up to more than 50% damage, in a sample with 15% artificially added damage. The comparisons between `metaDMG` and PyDamage for the other sets of simulation parameters can be found in ***Figure S24***–***Figure S31*** in ***Appendix 8***.

**Figure 11.**
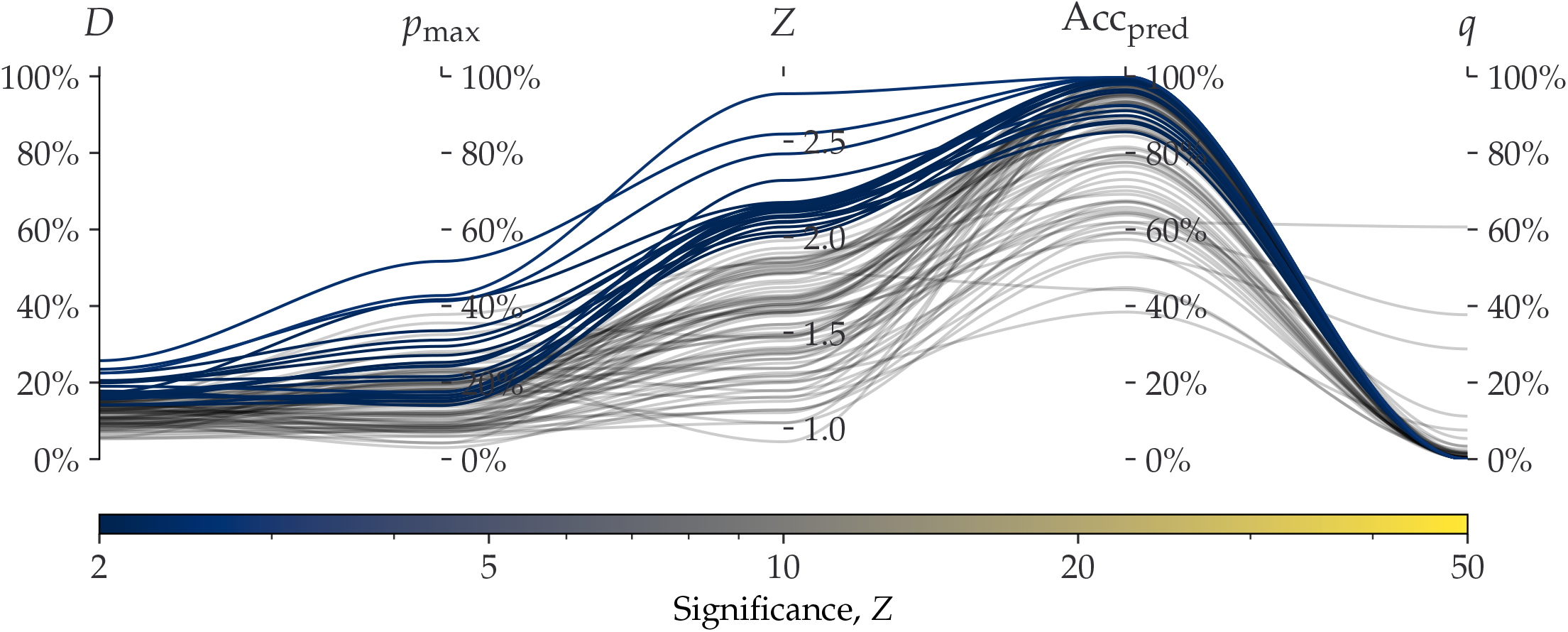
Parallel coordinates plot comparing `metaDMG` and PyDamage for the Homo Sapiens single-genome simulation with 100 reads and 15% added artificial damage. The two first axes show the estimated damage: *D*_fit_ by `metaDMG` and *p*_max_ by PyDamage. The following two axes show the fit quality: significance (*Z*_fit_) by `metaDMG` and the predicted accuracy (Acc_pred_) by PyDamage. The final axis shows the *q*-value by PyDamage. Each of the 100 replications are plotted as single lines. Replications passing the relaxed `metaDMG` damage threshold (*D*_fit_ > 1% and *Z*_fit_ > 2) are shown in color proportional to their significance. Replications that did not pass are shown in semi-transparent black lines.

To compare the computational performance, we use the real-life Pitch-6 sample (i.e. non-simulated), see ***Table 1***. This alignment file (in BAM-format) takes up 857 MB of space and has 3.7 millions reads with a total of 19 million alignments to 11.433 unique taxa. When using only a single core, PyDamage took 1105s to compute all fits, while `metaDMG` took 88s, a factor of 12.6x faster. The rest of the timings are shown in ***Table 3***. PyDamage requires the alignment files to be sorted by chromosome position and be supplied with an index file, allowing it to iterate fast through the alignment file, at the expense of computational load before running the actual damage estimation. `metaDMG` on the other hand requires the reads to be sorted by name to minimize the time it takes to run the LCA.

**Table 3.** Computational performance of PyDamage and `metaDMG`. The table contains the times it takes to run either PyDamage or `metaDMG` on the full Pitch-6 sample containing 11.433 taxa. The timings are shown for both single-processing case (1 core) and multi-processing (2 and 4 cores). The timings were performed on a Macbook M1 Pro model from 2021. “12.6x” means that `metaDMG` was 12.6 times faster than PyDamage for that particular test.

| Cores | Pydamage (s) | metaDMG (s) | Improvement (x) |
| --- | --- | --- | --- |
| 1 | 1105 | 88 s | 12.6 |
| 2 | 592 | 66 s | 9.0 |
| 4 | 398 | 54 s | 7.4 |

## 5 DISCUSSION

To our knowledge there are no currently available methods other than `metaDMG` that is geared towards damage analysis in a metagenomic setting. It is the first general framework designed specifically for the quantification of ancient damage in all contexts. The toolkit contains various interlinked and independent modules including a state-of-the-art graphical user interface that allow researchers to explore their data.

Multiple areas of future improvements exists. Currently, our novel test statistic for the damage estimation *D*_fit_ is based on a statistical model where we only consider the C→T and G→A transitions and where each taxa is modelled as being fully independent, even for closely related species when provided a taxanomic tree. This could be improved upon with the use of a hierarchical model were information across taxonomic leaf nodes is shared. The current implementation, however, allows for easy parallelization of the individual fits which reduces the time spent on the inference. In addition to the mismatch matrices, another improvement would be to include the read length distribution as a covariate in the damage model, as, in addition to deamination, the fragment length distribution is also an indicator of ancient damage (Dabney, Meyer, and Pääbo, 2013; Peyrégne and Prüfer, 2020).

We show that the *D*_fit_ statistic that `metaDMG` provides is accurate across different damage levels and different number of reads. In the single-genome reference case, we further show that the estimates are stable across different species and fragment length distributions. In addition to this, we find that the results are independent of the contig size, in contrast to PyDamage (Borry et al., 2021).

The basis for the *D*_fit_ statistic is the leaf node mismatch matrices which contains the raw observed substitution frequencies. The computation of these could also take into account the computed mapping uncertainty and the uncertainty of the assigned called nucleotide. We include a regression approach for stabilizing the mismatch matrices across all covariates but this requires much more data than our current approach. Rather than regressing on all covariates, it might also be more biological meaningfull to regress on the four Briggs parameters.

In our toolkit we have included the PMDtools approach (Skoglund et al., 2014) that allows for the separation of highly damaged reads from undamaged reads. The method offers a reasonable way to distinguish the endogenous ancient DNA from possible modern contamination. But this method may suffer from the fact that some fixed empirical parameters are applied. A possible extension can be using several statistics estimated from the specific sample (e.g., taxa specific *D*_fit_ and the ancient fragment lengths) as priors in an empirical Bayes inference framework to learn the categories of reads unsupervisedly.

Our research indicate that the `metaDMG` results are conservative with very low false positive rates. This is particularly important with metagenomic samples as the number of taxa, and thus the number of damage estimates, tend to be large. As the number of fits increases, we strongly believe that a graphical user interface is important. This helps to select and filter the fit results, and to better understand the data at hand. We have tested `metaDMG` using a state of the art metagenomic simulation pipeline based on multiple metagenomic real-life sample from a variety of different environments. We hope that `metaDMG` can improve the knowledge about DNA damage degradation in different environments and be the foundation of a more general, metagenomic ancient damage study.

## 6 AUTHOR CONTRIBUTIONS

CM developed and implemented the damage model and all aspect of the python code including the CLI, all fits, and the dashboard. TP helped develop the model and with statistical discussions. TSK implemented the C/C++ code relating to the lowest common ancestor and mismatch matrices. LZ implemented the PMDtools and full multinomial regression subfunctionality. AFG and MWP designed the metagenomic simulation study and the application of `metaDMG` to real data. CM and MWP ran all analyses. CM, MWP and TSK initiated and designed the project. All authors contributed to writing the manuscript.

## Appendix 1 PMDTOOLS

Three non-mutually exclusive events can lead to an observation of C→T or G→A (Skoglund et al., 2014), namely (i) a true biological polymorphism (occurring at rate *π*), (ii) a sequencing errors (rate *ϵ*, can be extracted from the base quality scores of the site on the sampled strand), and (iii) in the case of damaged DNA, the damaged nucleotide frequencies are assumed to be only related to its position from either termini of the ancient fragment (C→T from 5’ end, and G→A from 3’ end). The error probability of the postmortem nucleotide misincorporation is under the pmdtools model given by:

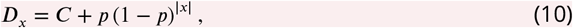

here *C* = 0.01 and *p* = 0.3 are both suitable constants. Skoglund et al., 2014 defines the likelihood ratio of a strand between the PMD model and the NULL model as its postmortem damage score (PMDS),

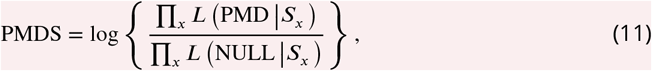

The reads with the PMDS exceeding an empirical p-value threshold can then be used for filtering intensively damaged fragments.

## Appendix 2 MULTINOMIAL LOGISTIC REGRESSIONS

## Full Multinomial Logistic Regression

Postmortem damages have impacts on the next generation sequencing reads. A common phenomenon is the increasing of the calling error rates from nucleotide C→T due to the cytosine deamination process. Unawareness of this will lead to inaccurate inferences. Evidences show that the magnitude of such changes are related to the positions the site is within a read (the fraction of the ancient DNA). Here we present four slightly different ways (i.e., full unconditional regression, full conditional regression, folded unconditional regression and folded conditional regression) to unveil the relationship between the calling error rates and the mismatching reference/read pairs as well as the site positions within a read. The methods are based on the multinomial logistic regressions.

## Data Description

We perform the regressions based on the summary statistic of the mismatch matrix,i.e., *M*(*x*), which is a table which contains the counts of reads of different reference/read categories (in total 16) and positions on the forward/reversed strand (15 positions on each direction). ***Table S1*** and ***Table S2*** give an example of the data format we use for the inference.

**Appendix 2–table S1.**
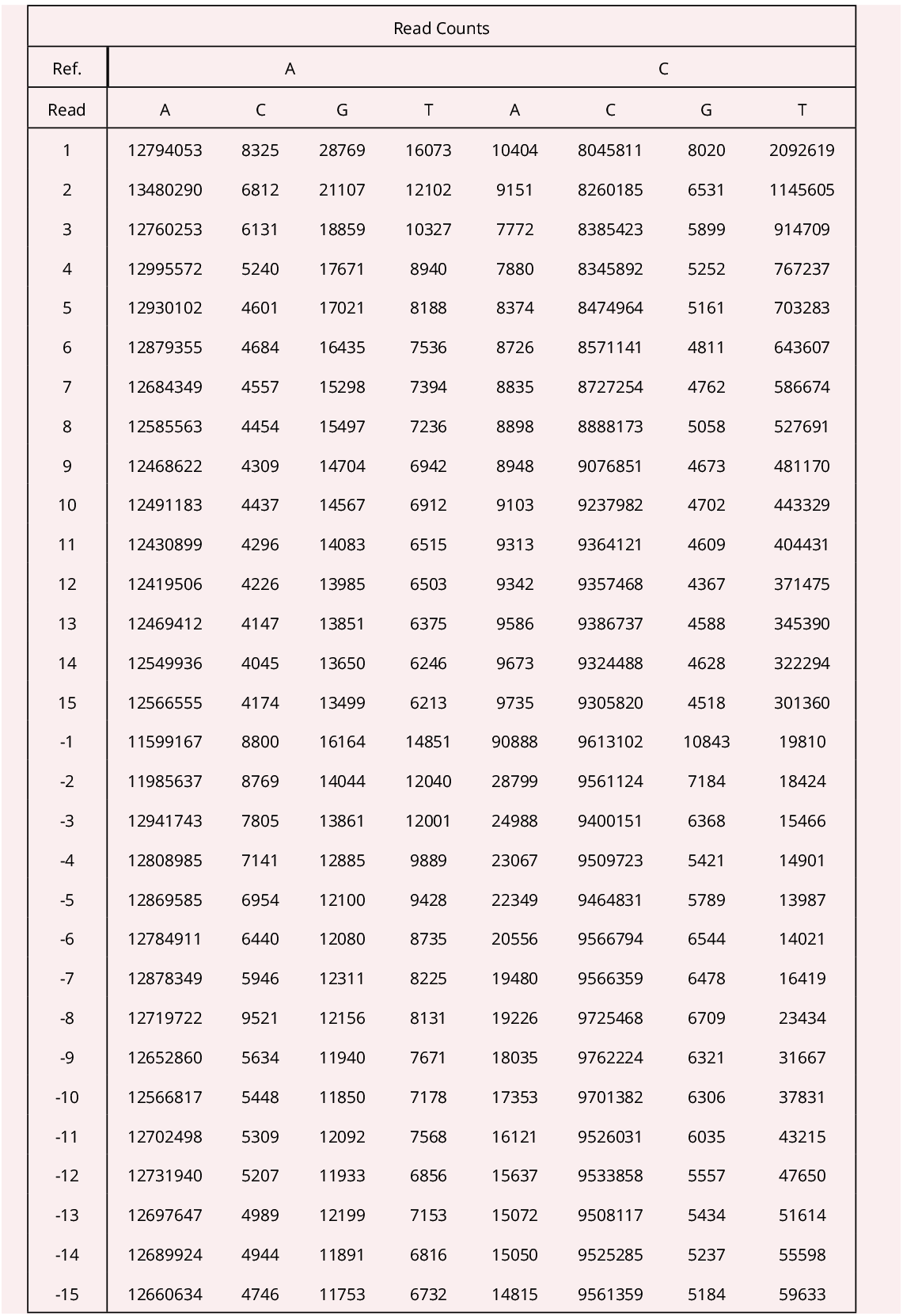
The read counts per position given the reference nucleotides are A or C of an ancient human data. The negative position indices are the position on the reversed strand. In the manuscript, the elements (the values of a specific nucleotide read counts per position given the reference nucleotide is A or C) in this table are denoted as *M*_*A*→*i*_(*x*) or *M*_*C*→*i*_(*x*).

| Read Counts |  |  |  |  |  |  |  |  |
| --- | --- | --- | --- | --- | --- | --- | --- | --- |
| Ref. | A |  |  |  | C |  |  |  |
| Read | A | C | G | T | A | C | G | T |
| 1 | 12794053 | 8325 | 28769 | 16073 | 10404 | 8045811 | 8020 | 2092619 |
| 2 | 13480290 | 6812 | 21107 | 12102 | 9151 | 8260185 | 6531 | 1145605 |
| 3 | 12760253 | 6131 | 18859 | 10327 | 7772 | 8385423 | 5899 | 914709 |
| 4 | 12995572 | 5240 | 17671 | 8940 | 7880 | 8345892 | 5252 | 767237 |
| 5 | 12930102 | 4601 | 17021 | 8188 | 8374 | 8474964 | 5161 | 703283 |
| 6 | 12879355 | 4684 | 16435 | 7536 | 8726 | 8571141 | 4811 | 643607 |
| 7 | 12684349 | 4557 | 15298 | 7394 | 8835 | 8727254 | 4762 | 586674 |
| 8 | 12585563 | 4454 | 15497 | 7236 | 8898 | 8888173 | 5058 | 527691 |
| 9 | 12468622 | 4309 | 14704 | 6942 | 8948 | 9076851 | 4673 | 481170 |
| 10 | 12491183 | 4437 | 14567 | 6912 | 9103 | 9237982 | 4702 | 443329 |
| 11 | 12430899 | 4296 | 14083 | 6515 | 9313 | 9364121 | 4609 | 404431 |
| 12 | 12419506 | 4226 | 13985 | 6503 | 9342 | 9357468 | 4367 | 371475 |
| 13 | 12469412 | 4147 | 13851 | 6375 | 9586 | 9386737 | 4588 | 345390 |
| 14 | 12549936 | 4045 | 13650 | 6246 | 9673 | 9324488 | 4628 | 322294 |
| 15 | 12566555 | 4174 | 13499 | 6213 | 9735 | 9305820 | 4518 | 301360 |
| -1 | 11599167 | 8800 | 16164 | 14851 | 90888 | 9613102 | 10843 | 19810 |
| -2 | 11985637 | 8769 | 14044 | 12040 | 28799 | 9561124 | 7184 | 18424 |
| -3 | 12941743 | 7805 | 13861 | 12001 | 24988 | 9400151 | 6368 | 15466 |
| -4 | 12808985 | 7141 | 12885 | 9889 | 23067 | 9509723 | 5421 | 14901 |
| -5 | 12869585 | 6954 | 12100 | 9428 | 22349 | 9464831 | 5789 | 13987 |
| -6 | 12784911 | 6440 | 12080 | 8735 | 20556 | 9566794 | 6544 | 14021 |
| -7 | 12878349 | 5946 | 12311 | 8225 | 19480 | 9566359 | 6478 | 16419 |
| -8 | 12719722 | 9521 | 12156 | 8131 | 19226 | 9725468 | 6709 | 23434 |
| -9 | 12652860 | 5634 | 11940 | 7671 | 18035 | 9762224 | 6321 | 31667 |
| -10 | 12566817 | 5448 | 11850 | 7178 | 17353 | 9701382 | 6306 | 37831 |
| -11 | 12702498 | 5309 | 12092 | 7568 | 16121 | 9526031 | 6035 | 43215 |
| -12 | 12731940 | 5207 | 11933 | 6856 | 15637 | 9533858 | 5557 | 47650 |
| -13 | 12697647 | 4989 | 12199 | 7153 | 15072 | 9508117 | 5434 | 51614 |
| -14 | 12689924 | 4944 | 11891 | 6816 | 15050 | 9525285 | 5237 | 55598 |
| -15 | 12660634 | 4746 | 11753 | 6732 | 14815 | 9561359 | 5184 | 59633 |

**Appendix 2–table S2.**
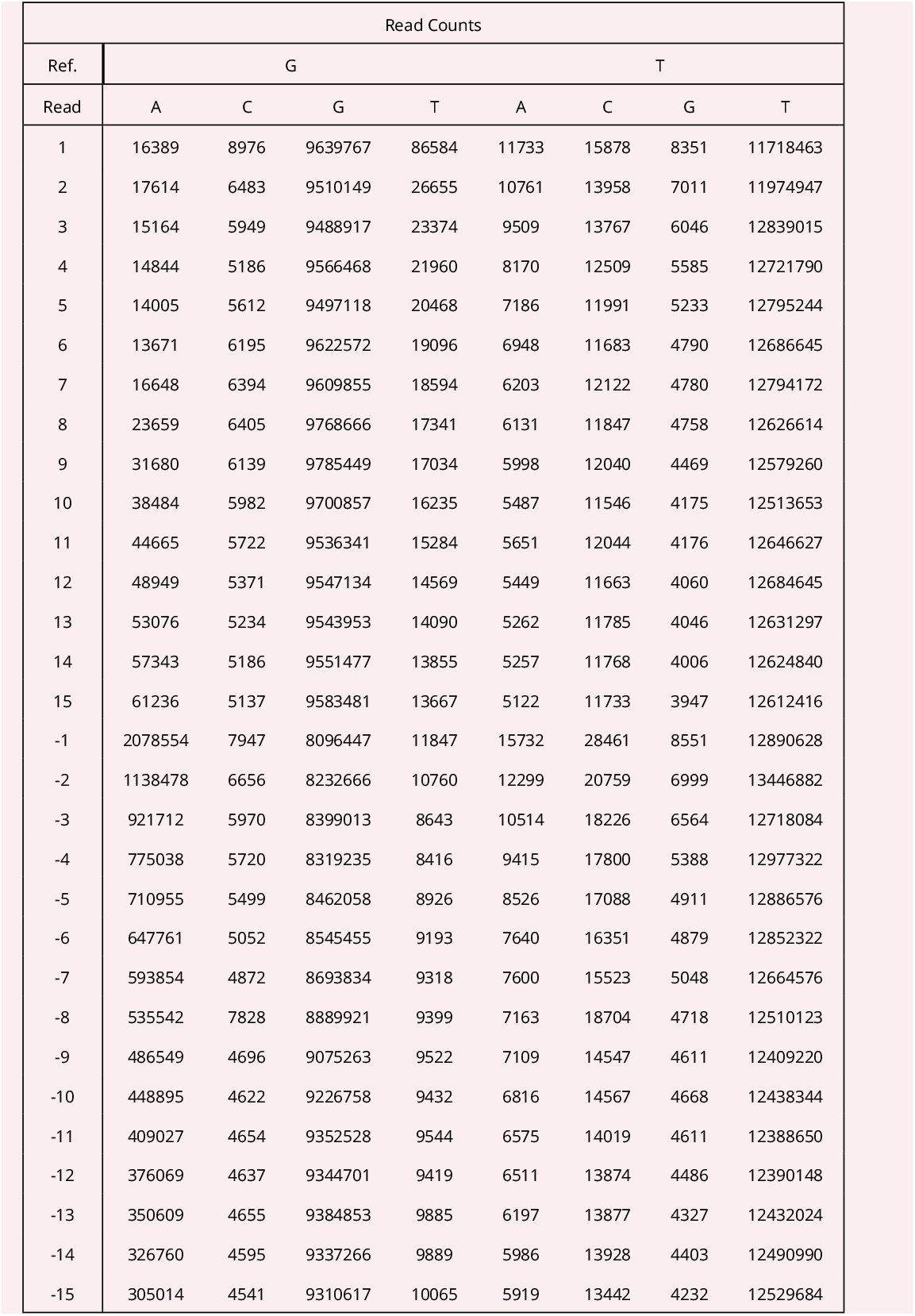
The read counts per position given the reference nucleotides are G or T of the same human data as in Table S1. The negative position indices are the position on the reversed strand. In the manuscript, the elements (the values of a specific nucleotide read counts per position given the reference nucleotide is G or T) in this table are denoted as *M*_*G*→*i*_(*x*) or *M*_*T*→*i*_(*x*).

| Read Counts |  |  |  |  |  |  |  |  |
| --- | --- | --- | --- | --- | --- | --- | --- | --- |
| Ref. | G |  |  |  | T |  |  |  |
| Read | A | C | G | T | A | C | G | T |
| 1 | 16389 | 8976 | 9639767 | 86584 | 11733 | 15878 | 8351 | 11718463 |
| 2 | 17614 | 6483 | 9510149 | 26655 | 10761 | 13958 | 7011 | 11974947 |
| 3 | 15164 | 5949 | 9488917 | 23374 | 9509 | 13767 | 6046 | 12839015 |
| 4 | 14844 | 5186 | 9566468 | 21960 | 8170 | 12509 | 5585 | 12721790 |
| 5 | 14005 | 5612 | 9497118 | 20468 | 7186 | 11991 | 5233 | 12795244 |
| 6 | 13671 | 6195 | 9622572 | 19096 | 6948 | 11683 | 4790 | 12686645 |
| 7 | 16648 | 6394 | 9609855 | 18594 | 6203 | 12122 | 4780 | 12794172 |
| 8 | 23659 | 6405 | 9768666 | 17341 | 6131 | 11847 | 4758 | 12626614 |
| 9 | 31680 | 6139 | 9785449 | 17034 | 5998 | 12040 | 4469 | 12579260 |
| 10 | 38484 | 5982 | 9700857 | 16235 | 5487 | 11546 | 4175 | 12513653 |
| 11 | 44665 | 5722 | 9536341 | 15284 | 5651 | 12044 | 4176 | 12646627 |
| 12 | 48949 | 5371 | 9547134 | 14569 | 5449 | 11663 | 4060 | 12684645 |
| 13 | 53076 | 5234 | 9543953 | 14090 | 5262 | 11785 | 4046 | 12631297 |
| 14 | 57343 | 5186 | 9551477 | 13855 | 5257 | 11768 | 4006 | 12624840 |
| 15 | 61236 | 5137 | 9583481 | 13667 | 5122 | 11733 | 3947 | 12612416 |
| -1 | 2078554 | 7947 | 8096447 | 11847 | 15732 | 28461 | 8551 | 12890628 |
| -2 | 1138478 | 6656 | 8232666 | 10760 | 12299 | 20759 | 6999 | 13446882 |
| -3 | 921712 | 5970 | 8399013 | 8643 | 10514 | 18226 | 6564 | 12718084 |
| -4 | 775038 | 5720 | 8319235 | 8416 | 9415 | 17800 | 5388 | 12977322 |
| -5 | 710955 | 5499 | 8462058 | 8926 | 8526 | 17088 | 4911 | 12886576 |
| -6 | 647761 | 5052 | 8545455 | 9193 | 7640 | 16351 | 4879 | 12852322 |
| -7 | 593854 | 4872 | 8693834 | 9318 | 7600 | 15523 | 5048 | 12664576 |
| -8 | 535542 | 7828 | 8889921 | 9399 | 7163 | 18704 | 4718 | 12510123 |
| -9 | 486549 | 4696 | 9075263 | 9522 | 7109 | 14547 | 4611 | 12409220 |
| -10 | 448895 | 4622 | 9226758 | 9432 | 6816 | 14567 | 4668 | 12438344 |
| -11 | 409027 | 4654 | 9352528 | 9544 | 6575 | 14019 | 4611 | 12388650 |
| -12 | 376069 | 4637 | 9344701 | 9419 | 6511 | 13874 | 4486 | 12390148 |
| -13 | 350609 | 4655 | 9384853 | 9885 | 6197 | 13877 | 4327 | 12432024 |
| -14 | 326760 | 4595 | 9337266 | 9889 | 5986 | 13928 | 4403 | 12490990 |
| -15 | 305014 | 4541 | 9310617 | 10065 | 5919 | 13442 | 4232 | 12529684 |

The terminology used here might not be standard. The term full regression here is to distinguish itself from the folded regression discussed later, which simply means inferring the coefficients of forward strand and reversed strand separately. Full regression includes both unconditional regression and conditional regression. The unconditional regression’s objective is to infer the probability of observing a read of nucleotide *j* and its reference is *i* at position *x*, i.e., *P*_*i*→*j*_ (*x*) while the conditional regression’s target is to estimate the probability of observing a read of nucleotide *j* <u>given</u> its reference is *i* at position *x*, i.e., *P*_*j*|*i*_(*x*). Their relationship is as follows:

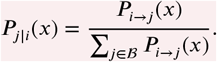

So in fact, unconditional regression can give us more detailed inferred results (extra information the nucleotide composition per position of the reference, which may be related to the prepared libraries).

## Unconditional Regression Likelihood

The unconditional regression’s log-likelihood function is defined as follows,

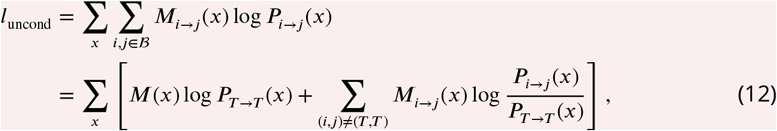

where 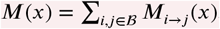. According to the multinomial logistic regression, we assume,

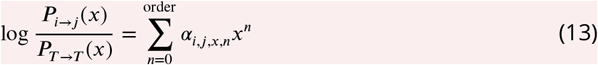

Applying Equation 13 to Equation 12, we have

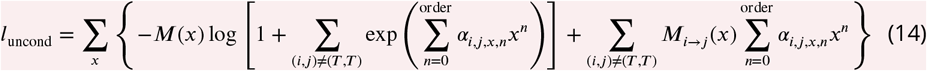

The number of inferred parameters (*α*_*i*,*j*,*x*,*n*_), for the full conditional regression is 30×(order + 1). And the relevant derivatives of the unconditional regression likelihood are as follows,

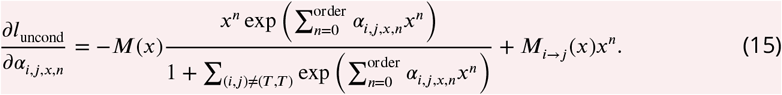

## Conditional Regression Likelihood

Viewed as the sum of log-likelihoods given the reference nucleotide 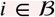, the conditional regression’s log-likelihood function is,

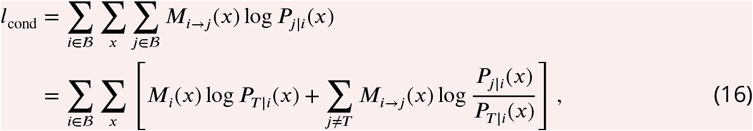

where 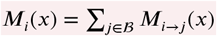. Furthermore, if we assume,

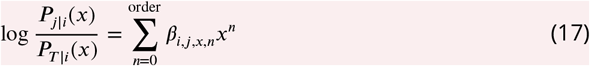

By applying Equation 17 to Equation 16, we can obtain,

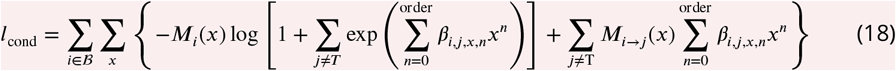

The number of inferred parameters (*β*_*i*,*j*,*x*,*n*_) for the full unconditional regression is 24 × (order + 1). And the relevant derivatives of the conditional likelihood are as follows,

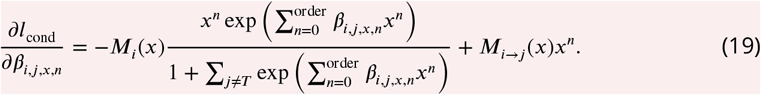

## Folded Multinomial Logistic Regression

The folded regressions use the same log-likelihood functions as the full regression (i.e., Equation 14 and 18) but are conducted based on a presumable symmetric PMD pattern, i.e., the probability of *C* → *T* at the position *x* of an random chosen ancient DNA strand is assumed to equal to the probability of *G* → *A* at the position −*x*. Such an theoretical assumption go match the current ancient library preparation process (Dabney, Meyer, and Pääbo, 2013; Henriksen, Zhao, and T. Korneliussen, 2022).

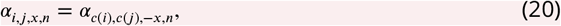

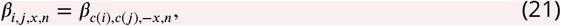

where *c*(*i*) means the complimentary nucleotide of the nucleotide *i*, e.g., *c*(*A*) = *T* and *c*(*G*) = *C*.

By doing the folded regression, we halve the number of inferred parameters (*α*_*i*,*j*,*x*,*n*_ or *β*_*i*,*j*,*x*,*n*_). Hence The number of inferred parameters for the folded unconditional regression is 15 × (order + 1), and that of folded conditional regression is 12 × (order + 1).

## Results for multinomial logistic regression

The optimization of the likelihood functions are based on the C++ library of gsl and use the function *gsl_multimin_fminimizer_nmsimplex2* with the initial searching point set to be the results of logistic regression. We here present here 4 figures pertaining to showcase the performance of our model. The regression methods are based on the summary statistic of the counts of mismatches and the optimization is therefore in the scale of miliseconds. ***Figure S1*** and ***Figure S2*** are the conditional regression results of the ancient and control human data correspondingly. And ***Figure S3*** and ***Figure S4*** are the folded conditional regression results of the same data as above.

**Appendix 2–figure S1.**
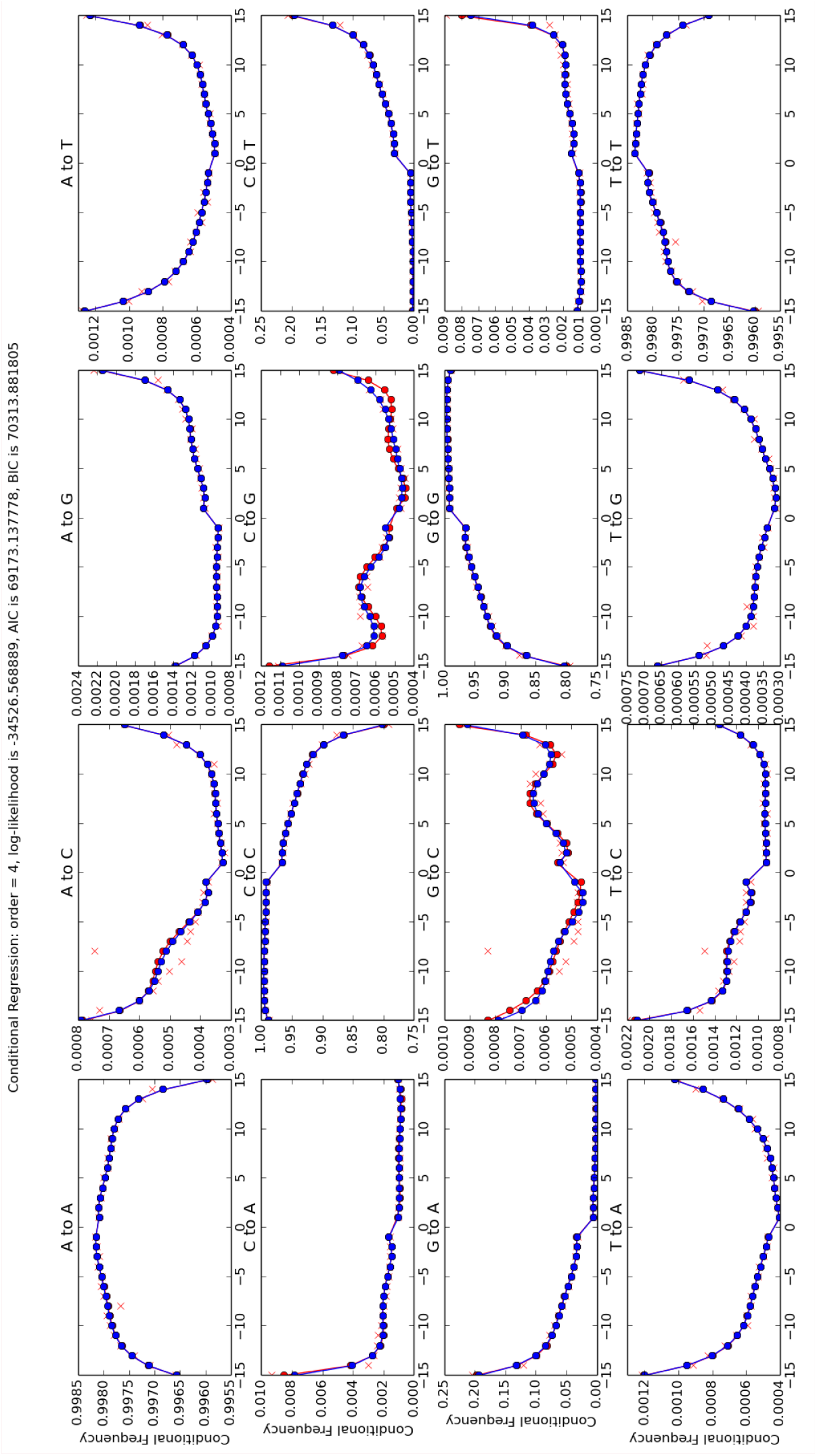
Conditional regression results with the order 4 of the ancient human data. Each panel of figure represents a specific reference/read pair and plots its frequency across different positions. The positions from left to right are −1 to −15 and 15 to 1.

**Appendix 2–figure S2.**
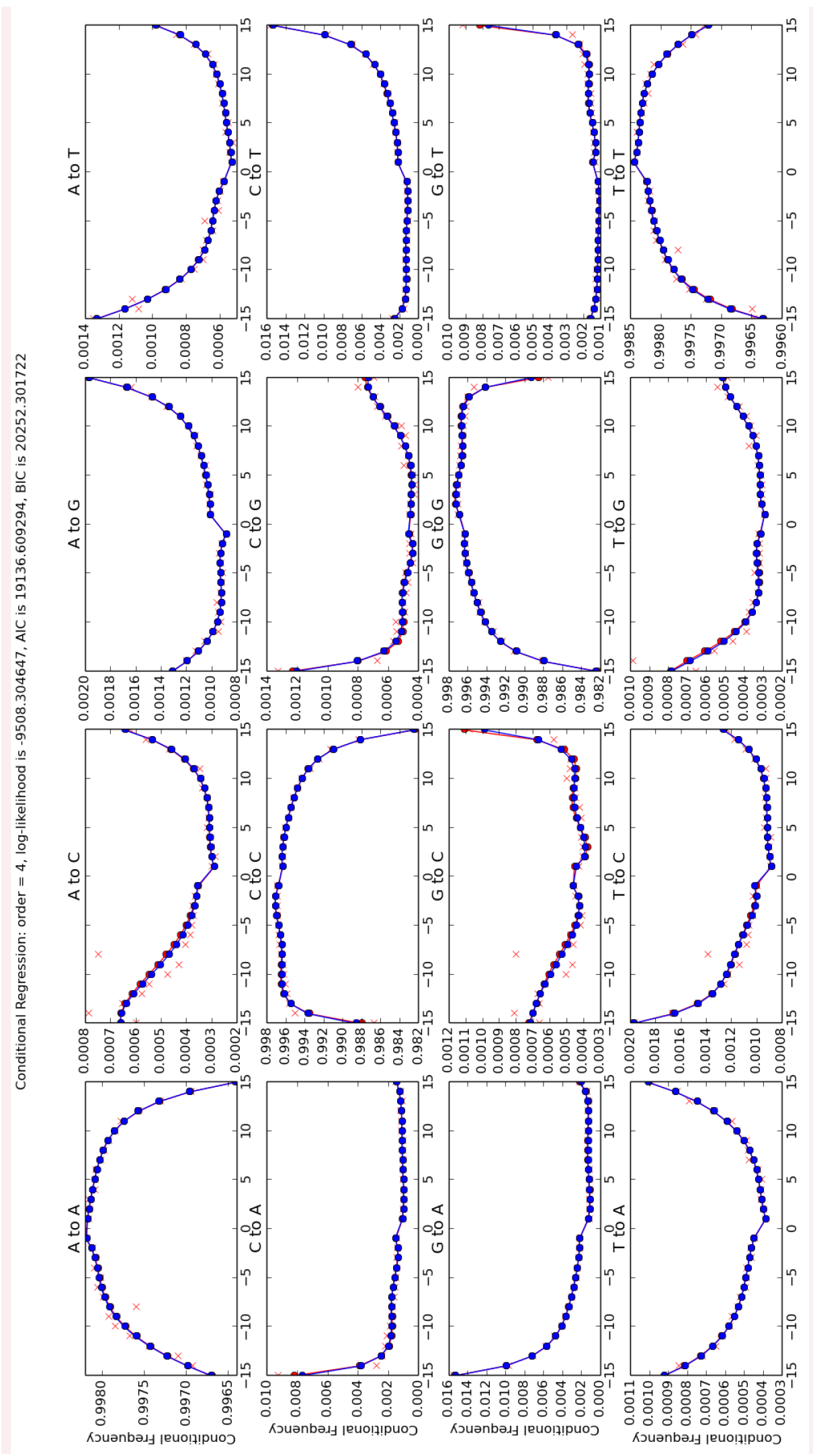
Conditional regression results with the order 4 of the control human data. Each panel of figure represents a specific reference/read pair and plots its frequency across different positions. The positions from left to right are −1 to −15 and 15 to 1.

**Appendix 2–figure S3.**
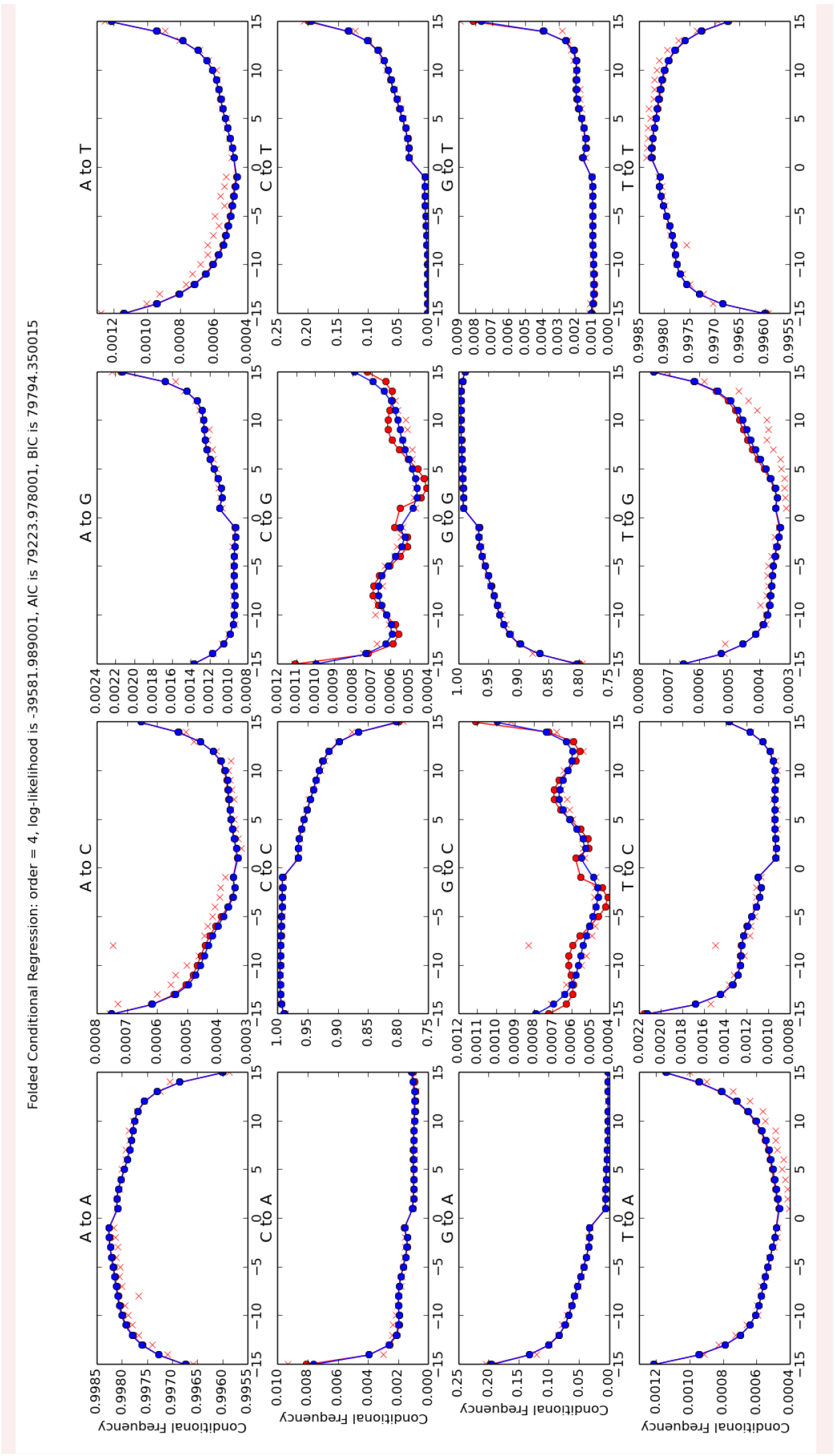
Folded conditional regression results with the order 4 of the ancient human data. Each panel of figure represents a specific reference/read pair and plots its frequency across different positions. The positions from left to right are −1 to −15 and 15 to 1.

**Appendix 2–figure S4.**
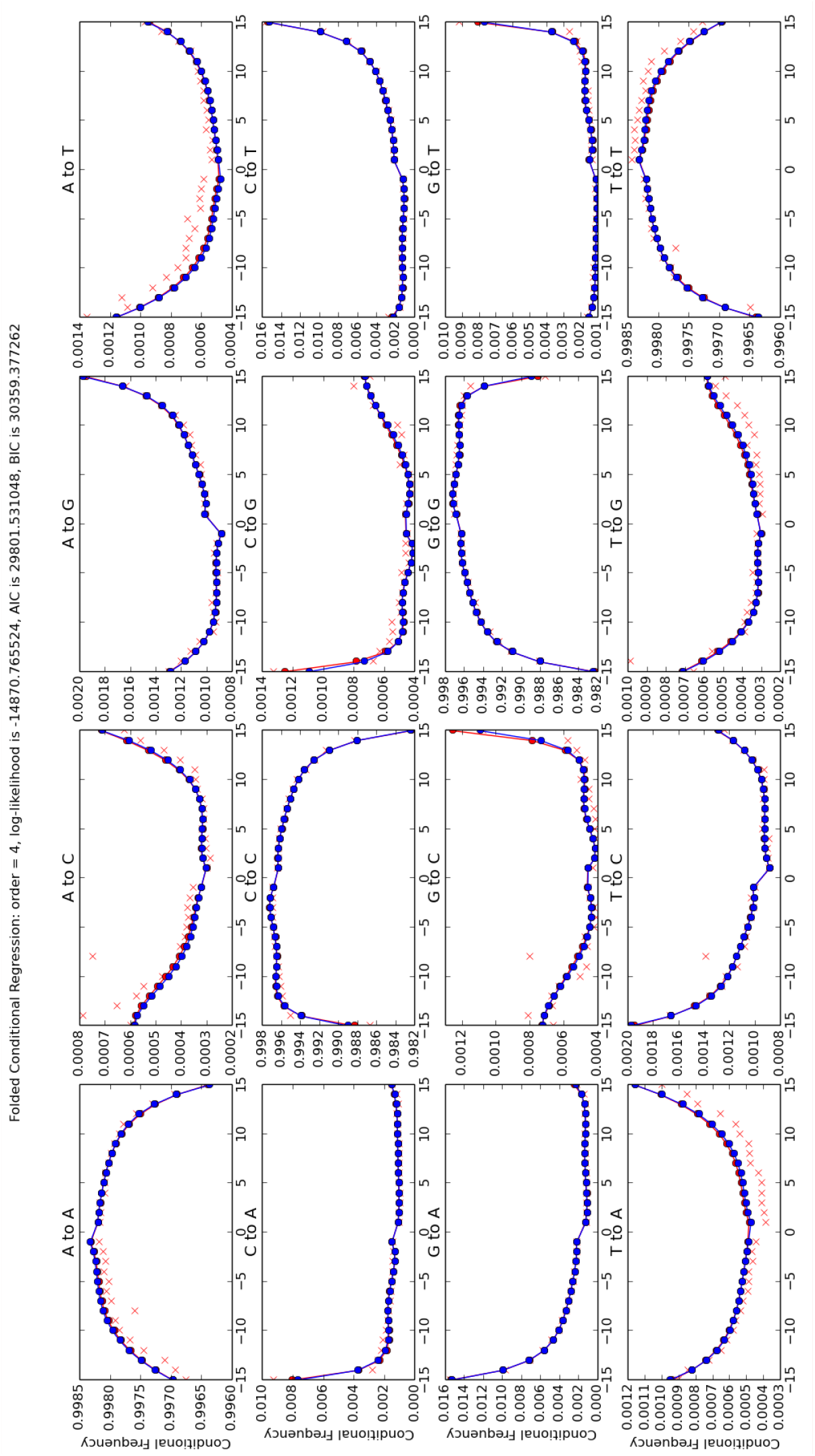
Folded conditional regression results with the order 4 of the control human data. Each panel of figure represents a specific reference/read pair and plots its frequency across different positions. The positions from left to right are −1 to −15 and 15 to 1.

As shown in the figures, the regression models stabilize the coarse mismatch matrices and describe a much more detailed PMD pattern (not only C→T and G→A, but also all other reference and read combinations), but they might suffer from an overfitting issue especially when the data is limited, while the simpler regression model in the main text (***subsection 2.4***) shows an acceptable statistic power even with extremely small amount of data, we thus recommend the readers to use the simpler regression model unless used with extremely high-coverage data.

Our code can also perform the unconditional regression, but as the unconditional regression needs to estimate more parameters based on the same dataset, it is more vulnerable to a possible overfitting issue. We thus only present the figures of the conditional results.

## Appendix 3 NGSNGS COMMANDS

The resulting read data files (fastq files) were simulated with NGSNGS using the above mentioned simulation parameters, all with the same quality scores profiles as used in ART (Huang et al., 2012), based on the Illumina HiSeq 2500 (150 bp). The mapping was performed using Bowtie-2 (Langmead and Salzberg, 2012):

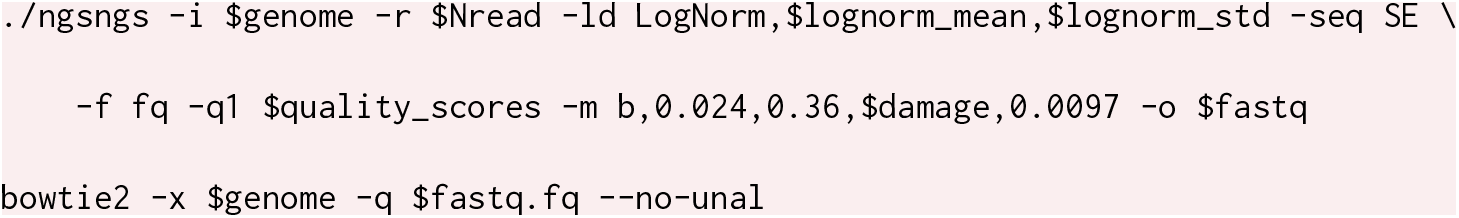

## Appendix 4 NGSNGS SIMULATIONS

The following figures show the `metaDMG` damage estimates for the different NGSNGS simulations (Henriksen, Zhao, and T. Korneliussen, 2022). These simulations include different species (Homo Sapiens and Betula), different GC-levels (low, middle, high), different fragment length distributions (with mean 35, 60, and 90), and different contig lengths (length 1.000, 10.000, 100.000), see ***subsection 3.1*** for more information.

**Appendix 4–figure S5.**
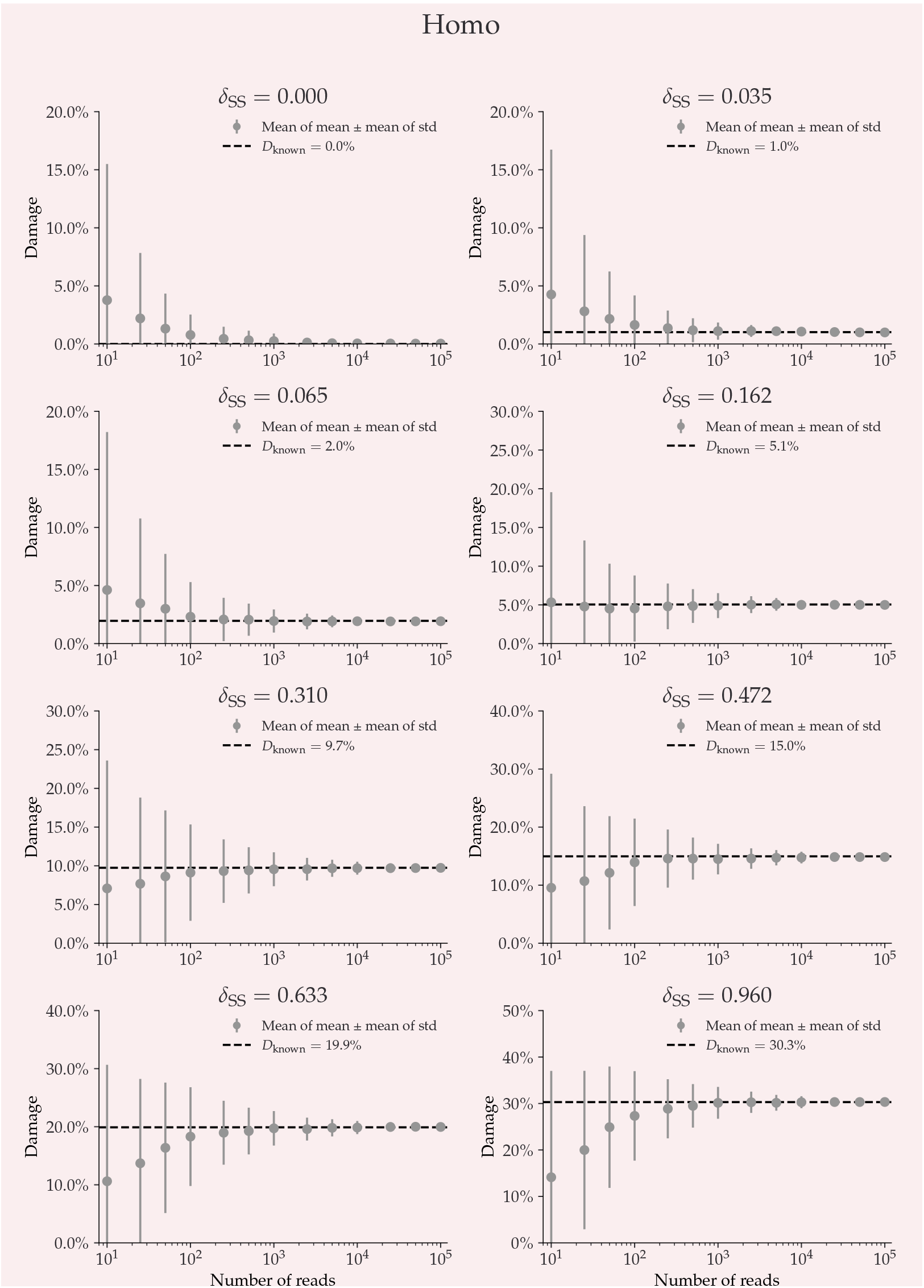
This plot shows the average damage as a function of the number of reads. The grey points show the average of the individual means (with the average of the standard deviations as errors.

**Appendix 4–figure S6.**
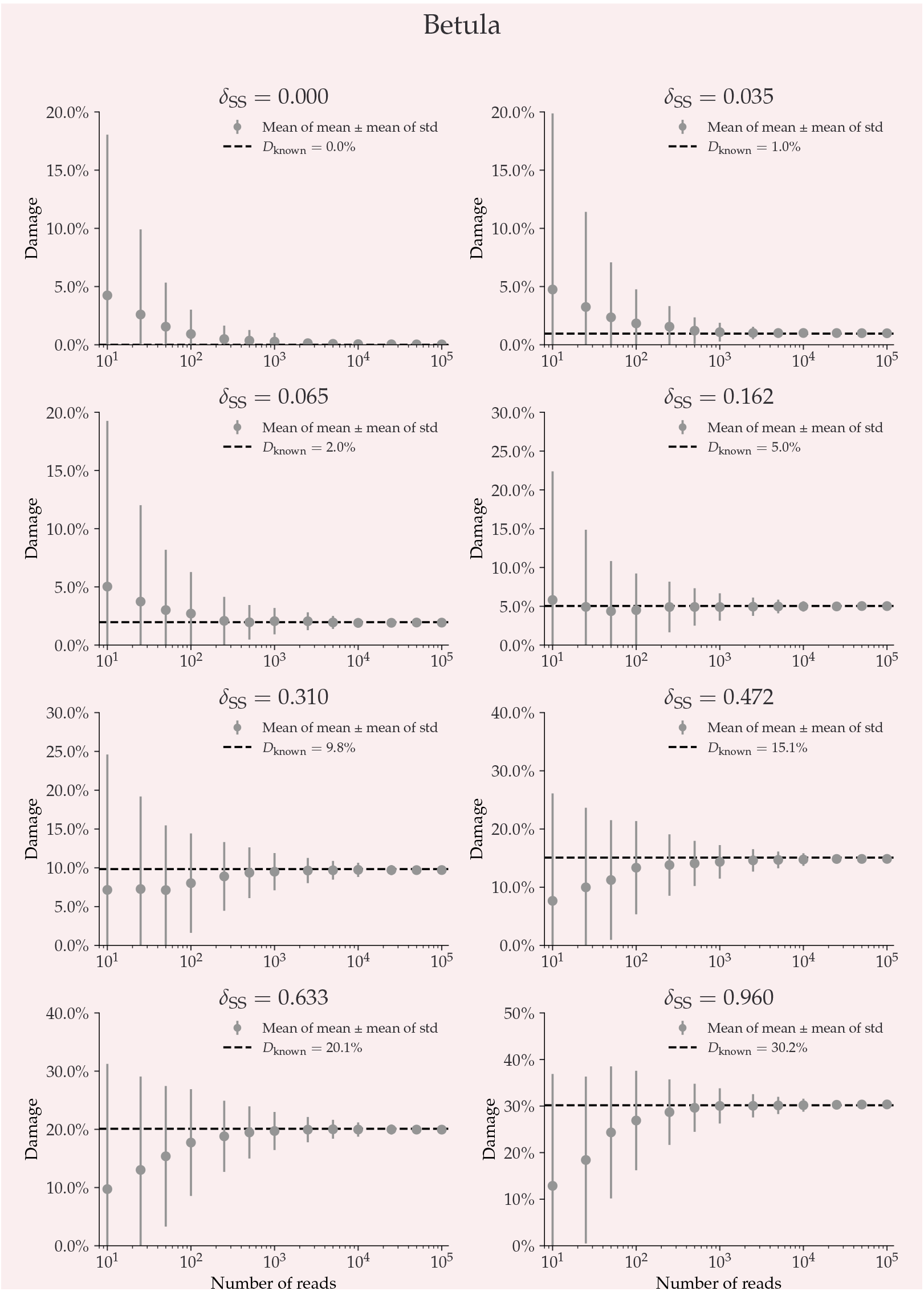
This plot shows the average damage as a function of the number of reads. The grey points show the average of the individual means (with the average of the standard deviations as errors.

**Appendix 4–figure S7.**
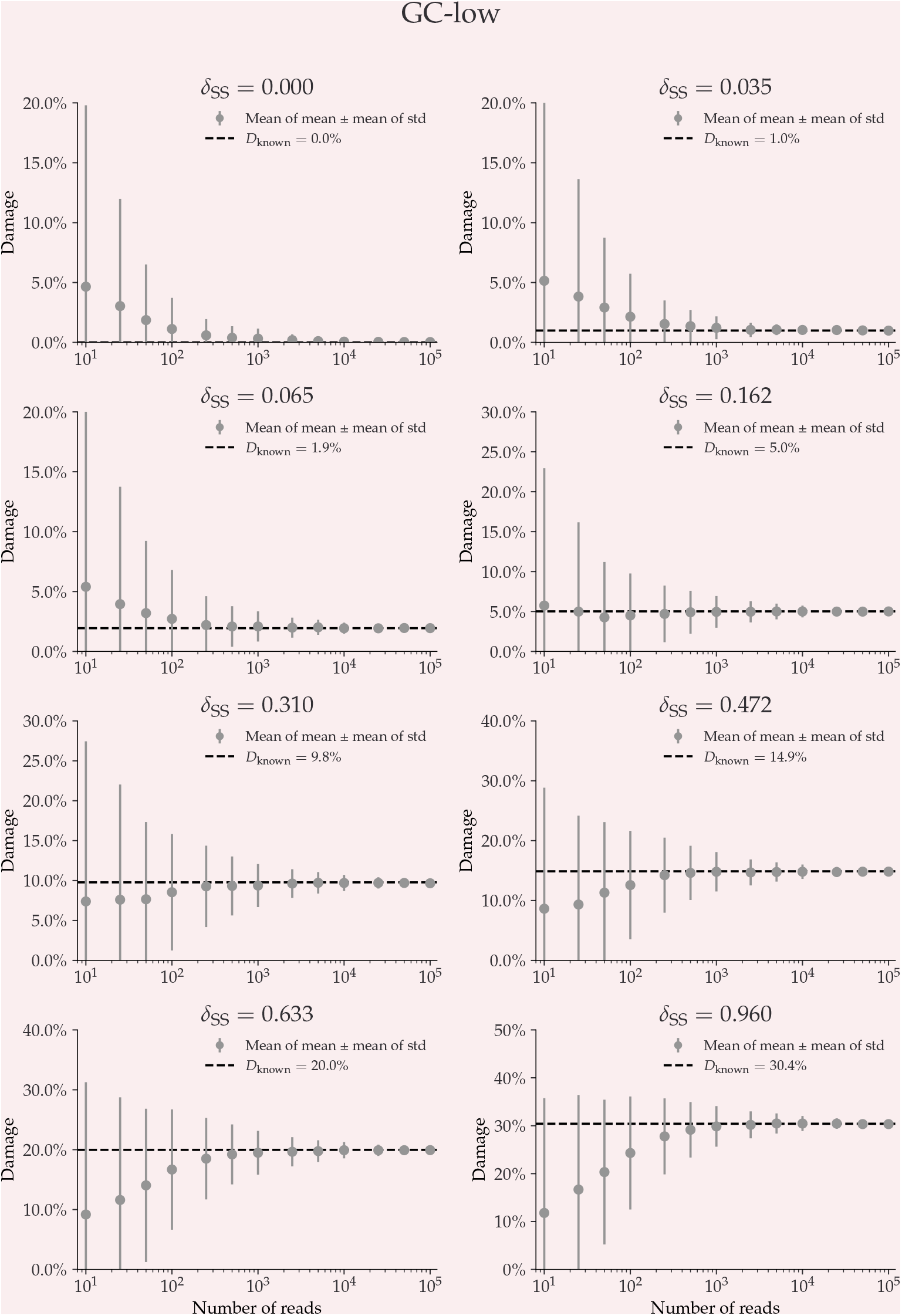
This plot shows the average damage as a function of the number of reads. The grey points show the average of the individual means (with the average of the standard deviations as errors.

**Appendix 4–figure S8.**
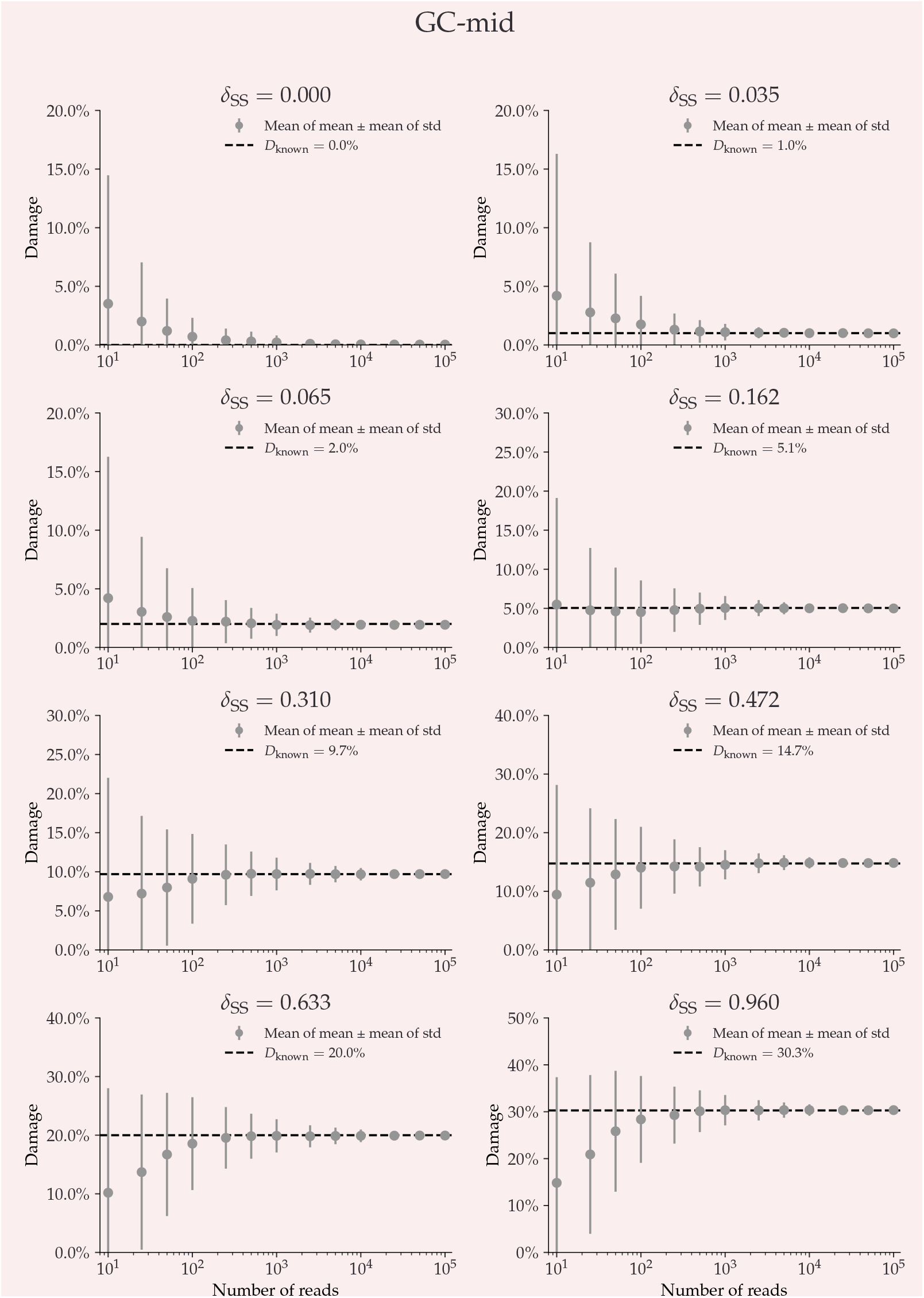
This plot shows the average damage as a function of the number of reads. The grey points show the average of the individual means (with the average of the standard deviations as errors.

**Appendix 4–figure S9.**
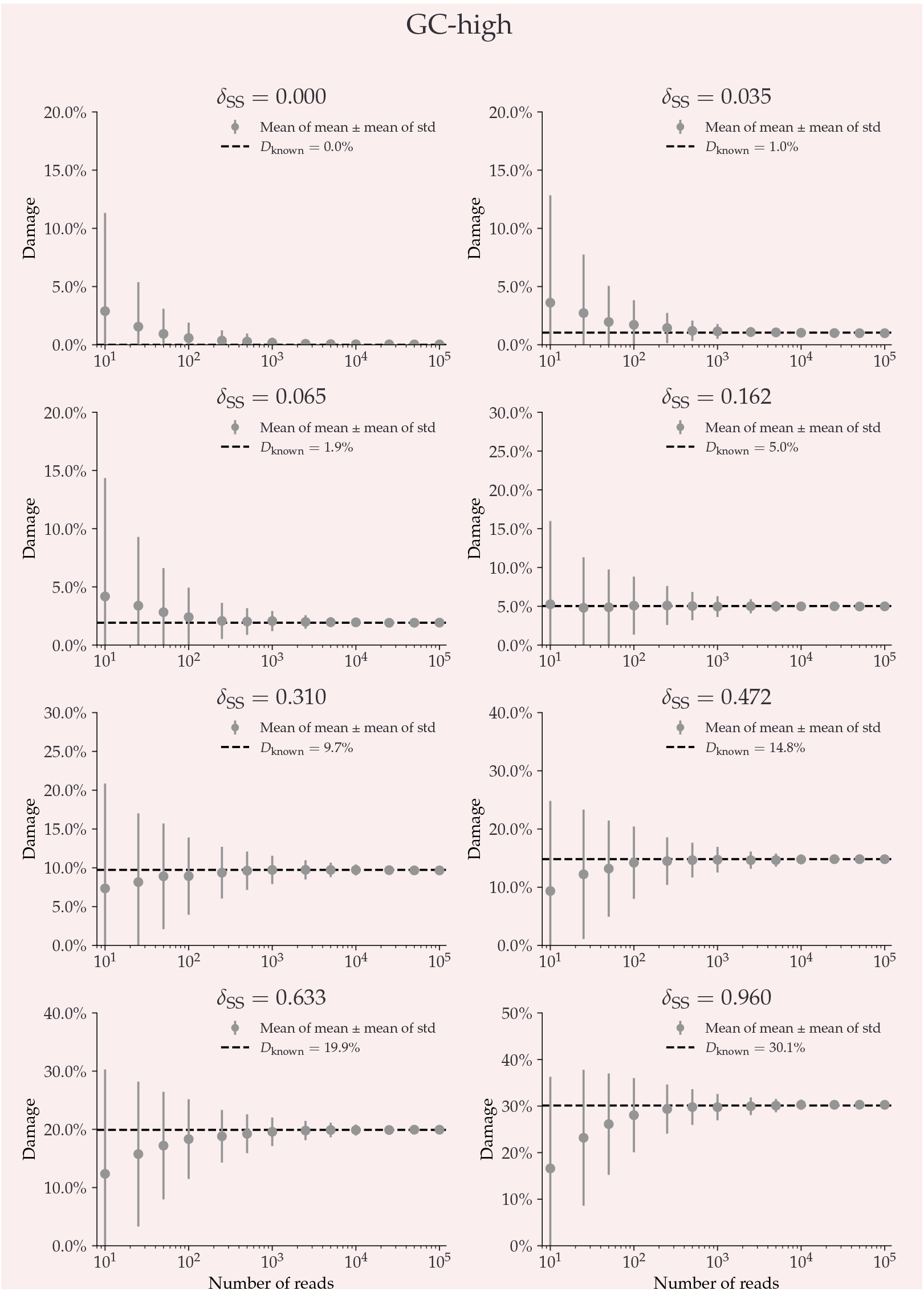
This plot shows the average damage as a function of the number of reads. The grey points show the average of the individual means (with the average of the standard deviations as errors.

**Appendix 4–figure S10.**
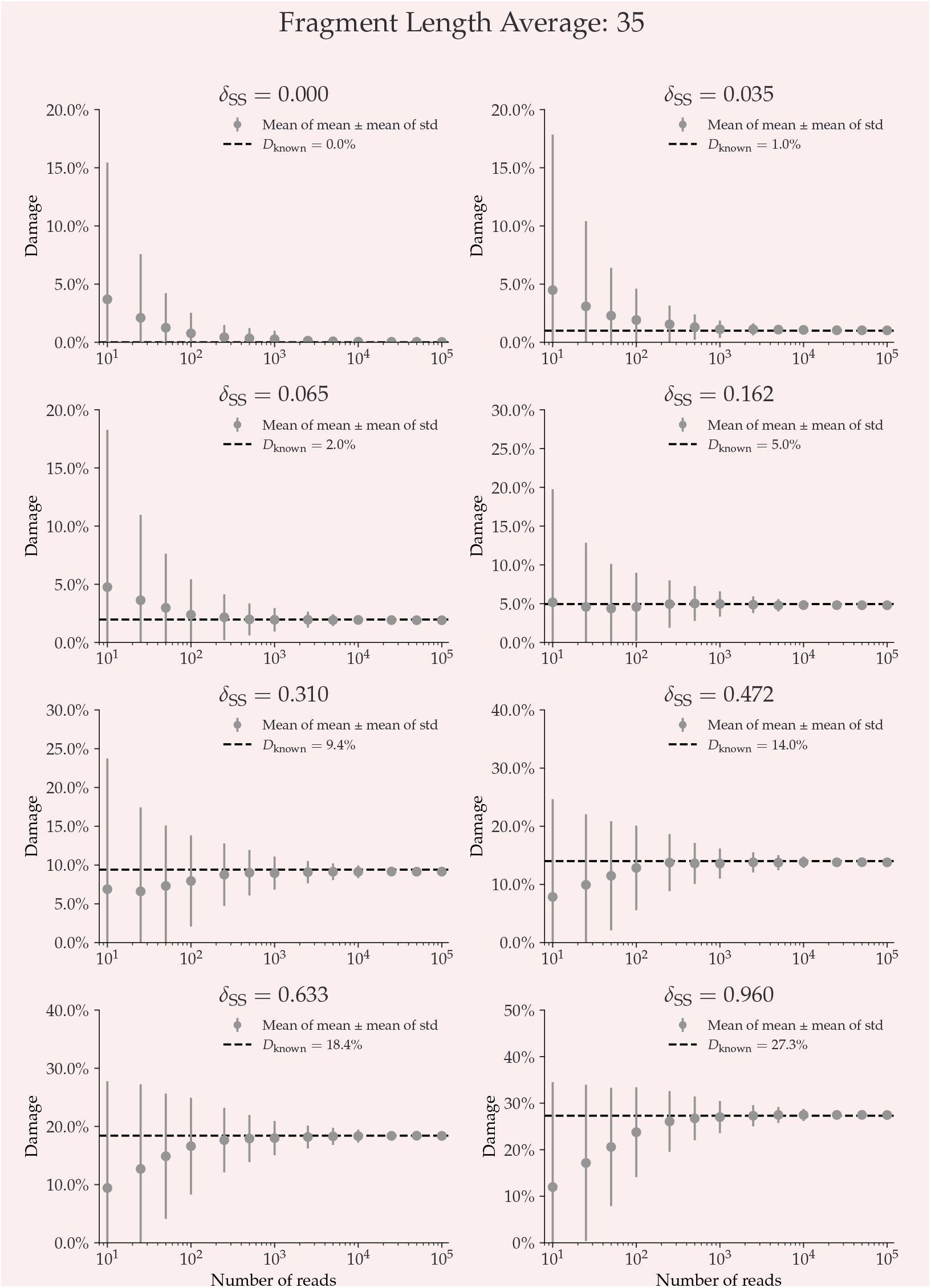
This plot shows the average damage as a function of the number of reads. The grey points show the average of the individual means (with the average of the standard deviations as errors.

**Appendix 4–figure S11.**
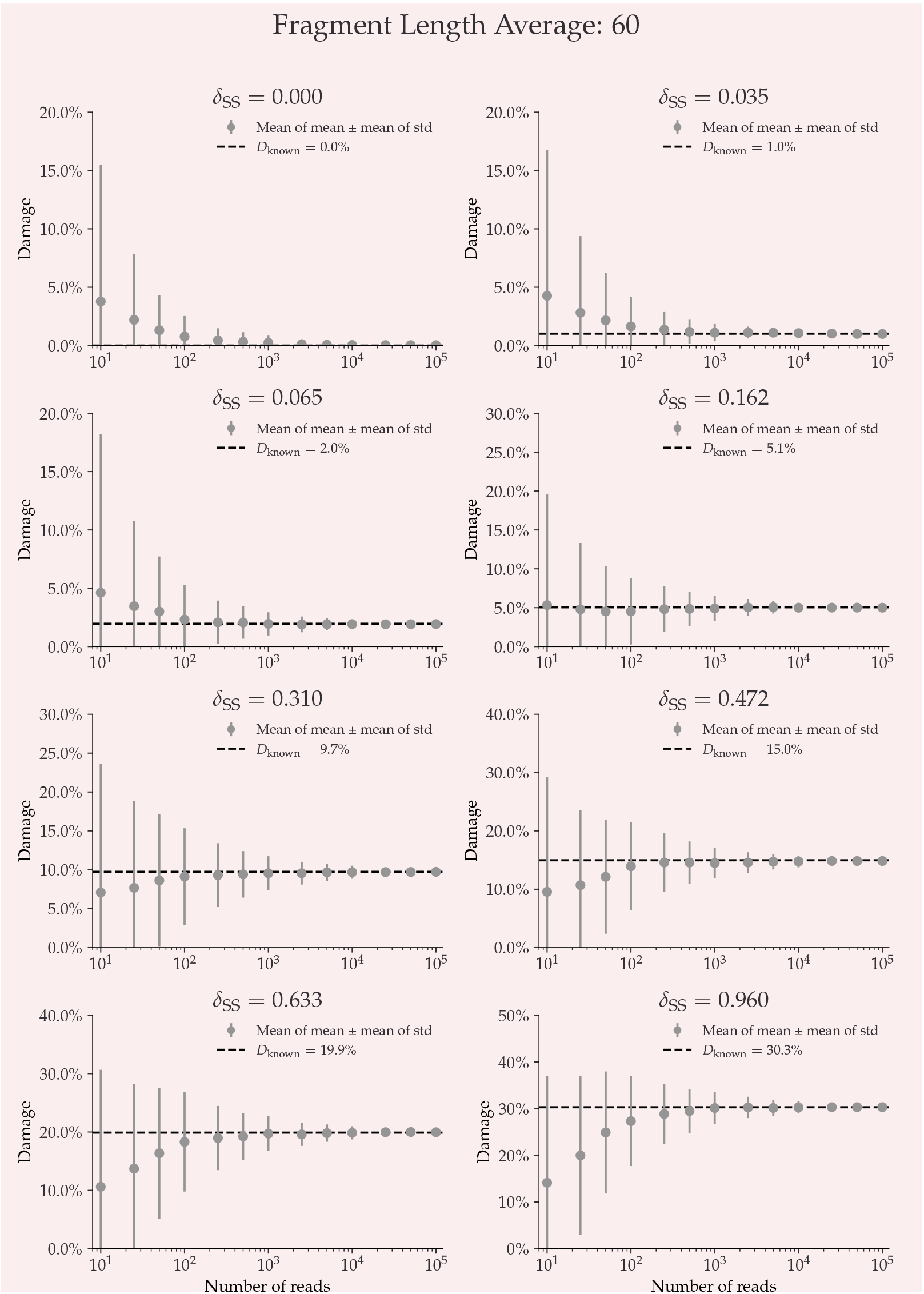
This plot shows the average damage as a function of the number of reads. The grey points show the average of the individual means (with the average of the standard deviations as errors.

**Appendix 4–figure S12.**
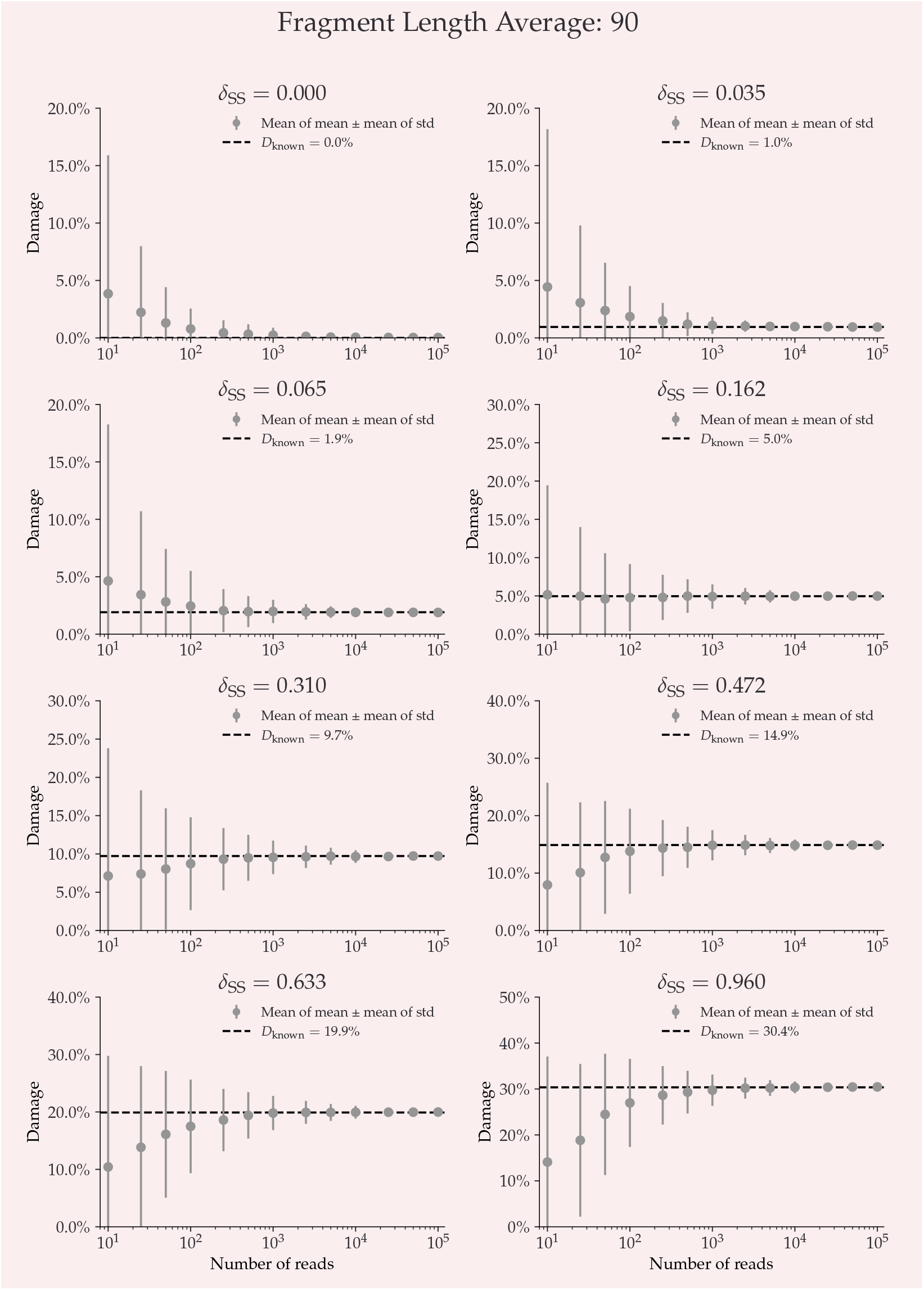
This plot shows the average damage as a function of the number of reads. The grey points show the average of the individual means (with the average of the standard deviations as errors.

**Appendix 4–figure S13.**
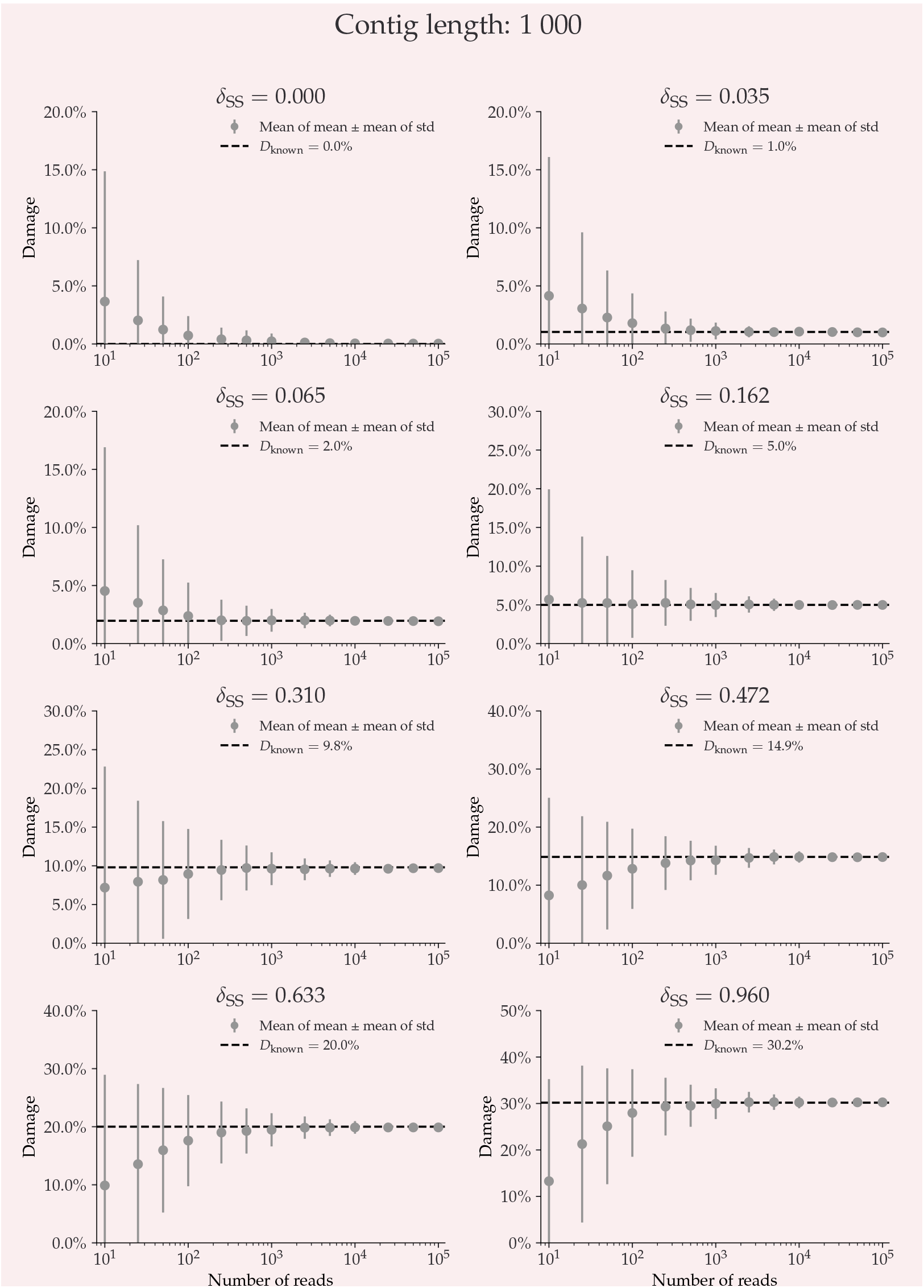
This plot shows the average damage as a function of the number of reads. The grey points show the average of the individual means (with the average of the standard deviations as errors.

**Appendix 4–figure S14.**
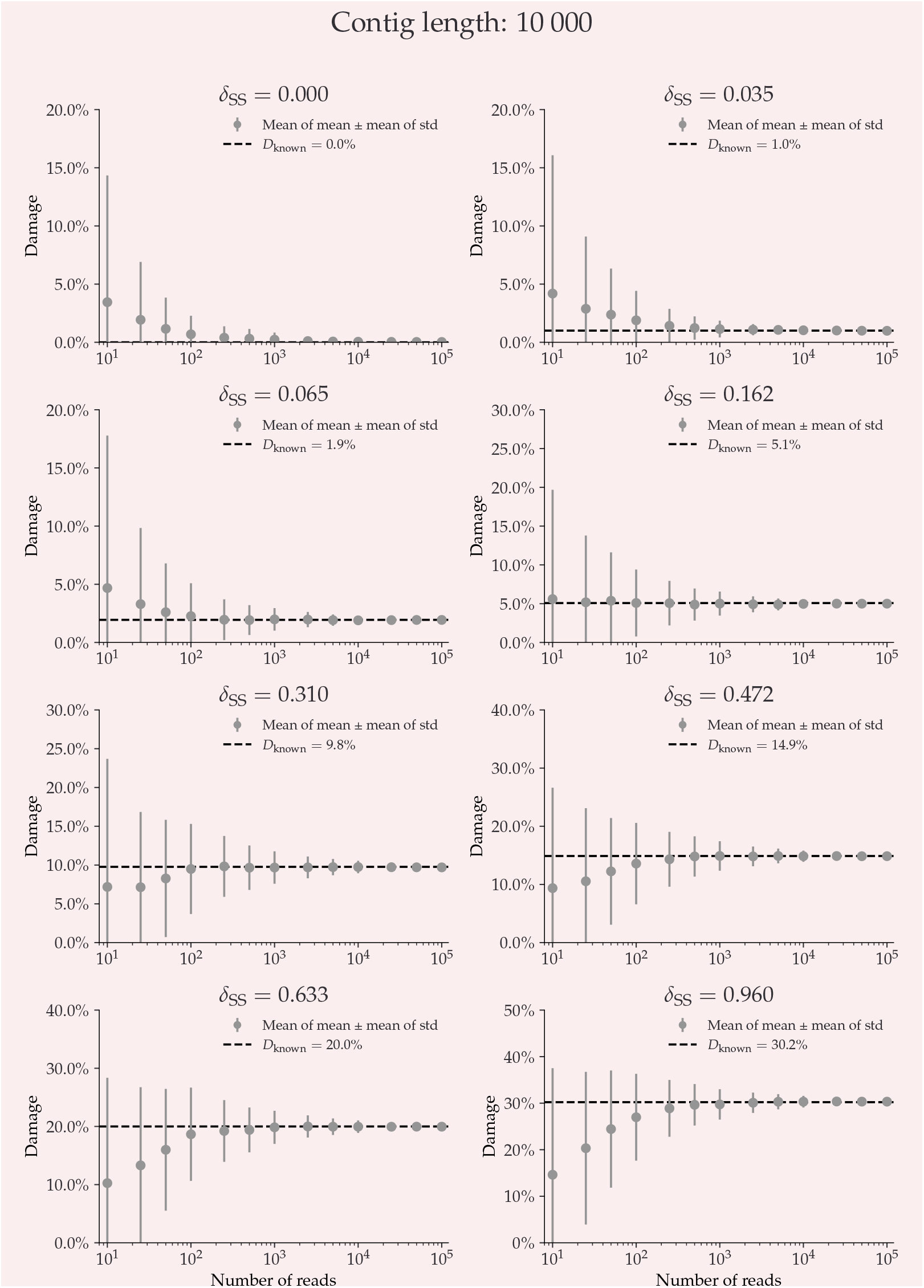
This plot shows the average damage as a function of the number of reads. The grey points show the average of the individual means (with the average of the standard deviations as errors.

**Appendix 4–figure S15.**
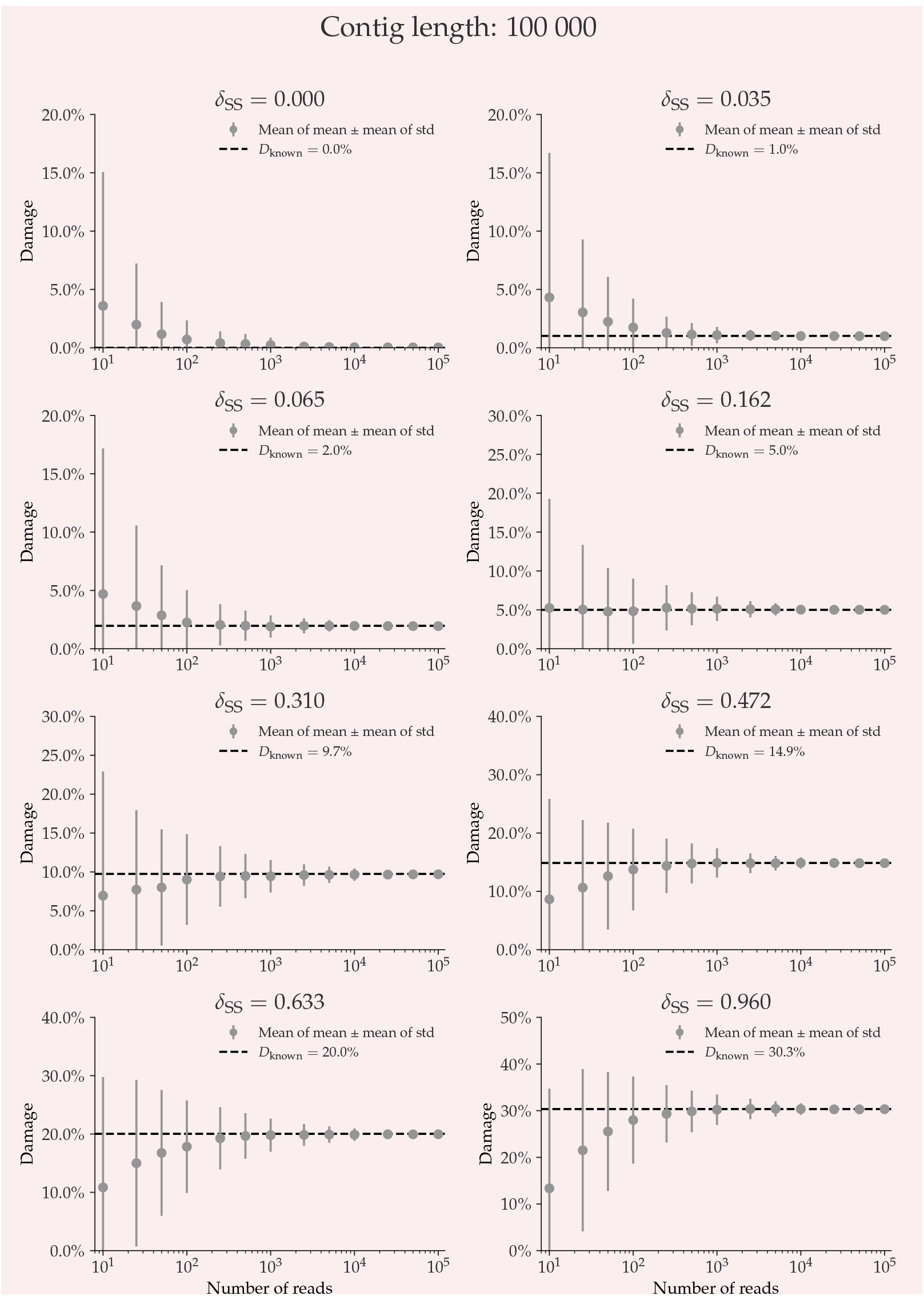
This plot shows the average damage as a function of the number of reads. The grey points show the average of the individual means (with the average of the standard deviations as errors.

## Appendix 5 NGSNGS SIMULATIONS – ZERO DAMAGE

Damage estimates for non-damaged simulated data, each with 1000 replications, see ***subsection 3.1***. The inferred damage is shown on the y-axis and the significance on the x-axis. Each simulation is shown as a single cross and the red lines show the kernel density estimate (KDE) of the damage estimates. The marginal distributions are shown as histograms next to the scatter plot.

**Appendix 5–figure S16.**
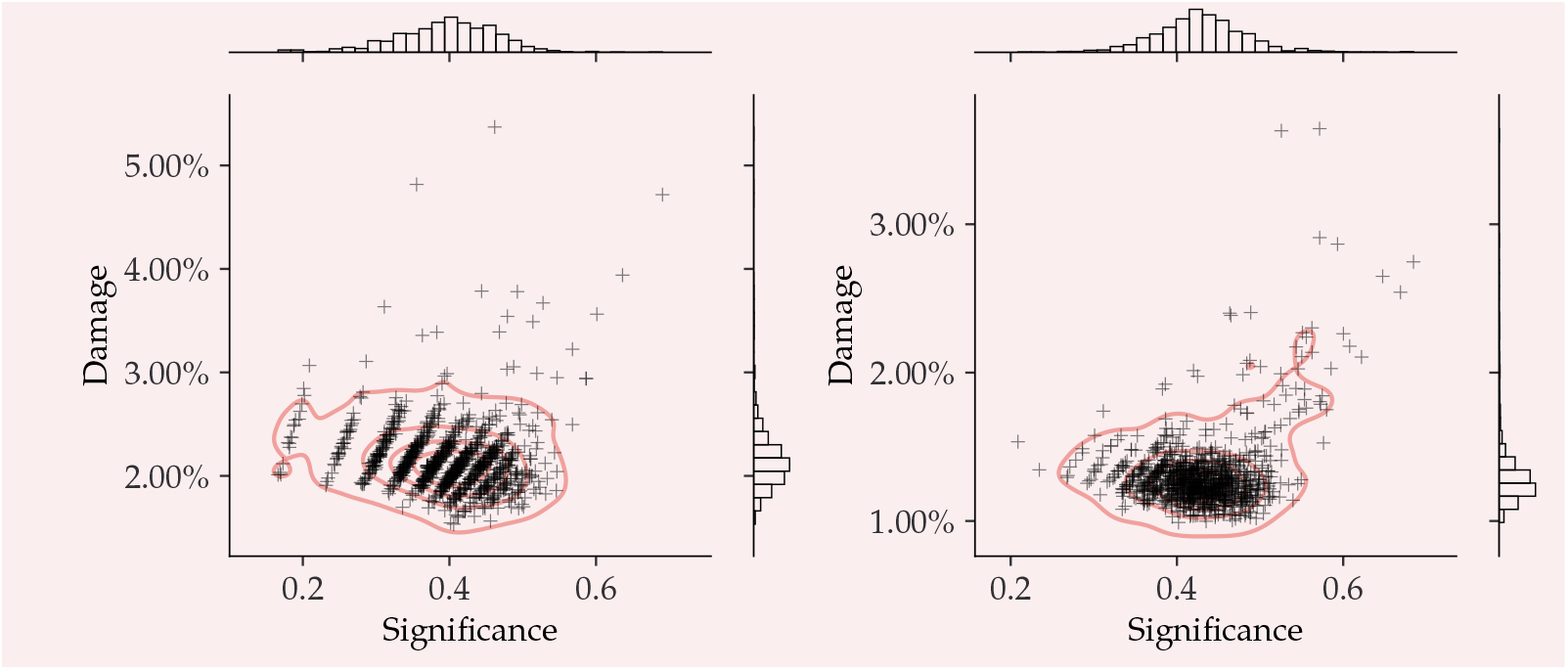
Left) 25 simulated reads. Right) 50 simulated reads.

**Appendix 5–figure S17.**
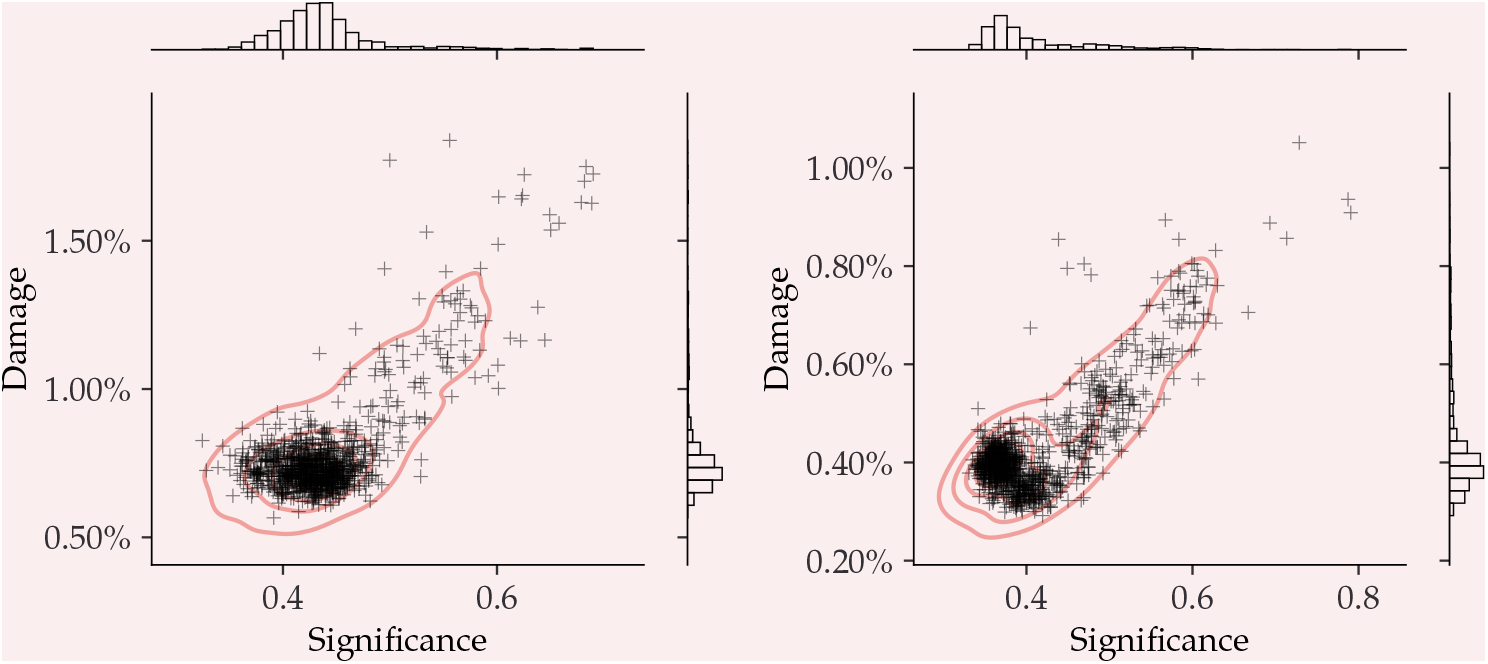
Left) 100 simulated reads. Right) 250 simulated reads.

**Appendix 5–figure S18.**
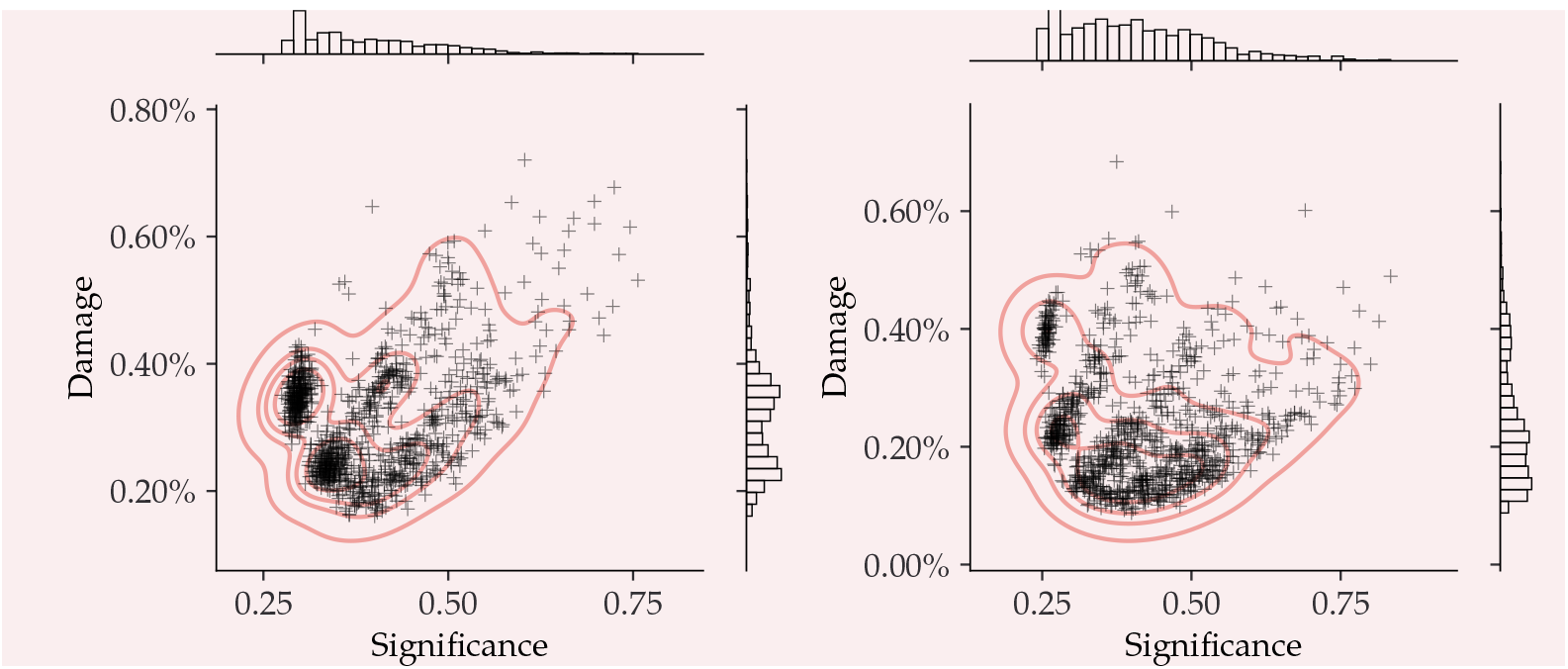
Left) 500 simulated reads. Right) 1.000 simulated reads.

**Appendix 5–figure S19.**
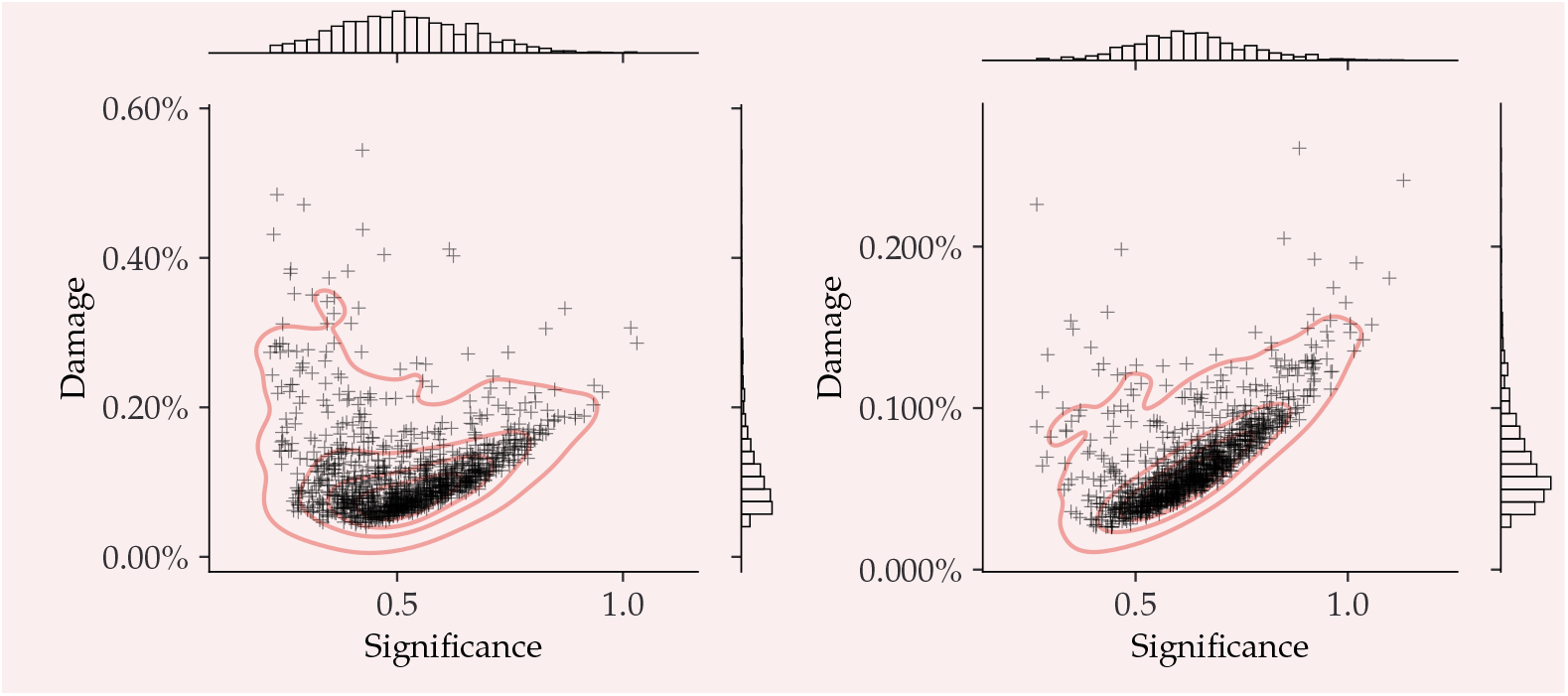
Left) 2.500 simulated reads. Right) 5.000 simulated reads.

**Appendix 5–figure S20.**
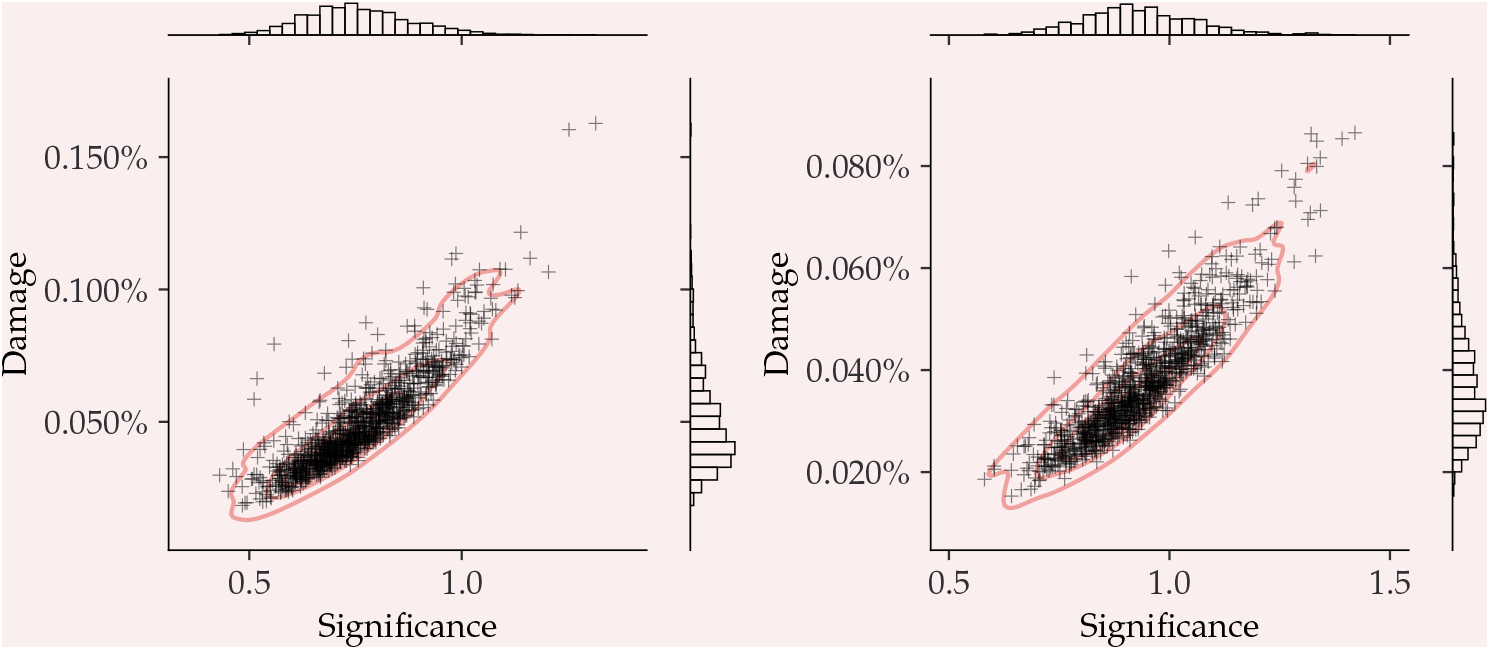
Left) 10.000 simulated reads. Right) 25.000 simulated reads.

**Appendix 5–figure S21.**
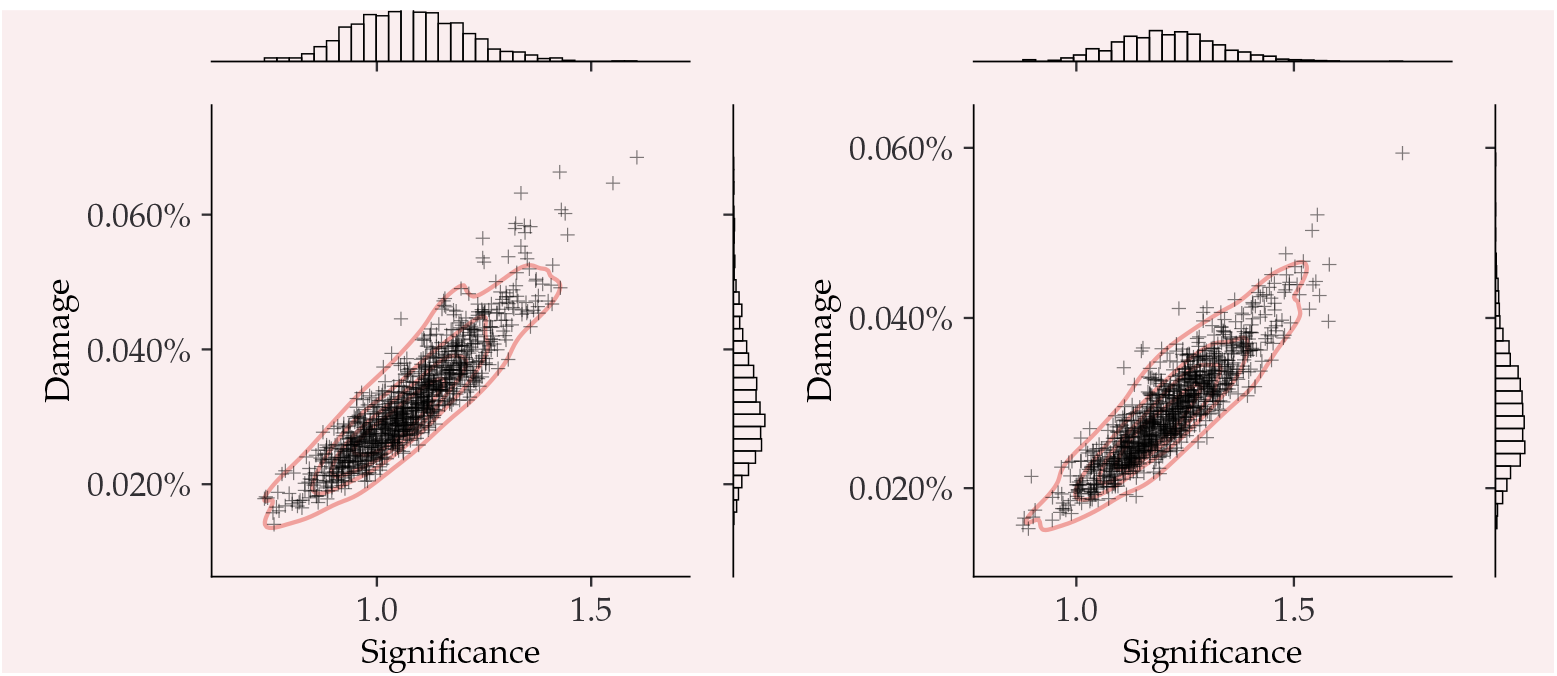
Left) 50.000 simulated reads. Right) 100.000 simulated reads.

## Appendix 6 FALSE NEGATIVES

Even though the simple requirement of having more than 100 reads drastically improves the performance of the damage estimates, see ***subsection 4.2***, it does not identify all of the species that were simulated to be ancient. One of these non-identified taxa is the Stenotrophomonas Maltophilia species in the Pitch-6 sample. We show the damage estimates for different simulations for this particular taxa in ***Figure S22*** to quantify the behaviour of the damage estimate at the different stages of the simulation pipeline. For the final stage in the gargammel pipeline, ie. including fragmentation, deamination, and sequencing noise (red in the figure), only 167 reads are assigned to this specific taxa after mapping, when a total of 1 million reads were simulated. The significance is *Z*_fit_ = 1.9, just below the damage threshold.

**Appendix 6–figure S22.**
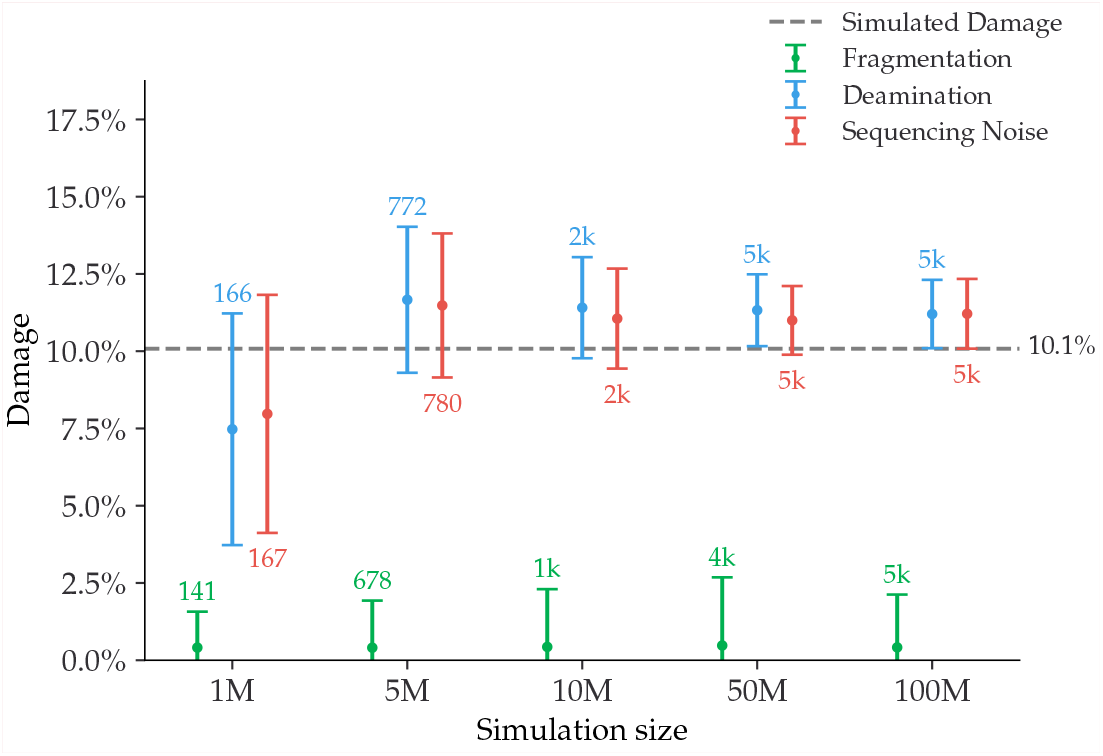
Damage estimates of the Stenotrophomonas maltophilia species from the Pitch-6 sample. Damage is shown as a function of the total simulation size, with the fragmentation files in green, the deamination files in blue and the final files including sequencing errors in red. All errors are 1*σ* error bars (standard deviation). The number of reads for each fit is shown as text the simulated amount of damage is shown as a dashed grey line.

## Appendix 7 BAYES VS. MAP

**Appendix 7–figure S23.**
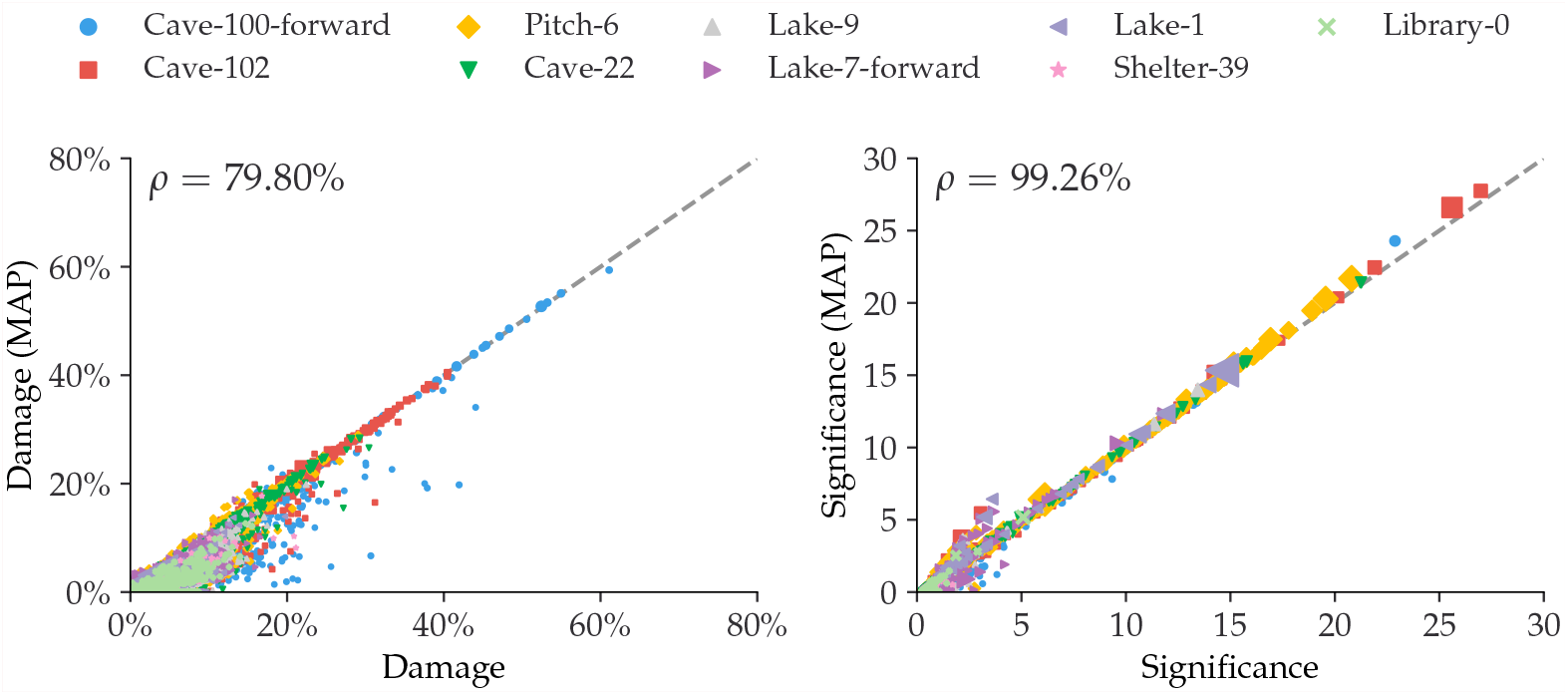
Comparison between the full Bayesian model and the fast, approximate, MAP model for the estimated damage and significance. The dashed, grey line shows the 1:1 ratio.

## Appendix 8 PYDAMAGE COMPARISON

The following figures show the parallel coordinates plot comparing `metaDMG` and PyDamage for the Homo Sapiens single-genome simulation with 100 reads for different amount of artificially added damage, see ***subsection 4.5***. The two first axes show the estimated damage: *D*_fit_ by `metaDMG` and *p*_max_ by PyDamage. The following two axes show the fit quality: significance (*Z*_fit_) by `metaDMG` and the predicted accuracy (Acc_pred_) by PyDamage. The final axis shows the *q*-value by PyDamage. Each of the 100 replications are plotted as single lines. Replications passing the relaxed `metaDMG` damage threshold (*D*_fit_ > 1% and *Z*_fit_ > 2) are shown in color proportional to their significance. Replications that did not pass are shown in semi-transparent black lines.

**Appendix 8–figure S24.**
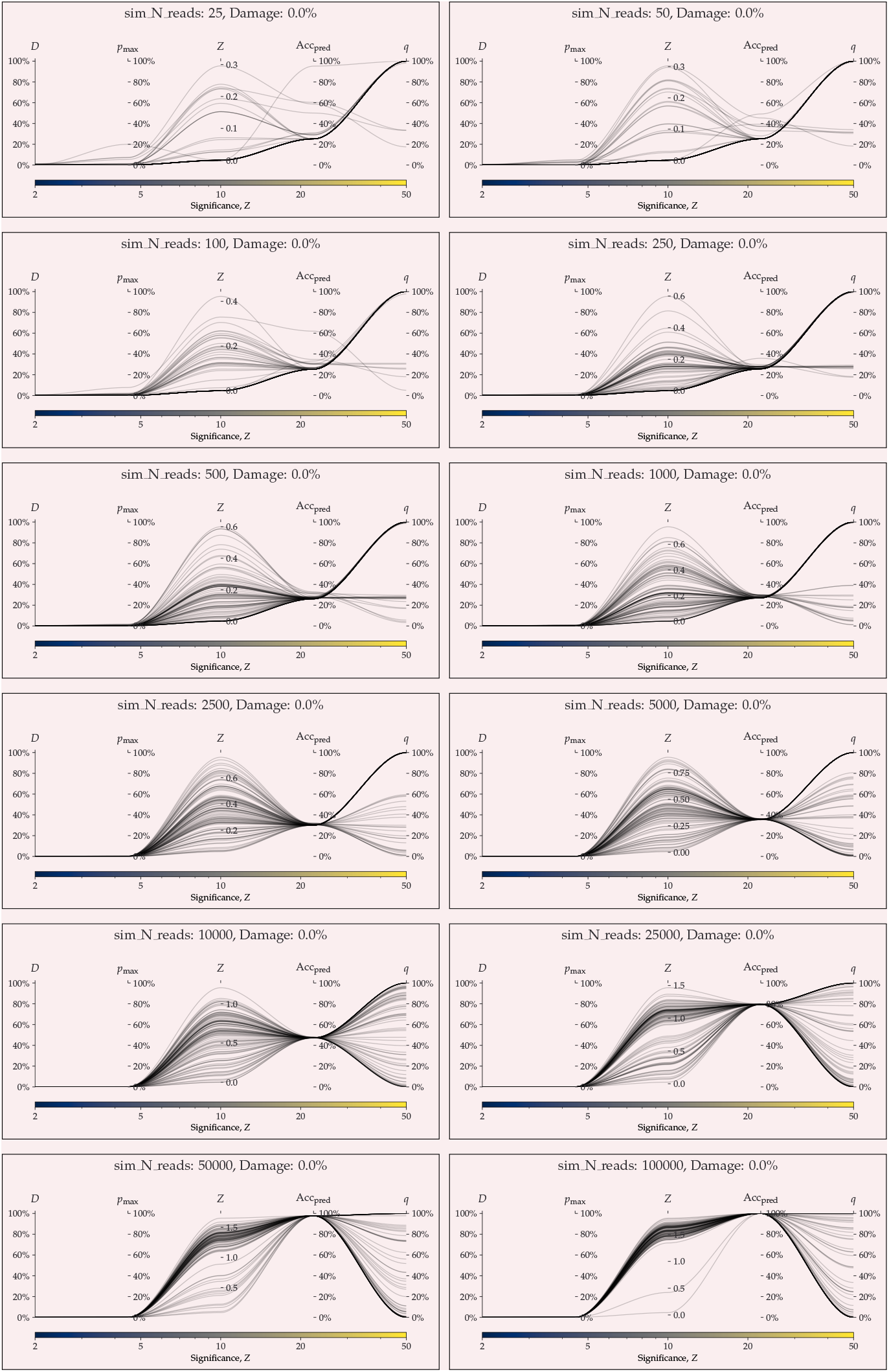
parallel coordinates plot comparing `metaDMG` and PyDamage for 0% artificial damage.

**Appendix 8–figure S25.**
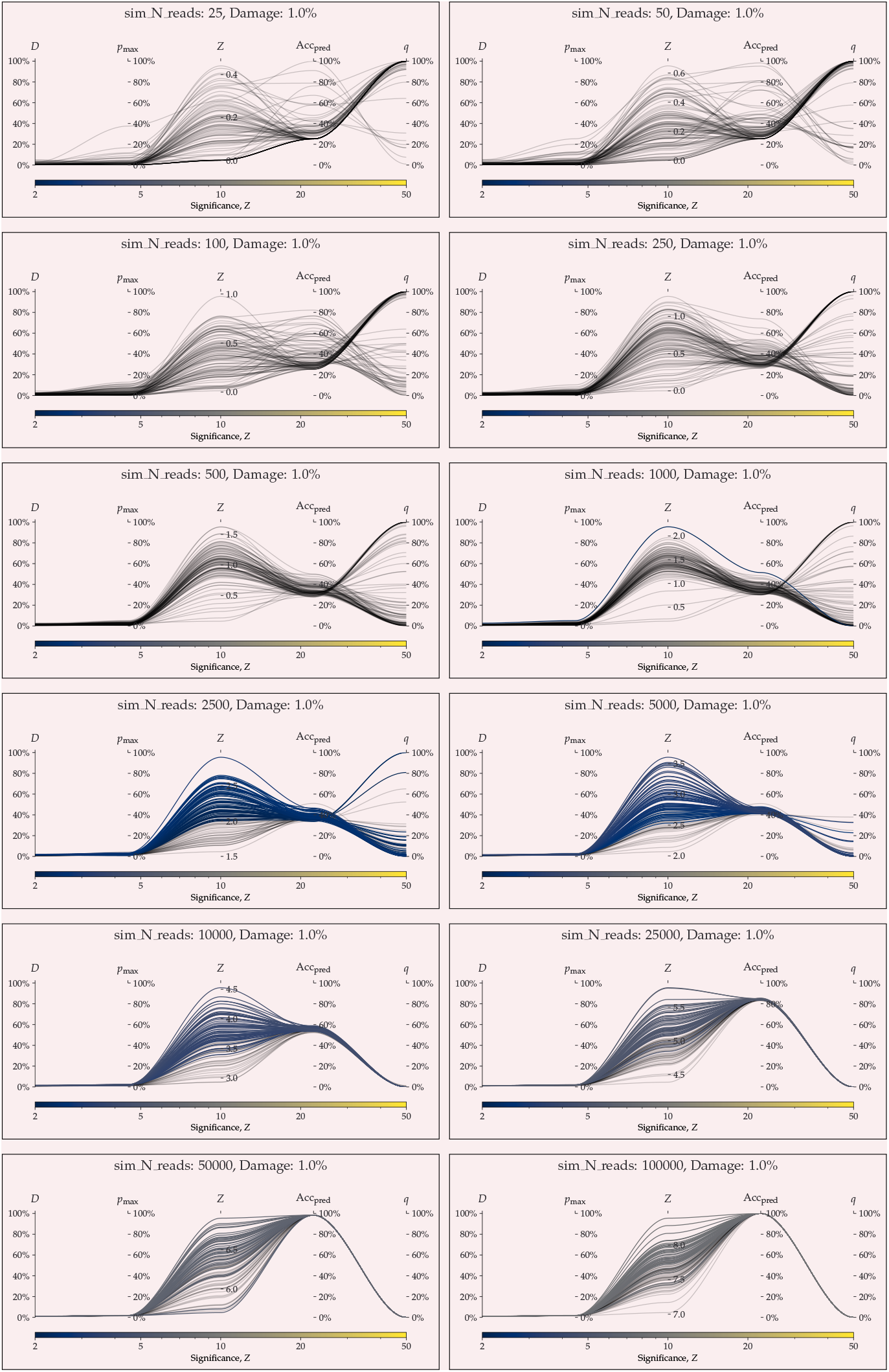
parallel coordinates plot comparing `metaDMG` and PyDamage for 1% artificial damage.

**Appendix 8–figure S26.**
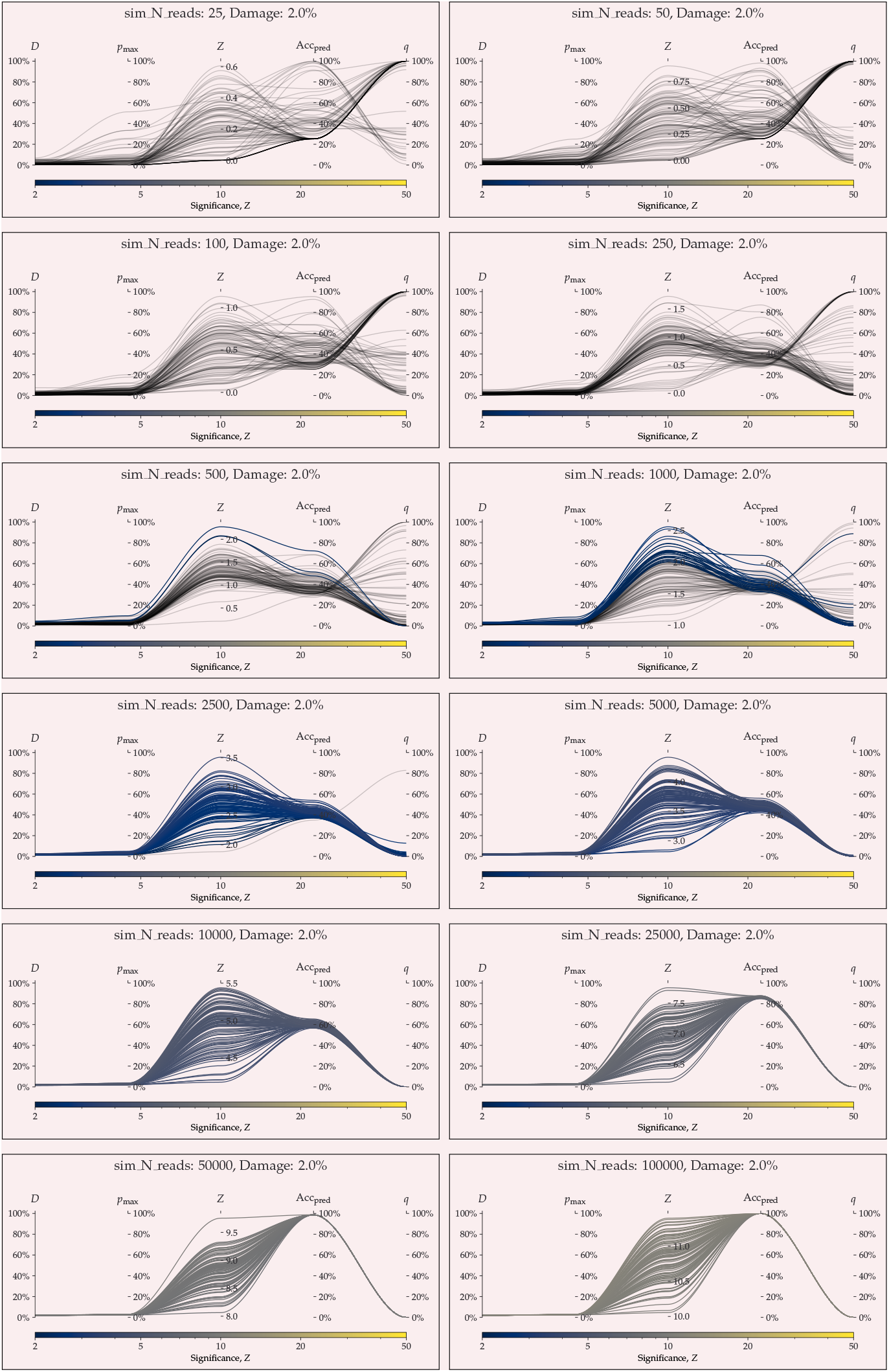
parallel coordinates plot comparing `metaDMG` and PyDamage for 2% artificial damage.

**Appendix 8–figure S27.**
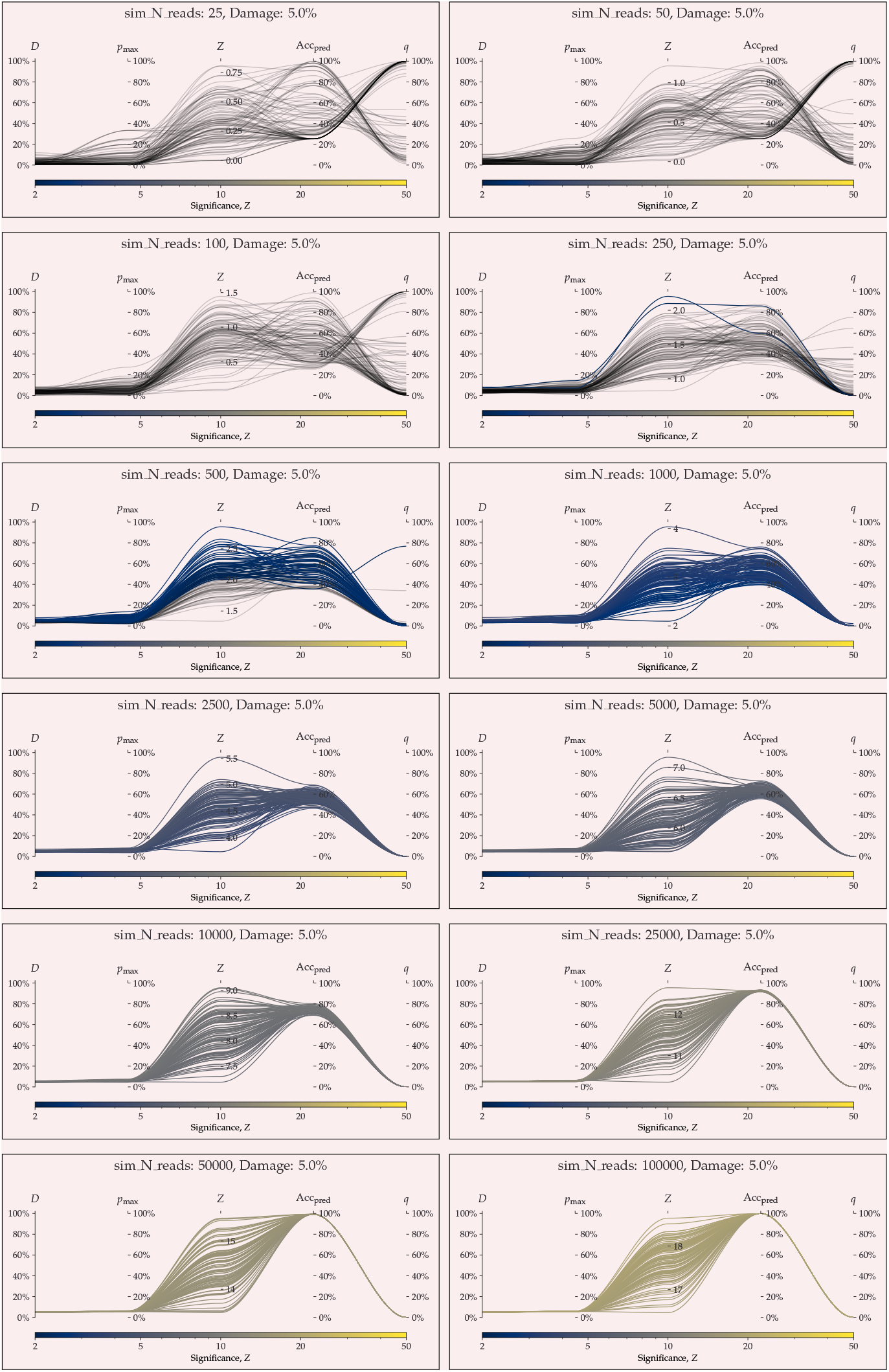
parallel coordinates plot comparing `metaDMG` and PyDamage for 5% artificial damage.

**Appendix 8–figure S28.**
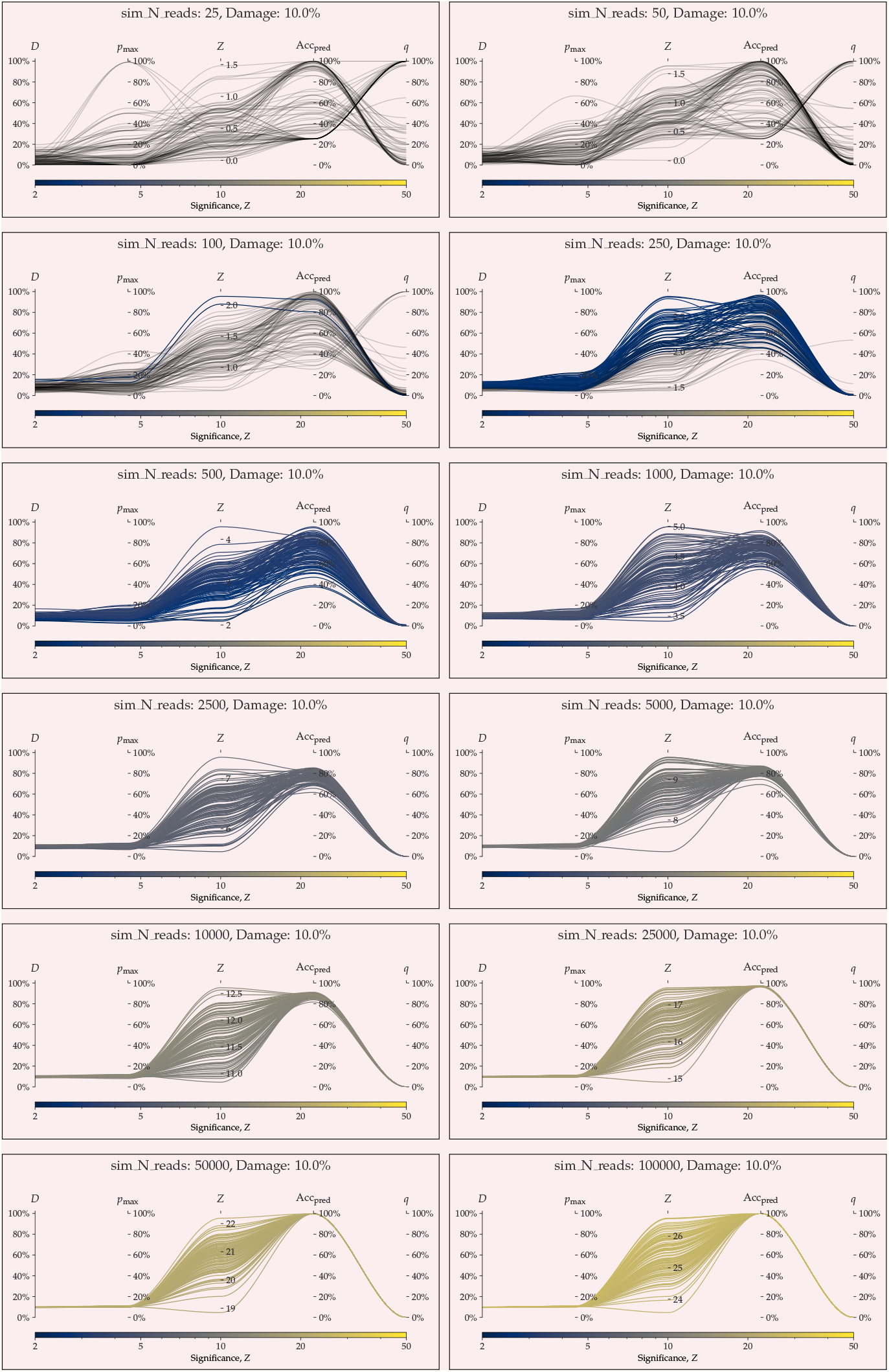
parallel coordinates plot comparing `metaDMG` and PyDamage for 10% artificial damage.

**Appendix 8–figure S29.**
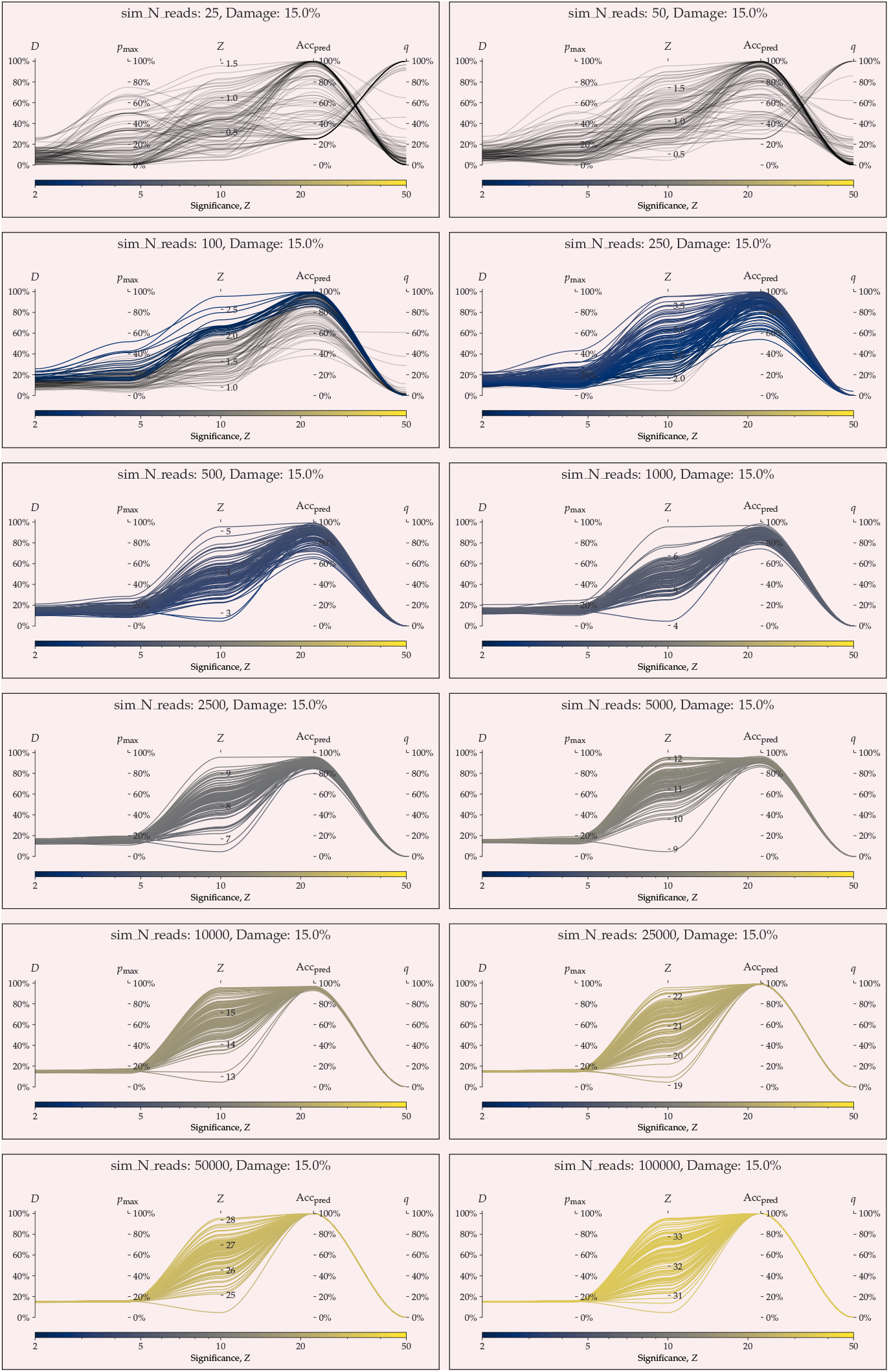
parallel coordinates plot comparing `metaDMG` and PyDamage for 15% artificial damage.

**Appendix 8–figure S30.**
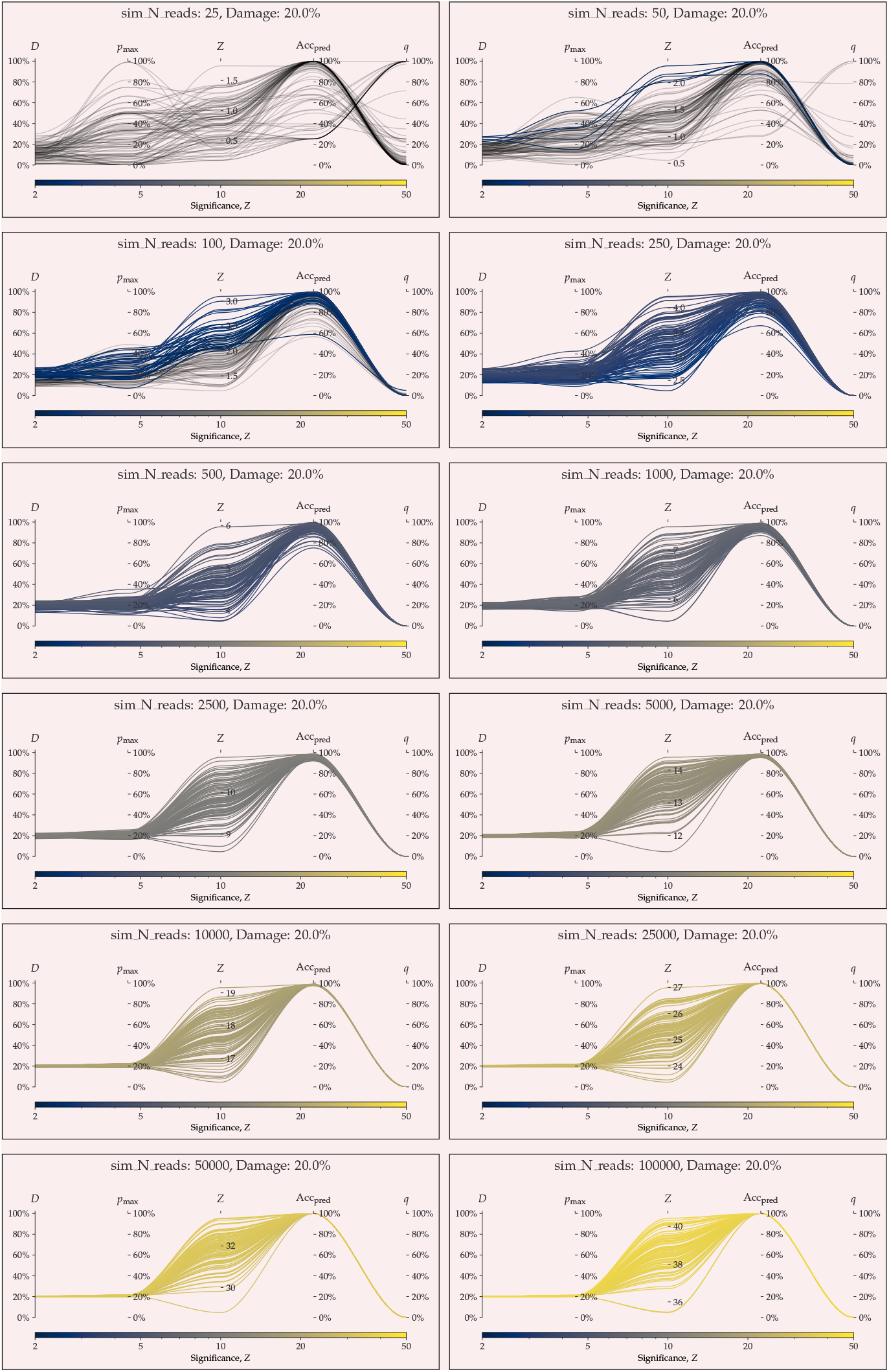
parallel coordinates plot comparing `metaDMG` and PyDamage for 20% artificial damage.

**Appendix 8–figure S31.**
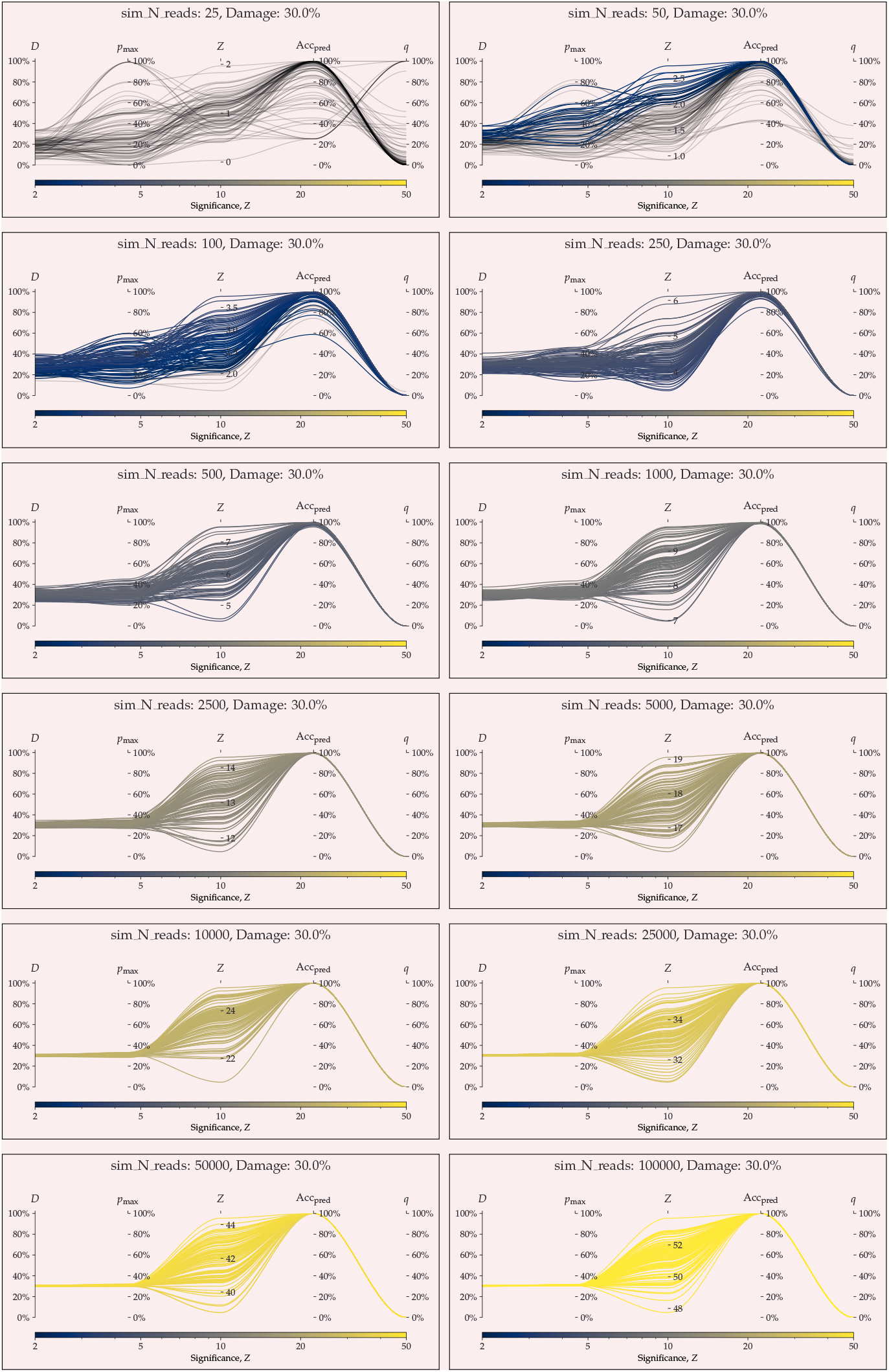
parallel coordinates plot comparing `metaDMG` and PyDamage for 30% artificial damage.

## Footnotes

1 Note that we do not parameterize the beta distribution in terms of the common (*α*, *β*) parameterization, but instead using the more intuitive (*μ*, *ϕ*) parameterization. One can reparameterize (*α*, *β*) → (*μ*, *ϕ*) using the following two equations: 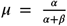 and *ϕ* = *α* + *β* (Cepeda-Cuervo and Cifuentes-Amado, 2017).

2 Parameterized as (*μ*, *ϕ*)

3 NCBI: NC_012920.1

4 NCBI: KX703002.1

5 NCBI: NZ_CP024731.1

6 NCBI: NZ_LS483369.1

7 NCBI: GCA_001929375.1

